# Ribosome transfer via tunnelling nanotubes rescues protein synthesis in pancreatic cancer cells

**DOI:** 10.1101/2024.06.06.597772

**Authors:** Stanislava Martínková, Denisa Jansová, Jana Vorel, Lucie Josefa Lamačová, Petr Daniel, Michael Hudec, Mário Boďo, Jan Hajer, Andrej Susor, Jan Trnka

## Abstract

Graphical abstract

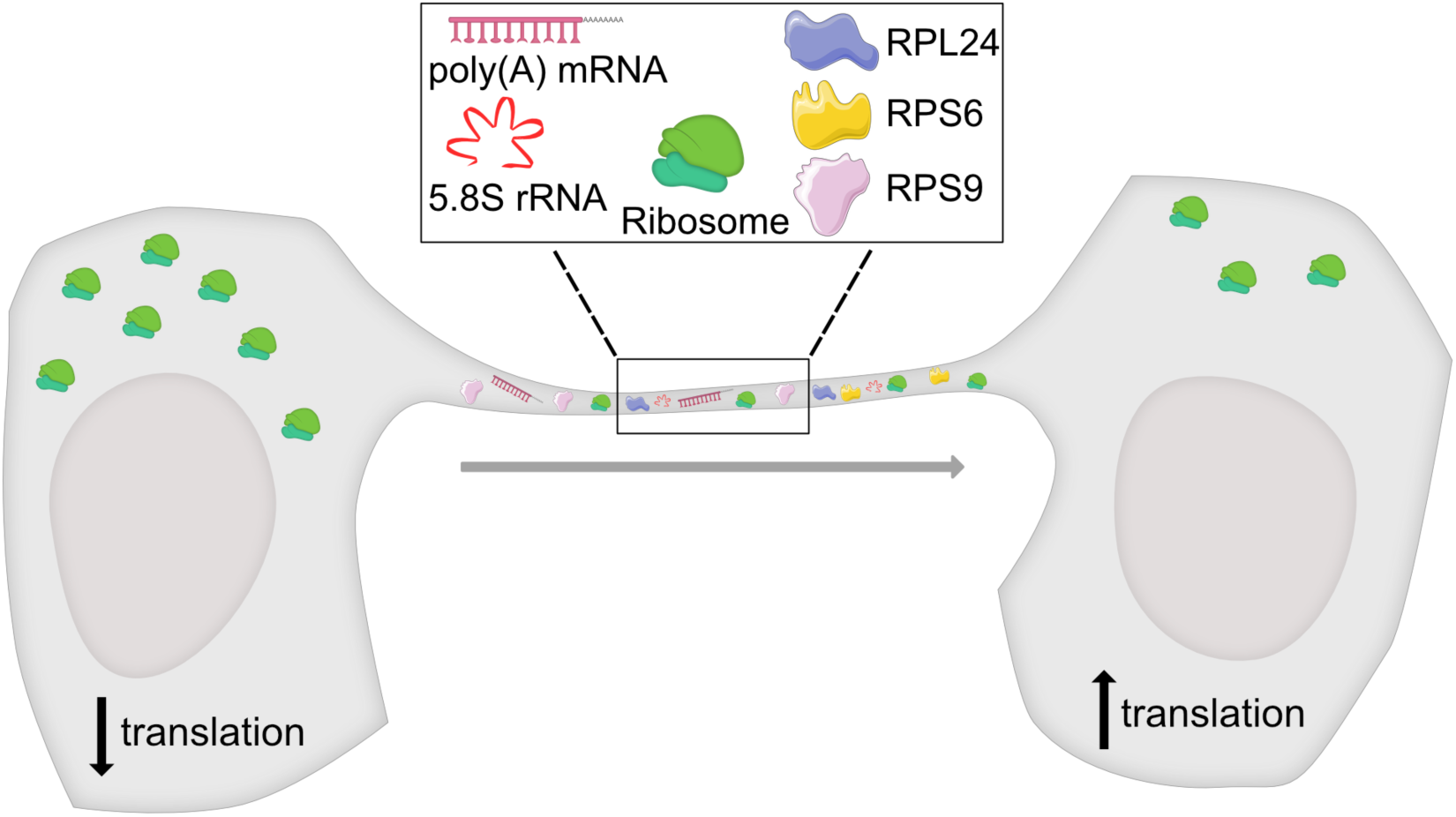

**Key points:**

- pancreatic tumour cell lines and cells from patient biopsies form tunnelling nanotubes in 2D culture
- formation of tunnelling nanotubes is promoted by gemcitabine
- polyadenylated mRNAs, ribosomal components and assembled ribosomes are present in tunnelling nanotubes
- ribosomes and their components are transferred via nanotubes to acceptor cells
- silencing of ribosomal proteins S6 and L24 reduces the number of assembled ribosomes
- global protein synthesis and number of ribosomes in pancreatic cancer cells with silenced ribosomal proteins increases when co-cultured with translationally unimpaired cells

**Background:** Pancreatic ductal adenocarcinoma (PDAC) is considered as one of the deadliest types of cancer. Tunnelling nanotubes (TNTs) are thin, membranous, intercellular communication structures observed in normal and cancer cells, where they mediate the exchange of intracellular material and promote cell fitness, cancer spread and treatment resistance.

**Results:** PDAC cells increase the formation of TNTs upon exposure to gemcitabine. In the PANC-1 cell line and in tumour explants from patients, we observe polyadenylated mRNA, 5.8S rRNA, ribosomal proteins and assembled 80S ribosomes within the TNTs. Using HaloTag-labelled small ribosomal subunit component RPS9 we demonstrate the transport of ribosomes via TNTs into acceptor cells. Downregulation of ribosomal proteins S6 and L24 decreases the number of assembled ribosomes and the global protein translation in PDAC cells, while a co-culture with translationally unimpaired cells partially restores protein synthesis in cells with impaired protein translation.

**Conclusions:** PDAC cells can exchange components of the protein translation machinery and mRNA. The intercellular transfer of these components causes a partial restoration of protein translation in cells with impaired protein synthesis, which may contribute to the resilience of pancreatic cancer cells, highlighting the potential of targeting TNT dynamics as a therapeutic approach for PDAC.

## INTRODUCTION

Pancreatic ductal adenocarcinoma (PDAC) is one of the deadliest cancers with a poor prognosis with a five year overall survival rate of approximately 10%.^1^ One of the reasons for the aggressive nature of PDAC is its ability to develop chemoresistance, which ultimately leads to treatment failure and disease progression. Detecting this aggressive cancer in its early stages is crucial for successful treatment outcomes. Gemcitabine monotherapy serves as the established standard of care for patients with worse performance status diagnosed with locally advanced and metastatic pancreatic adenocarcinoma.^2^ For patients with good performance status, the FOLFIRINOX (oxaliplatin, irinotecan, leucovorin, fluorouracil) regimen is indicated as the method of first choice, as it has shown a significant prolongation of median overall survival.^3^

The upregulation of ribosome biogenesis is an important characteristic of rapidly dividing cells,^4^ EMT, metastasis^5^ and also of the stem cell phenotype, where the maintenance of long-term replicative plasticity requires a high rate of ribosome synthesis.^6^ Dependence on a high rate of ribosome biogenesis has been documented also in cancer stem cells and related to poor disease prognosis.^7^ The inhibition of rRNA production has been shown to inhibit tumour growth.^8^ An inhibitor of RNA polymerase I exhibited antiproliferative activity in cancer cells and tumour xenografts,^9–11^ further supporting the importance of ribosomal biogenesis for cancer cell population.

Tunnelling nanotubes (TNTs) are intercellular direct cytoplasmic communication channels that allow cells to transmit signals and cellular components over long distances.^12^ Tunnelling nanotubes were first observed in rat pheochromocytoma PC12 cells, as well as in human embryonic kidney cells and normal rat kidney cells. These structures typically had a diameter ranging from 50 to 200 nm and could extend several cell diameters in length. Importantly, they did not lie on the substrate.^13^ Typically reported intercellular distances are between ten to several hundred µm.^14,15^ TNTs can transport mitochondria,^16^ ions,^17^ organelle-derived vesicles (endosomes, lysosomes), proteins, non-coding RNA.^16,18–20^ Recent studies have documented the role of TNTs in the transmission of SARS-CoV-2 and Zika virus.^21,22^ In cancer research, TNTs have been observed transporting mitochondria in glioblastoma patient-derived organoids^23^ and in mesenchymal stem cells transferring mitochondria to glioblastoma stem cells and enhancing resistance to temozolomide;^24^ in breast cancer cells transferring mitochondria from immune cells.^25^ In prostate cancer, TNTs contribute to the adaptation and treatment resistance through mechanisms involving PI3K/AKT signalling, actin remodelling, and the transport of stress-induced proteins;^26^ in bladder cancer TNTs connect heterogeneous cells and facilitate the mitochondrial transfer, leading to increased invasiveness both *in vivo* and *in vitro.*^27^ The formation of TNTs can also be induced in a dose-dependent manner by doxorubicin *in vitro* with the resulting drug redistribution via TNTs presenting a new addition on cellular mechanisms of drug efflux and the development of drug resistance in cancers.^28^ TNT formation can also be triggered by low serum, hyperglycemic, acidic growth medium^29^ and hypoxia.^30^ TNTs are also known to play a role in tumour progression, invasiveness and metastasis.^27,31,32^

In this study, we investigated whether sub-IC50 doses of the chemotherapeutic agent gemcitabine may enhance the formation of tunnelling nanotubes in PDAC cells. When we detected ribosomal components and fully assembled ribosomes in TNTs formed between pancreatic cancer cells we further wanted to see, whether this intercellular transport of the translational machinery may increase protein synthesis in cells with impaired protein synthesis.

## RESULTS

### PDAC biopsy samples develop tunnelling nanotubes in 2D cultures

TNTs have previously been detected in cultures of pancreatic cancer cell lines and tumour tissue sample.^28^ We used 2D cultures of patient explants from endoscopic biopsies to see whether cultured primary cancer cells can also form these connections. We assessed the formation of tunnelling nanotubes in three tissue samples obtained from patients diagnosed with pancreatic ductal adenocarcinoma following endoscopic ultrasound-guided fine needle aspiration biopsy. The patients had not undergone any chemotherapy before the biopsy and their characteristics are summarised in Supplementary Table S1. TNTs were observed in all three cultured tissue samples (Figure 1A, Supplementary Figure S1A). Cells started growing out from the biopsy fragments between 2 to 5 days after plating in a growth medium. All three patients received palliative care. Given that gemcitabine is the most frequently used first-line treatment for palliative chemotherapy in unresectable PDAC, we investigated whether gemcitabine could affect the formation of TNTs in culture.

**Figure 1.**
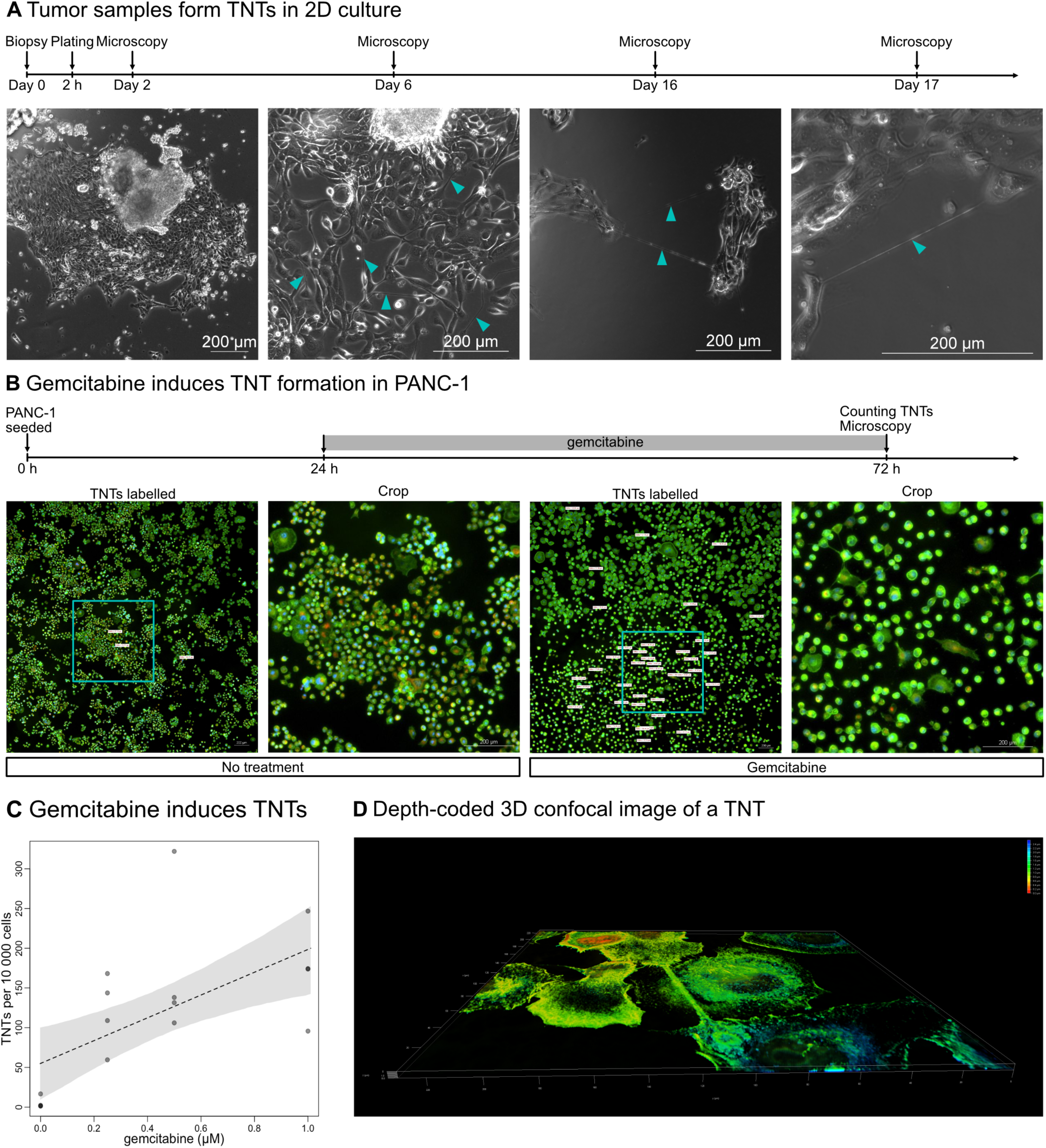
Tunnelling nanotubes form in 2D cultures of primary PDAC and in PANC-1 cell line treated with gemcitabine. **(A)** Cells in 2D culture of fine-needle biopsy sample from a PDAC patient BJPN15 develop TNTs. **(B)** PANC-1 cell line increases the number of TNTs in response to gemcitabine. Green, CellMask Green (actin); red, TMRM (mitochondria); blue, Hoechst 33342 (DNA). **(C)** Linear model of the dose-dependence of TNT formation on gemcitabine concentration. **(D)** Confocal 3D reconstruction of one TNT between PANC-1 cells with depth coding showing that the TNT is above the substrate. Further images in Supplementary Figure S1.

### Gemcitabine induces TNTs formation in PANC-1 in dose-dependent manner

To determine the optimal dosing regimen, we first estimated the IC50 for gemcitabine in PANC-1 cells to be 1.6 μM (Supplementary Figure S1B). The dense stromal composition and poor vascularization characteristic of PDAC hinder effective drug penetration into the tumour *in vivo*, therefore we used sub-IC50 concentrations for further experiments. We exposed PANC-1 cells to three different sub-IC50 doses of gemcitabine: 0.25 μM, 0.5 μM, and 1 μM for 48 hours and observed a dose-dependent stimulation of the formation of tunnelling nanotubes (Figure 1B, 1C), considering only tunnelling nanotubes measuring 20 µm or more in length. The observed tunnelling nanotubes form above the substrate (Figure 1D), as expected from these structures. This observation suggests that sublethal doses of gemcitabine induce the formation of TNTs and potentially influence the survival and growth ability of pancreatic cancer cells.

### Messenger RNA, ribosomal components and assembled ribosomes are transported in tunnelling nanotubes

Since rapidly dividing cancer cells are highly dependent on proteosynthesis we hypothesised that RNAs and components of the translation machinery may also be transported through TNTs between PDAC cells. Following a treatment with 0.5 µM gemcitabine for 48 hours to induce the formation of TNTs we detected 5.8S ribosomal RNA (rRNA) within these structures (Figure 2A and Supplementary Figure S2A, S7A). A treatment with RNAse eliminated the RNA signal in the samples (negative control, Supplementary Figure S7A). 5.8S ribosomal rRNA, which is evolutionarily conserved and a critical component of the large subunit of eukaryotic ribosomes, plays an essential role in protein synthesis.^33^ Furthermore, we identified 5.8S rRNA within the TNTs between pancreatic ductal adenocarcinoma cells and between stromal cells from a patient’s biopsy (Figure 3A, B and Supplementary Figure S3A, B).

**Figure 2.**
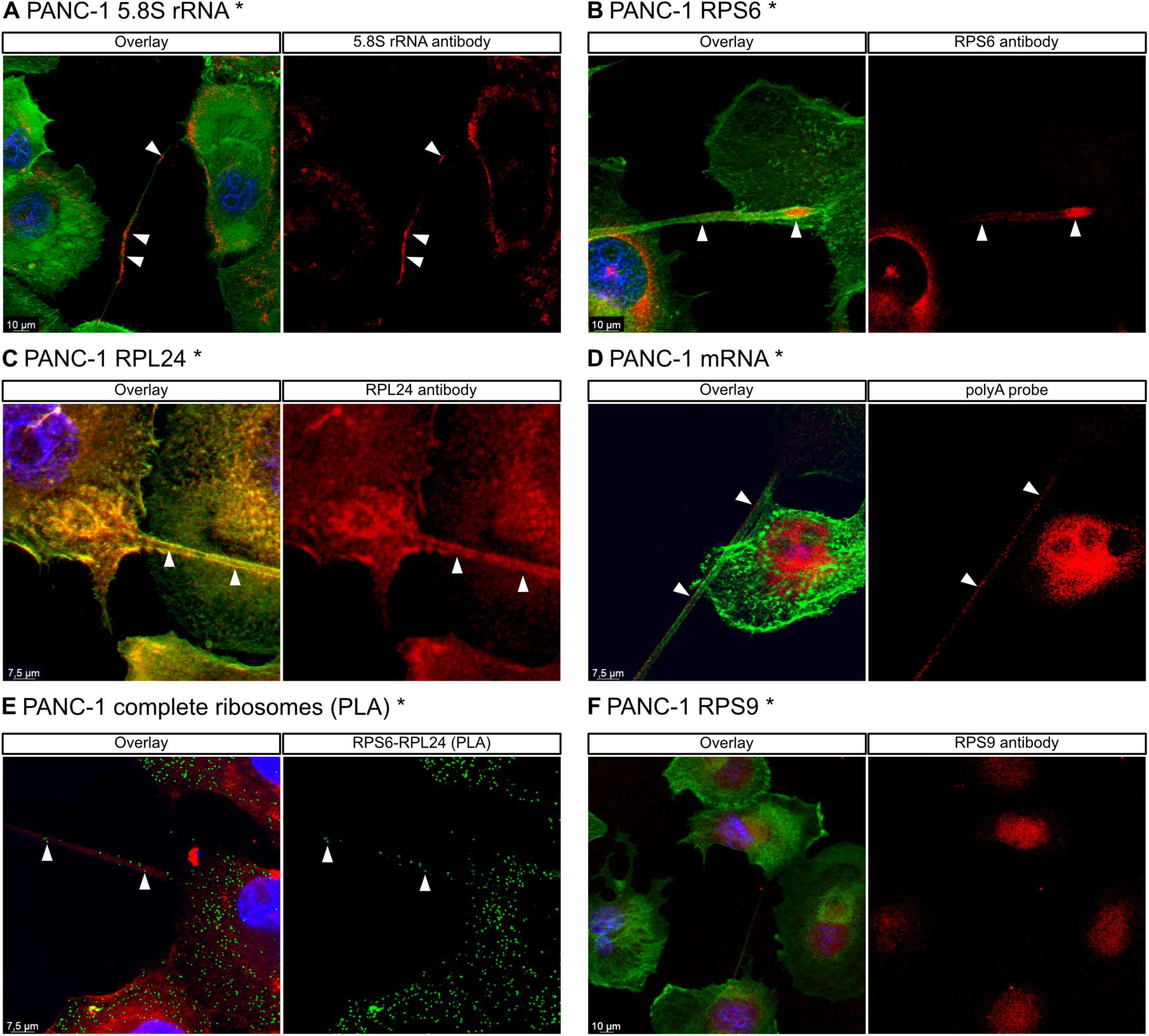
Ribosomal components, mRNA and fully assembled ribosomes are detected in PANC-1 tunnelling nanotubes. Ribosomal components: **(A)** 5.8S rRNA, **(B)** ribosomal protein S6, **(C)** ribosomal protein L24, **(D)** poly-A mRNA **(E)** complete ribosomes were detected by a proximity ligation assay (PLA) using anti-RPS6 and anti-RPL24 antibodies and **(F)** ribosomal protein S9 found in TNTs formed between PANC-1 cells in culture after a 48-hour exposure to 0.5 µM gemcitabine. Images from a confocal microscope (*) and from a fluorescent microscope (#). Further images in Supplementary Figure S2.

**Figure 3.**
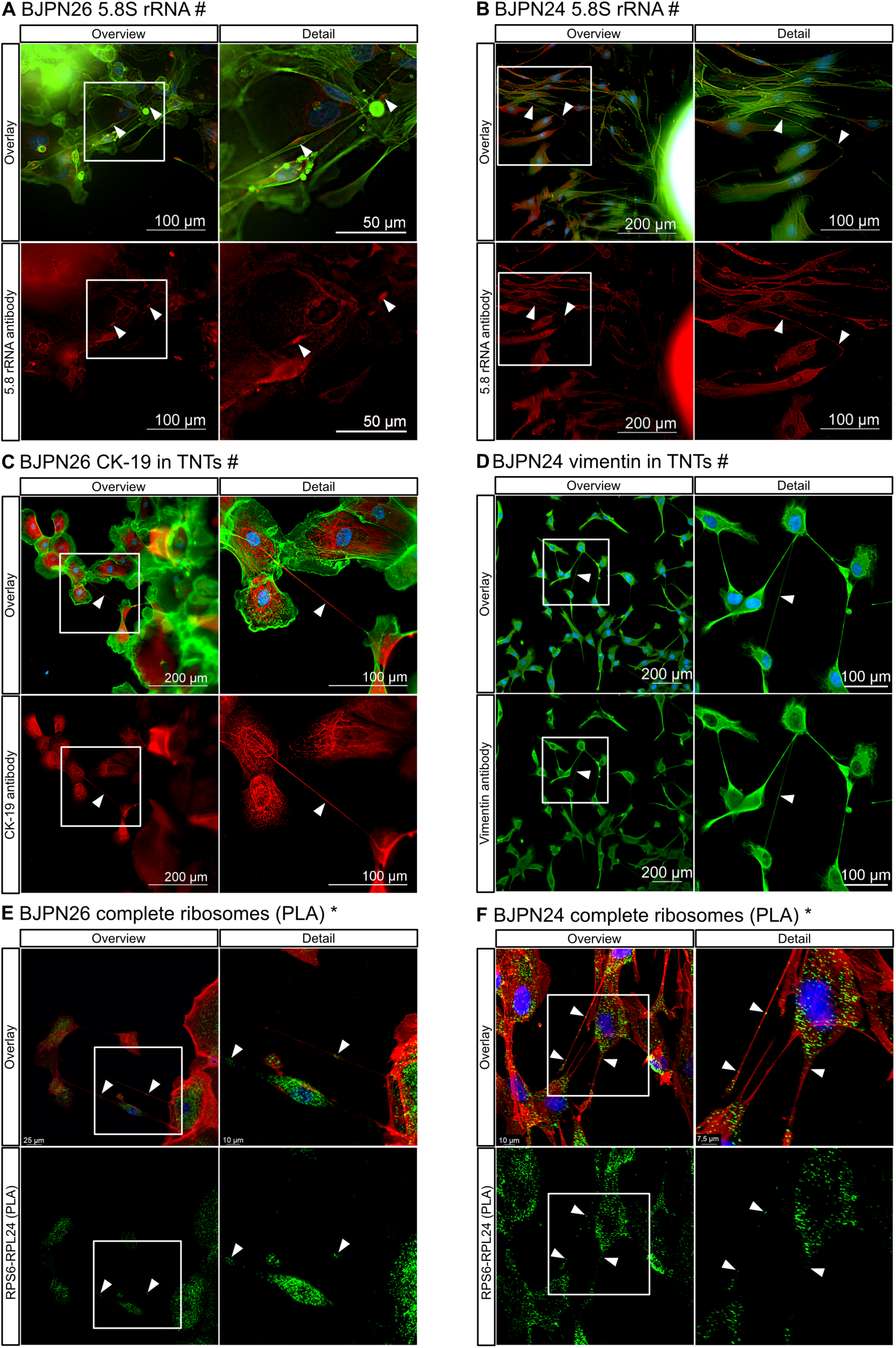
Detection of rRNA, fully assembled ribosomes, and cytoskeletal markers in TNTs formed between patient-derived cells BJPN26 and BJPN24. **(A, B)** Ribosomal 5.8S rRNA was detected in TNTs formed between patient-derived cells. **(C)** TNTs formed between BJPN26 cells in 2D culture, accompanied by cytokeratin-19 (CK-19). (**D)** TNTs formed between BJPN24 cells in 2D culture, accompanied by vimentin. **(E, F)** Fully assembled ribosomes were detected in BJPN26 and BJPN24 cells in 2D culture using a proximity ligation assay (PLA) with anti-RPS6 and anti-RPL24 antibodies. Images from a confocal microscope (*) and from a fluorescent microscope (#). Further images in Supplementary Figure S3.

We then set out to identify other components of ribosomes in TNTs and found the presence of ribosomal protein S6 and S9 (key components of the 40S subunit) and, ribosomal protein L24 (key component of the 60S subunit) in gemcitabine-activated TNTs in PANC-1 cells (Figure 2B, C, F and Supplementary Figure S2B, C, F) suggesting the possibility that fully assembled ribosomes may be transported between cancer cells via TNTs.

To visualise polyadenylated RNA (mRNA) in tunnelling nanotubes we performed *in situ* hybridization with RNA oligo-d(T) probes (Figure 2D and Supplementary Figure S2D, S7D, RNAse-treated negative control Supplementary Figure S7D). The presence of polyadenylated RNAs in TNTs complements the detection of ribosomal components and suggests the possibility of transfer of functional ribosomes and mRNA between cells via TNTs.

### Complete ribosomes are transferred in TNTs

To investigate whether the presence of 5.8S ribosomal RNA (rRNA), both ribosomal subunits and polyadenylated mRNA within tunnelling nanotubes supports the hypothesis that functional ribosomes can be transferred between cells, we used the proximity ligation assay (PLA) to detect fully assembled, translationally competent ribosomes.^34,35^ We used primary antibodies targeting RPL24 and RPS6, spatially close ribosomal proteins from the 60S, respectivelly the 40S ribosomal subunit,^36^ to generate a multiplexed fluorescent signal through secondary antibodies with hybridization probes that produced single spots indicative of functional 80S ribosomes (Figure 2E and Supplementary figure S2E, S7E). The assembled ribosomes identified by the RPS6-RPL24 interaction were distributed in the cytoplasm of PANC-1 cells and in intercellular TNTs. The interaction between RPS6 and RPL24 indicates the formation of fully assembled ribosomes and ongoing protein synthesis.^34,35,36^ Elimination of assembled ribosomes by a treatment with puromycin,^37^ significantly reduced the RPS6-RPL24 PLA signal (Supplementary figure S7E), confirming the fact that we only obtained signals from assembled ribosomes in the cytoplasm and TNTs (Figure 2E and Supplementary figure S2E, S7E). Our results clearly show that tunnelling nanotubes contain fully assembled ribosomes *en route* between cancer cells.

### TNTs are present between primary PDAC cancer cells, fibroblasts from biopsy samples and contain assembled ribosomes

To establish that ribosome transport via tunnelling nanotubes is not exclusive to the PANC-1 cell line but also occurs in primary human pancreatic tumour cells, we cultured primary human PDAC cells in a 2D culture from thin needle biopsy samples. We detected 5.8S rRNA within TNTs in two patient samples BJPN26 and BJPN24 (Figure 3A, B and Supplementary Figure S3A, B). To ascertain the presence of epithelial tumour cells *in vitro*, we used a primary antibody against cytokeratin-19, while the detection of stromal cells was confirmed by a primary antibody against vimentin. One patient sample BJPN26 was positive for epithelial cells (Figure 3C, Supplementary Figure S3C), while the other BJPN24 contained vimentin-positive stromal fibroblasts without any cytokeratin signal (Figure 3D, Supplementary Figure S3D). Both patient samples expanded in culture and formed TNTs, which, interestingly, contained significant signals of the two intermediate filament proteins, cytokeratin-19 or vimentin, respectively. Using the RPS6-RPL24 proximity ligation assay we detected assembled ribosomes in these TNTs (Figure 3E, F and Supplementary Figure S3E, F). These data show that functional assembled ribosomes are present in TNTs of primary PDAC cells, as well as in those formed between primary stromal cells.

### Ribosomal components and assembled ribosomes are transported from donor cells to acceptor cells via TNTs

To demonstrate that ribosomal components are transported from donor to acceptor cells via gemcitabine-activated TNTs, inspired by Gallo *et al.*,^38^ we generated PANC-1 cells transfected with a mammalian expression vector encoding the human small ribosomal subunit RPS9, C-terminally tagged with HaloTag (donor cells). These transfected donor cells expressed tagged RPS9 in the nucleolus, nucleus, and cytoplasm (Supplementary Figure S4A), and a time-lapse video shows the movement of the labelled ribosomal component (Video 1).

Acceptor cells were stably labelled either by staining with the covalent CellTrace Violet stain (CellTrace Violet+) or by a transfection with GFP with NLS localized in the nucleus. After washing and trypsinization, equal numbers of donor and acceptor cells were seeded and cultured in 24-well plates. Next, co-cultured donor and acceptor cells were exposed to gemcitabine for two days to induce the formation of TNTs. Subsequently, the cells were labelled with HaloTag-TMR ligand (HaloTag TMR+), allowing for the observation of double-stained cells, in some cases with a TNT connecting a HaloTag positive donor cell with an acceptor cell (Figure 4A, B and Supplementary Figure S4A, S4B). These observations suggest that HaloTag-RPS9 was exchanged between the two cell populations through induced TNTs, but a possible contribution of mRNA or protein transfer mediated by extracellular vesicles cannot be excluded.

**Figure 4.**
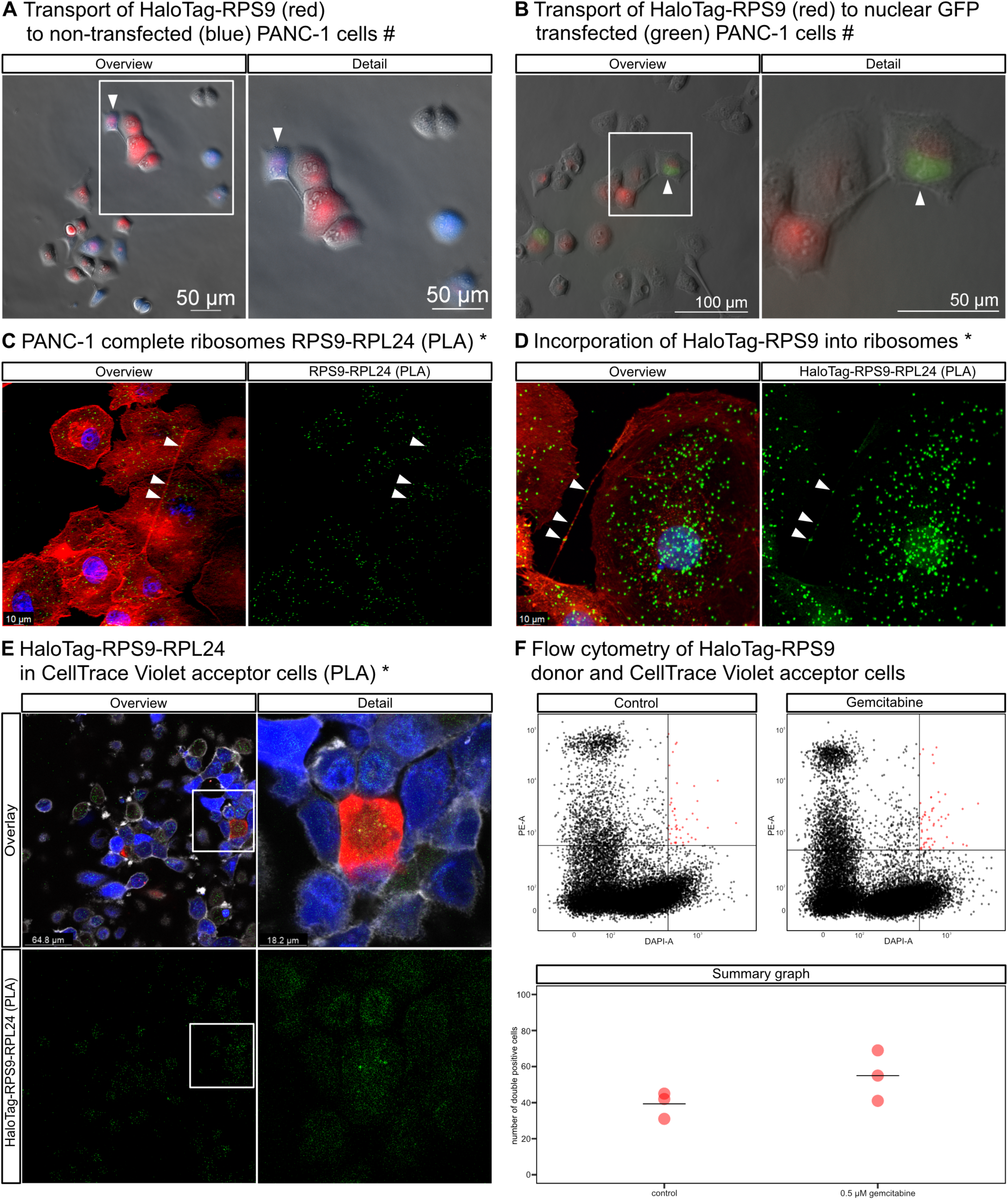
Ribosomal component RPS9 is transported between cells via TNTs and incorporated into ribosomes. HaloTag-labelled RPS9 (red) was transported **(A)** to non-transfected PANC-1 cells stained with a blue CellTrace Violet stain (blue) or **(B)** to PANC-1 cells transfected with nuclear targeted GFP (green). **(C)** Complete ribosomes (green spots) were detected by a proximity ligation assay (PLA) with anti-RPS9 and anti-RPL24 antibodies in TNTs between PANC-1 cells. Nuclei are stained by DAPI (blue) and actin filaments are stained using the ActinRed Reagent (red). **(D)** Complete ribosomes (green spots) were detected using a proximity ligation assay (PLA) with anti-HaloTag-RPS9 and anti-RPL24 antibodies in TNTs between PANC-1 cells. Nuclei are stained by DAPI (blue) and actin filaments are stained using the ActinRed Reagent (red). **(E)** Alternatively, the HaloTag-RPS9–RPL24 PLA assay demonstrated that HaloTag-RPS9, labelled with HaloTag-TMR ligand (red) in donor non-stained cells, was subsequently transferred to non-transfected, CellTrace Violet-stained (blue) acceptor cells, where it was incorporated into fully assembled ribosomes (green spots). Actin filaments are stained using the CellMask Deep Red Actin Tracking Stain (white). **(F)** Transfer of HaloTag-RPS9 (red) to non-transfected CellTrace Violet (blue) cells detected by flow cytometry as double-positive cells is increased by sub-IC50 exposure to gemcitabine (mean counts of three experiments shown as lines in the summary graph). Images from a confocal microscope (*) and from a fluorescent microscope (#). Further images in Supplementary Figure S4.

To confirm our hypothesis that the intercellularly transferred HaloTag-RPS9 subunits are incorporated into complete ribosomes we first performed the RPS9-RPL24 proximity ligation assay to ascertain the necessary close proximity of these two subunits (Figure 4C and Supplementary Figure S4C). Next, we performed anti-HaloTag-anti-RPL24 proximity ligation assay and confirmed that the transgenic RPS9 subunits are also incorporated into 80S ribosomes (Figure 4D and Supplementary Figure S4D). Finally, we mixed donor HaloTag-RPS9 transfected cells with acceptor non-transfected CellTrace Violet-stained cells and performed the anti-HaloTag-anti-RPL24 proximity ligation assay. The positive PLA signal in CellTrace Violet-stained acceptor cells (Figure 4E and Supplementary Figure S4E) is a clear evidence for complete ribosomes in the Violet cells containing the HaloTag-RPS9 subunit transferred from the donor cells, which strongly supports our hypothesis. A negative control with cells without the HaloTag-RPS9 transgene is shown in Supplementary Figure S4E.

To support further the evidence for the transport of ribosomal components from donor to acceptor cells we used flow cytometry to quantify double-positive (HaloTag TMR+ and CellTrace Violet+) cells formed in a mixed population of HaloTag-RPS9 transgenic donor cells and non-transfected CellTrace Violet labelled acceptor cells (Figure 4F and Supplementary Figure S4F). The exposure to gemcitabine increases somewhat the number of double-stained cells, which is consistent with our observations above regarding the stimulation of TNT formation by this drug and with the hypothesis that TNTs are the conduit for transferring ribosomal components between cells.

### PDAC cells with gemcitabine-induced TNTs rescue global protein synthesis in cancer cells with depleted ribosomal proteins S6 and L24

To investigate whether PDAC cells with gemcitabine-induced tunnelling nanotubes can compensate for an impaired global protein synthesis in translation-deficient cells, we performed siRNA-mediated knockdown of RPS6 and RPL24—two essential components of the 40S and 60S ribosomal subunits. First, using Western blot analysis with primary antibodies against the ribosomal proteins S6 and L24, we demonstrated that PANC-1 cells transfected with siRNAs targeting both RPS6 and RPL24 simultaneously exhibited a consistent reduction in RPS6 and RPL24 protein expression over a three-day period (Figure 5A and Supplementary Figure S5A). To demonstrate a reduced global translational capacity in cells co-silenced with siRNA-RPS6 + siRNA-RPL24, we used the Click-iT HPG protein synthesis assay in combination with flow cytometry.^39^ To verify the cytometry-based protein synthesis assay, we treated PANC-1 cells with cycloheximide (CHX), which inhibited translation compared to non-treated control cells, resulting in reduced overall protein synthesis in CHX-treated cells (Figure 5B). Figure 5C provides an overview of the experimental setup and workflow of the flow cytometry-based assay used to measure global protein synthesis using the Click-iT translation assay.

**Figure 5.**
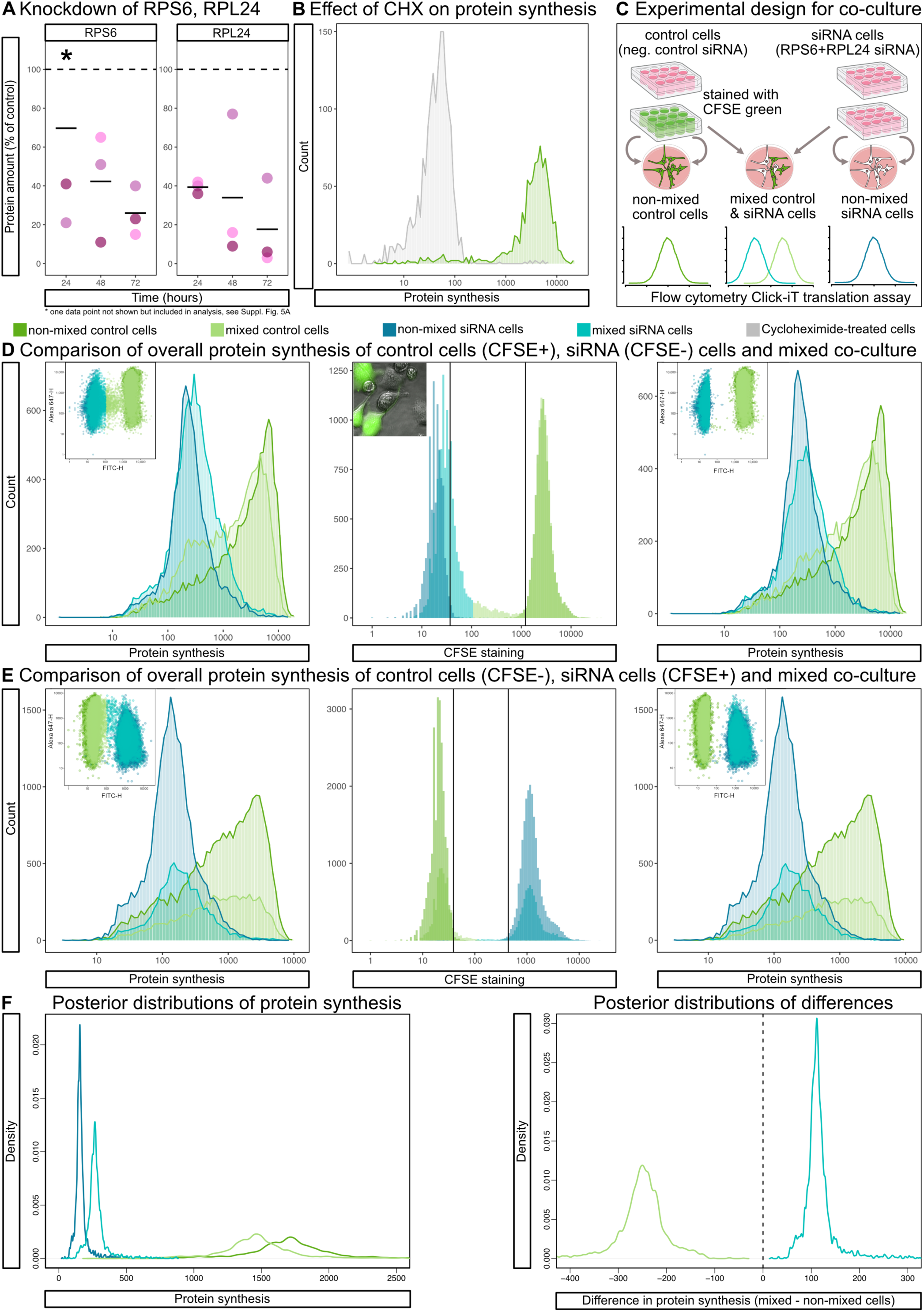
siRNA targeting ribosomal proteins S6 and L24 reduce the levels of both ribosomal proteins and led to a decrease in overall protein synthesis in PANC-1 cells. Co-culture of siRNA cells with translation-competent control cells increase protein synthesis in translationally impaired siRNA cells. **(A)** The Western blot dot plot shows the levels of ribosomal proteins S6 and L24 following siRNA-mediated knockdown of RPS6 and RPL24 in PANC-1 cells over a three-day period, compared to control cells (100%), from three biological replicates. **(B)** The flow cytometry histogram illustrates a reduction in overall protein synthesis in PANC-1 cells detected using the Click-iT protein synthesis assay following a treatment with cycloheximide (CHX) compared to control cells. **(C)** Graphical overview of the co-culture experiment for measuring global protein synthesis in mixed cultures by the Click-iT assay. **(D)** A comparison of overall protein synthesis of control cells stained with CFSE, non-stained siRNA cells and mixed control (CFSE) & siRNA cells (non-stained). Miniature scatter plots show the fluorescence intensity of translation (Alexa 647-H) and CFSE staining (FITC-H). The middle histogram and microscopic image show partial CFSE transfer from control cells to siRNA cells via TNTs and the stricter gates (black lines) to exclude partially stained cells. The third plot represents a comparison of overall protein synthesis using stricter gate settings to exclude partially stained cells. **(E)** A comparison of overall protein synthesis of non-stained control cells, siRNA cells stained with CFSE and mixed control (non-stained) & siRNA cells (CFSE) in colour-reversed experiment. Miniature scatter plots show the fluorescence intensity of translation (Alexa 647-H) and CSFE staining (FITC-H). The middle histogram shows partial CFSE transfer from siRNA cells to control cells and the stricter gates (black lines) to exclude partially stained cells. The third plot represents a comparison of overall protein synthesis using stricter gate settings to exclude partially stained cells. **(F)** Bayesian model posterior predictions of overall protein synthesis in average populations of control cells stained with CFSE, non-stained siRNA cells and mixed control (CFSE+) & siRNA cells (CFSE-) using the strict gating strategy with clear differences between mixed and non-mixed populations showing the overall effect of mixed co-culture. Further graphs and images in Supplementary Figure S5.

We then hypothesised that a co-culture of siRNA-negative control PANC-1 cells (translationally unimpaired) with PANC-1 cells co-silenced with siRNA-RPS6 + siRNA-RPL24 (translationally impaired) under treatment with 0.5 μM gemcitabine for 24 hours (a condition that induces TNT formation and facilitates the intercellular exchange of ribosomal components) could lead to a net transfer of ribosomal components from unimpaired to impaired cells and increase the global protein translation rate in the latter cells. Therefore, we performed flow cytometry analysis in which siRNA-negative control PANC-1 cells (control cells) were stained with the green CellTrace CFSE Cell Proliferation Kit, enabling their distinction from PANC-1 non-stained cells co-silenced with siRNA-RPS6 + siRNA-RPL24 (siRNA cells) within the mixed co-culture experimental group (Figure 5C). Using flow cytometry in combination with the Click-iT HPG protein synthesis assay we observed a reduction in overall protein translation in PANC-1 cells transfected for two days with siRNAs co-targeting RPS6 and RPL24, relative to cells treated with a siRNA-negative control (Figure 5D). These results show that ribosomal proteins S6 and L24 are essential for maintaining global protein translation levels and strongly support the previous observation that the proximity and interaction of these two ribosomal proteins are critical for the functional assembly of ribosomes.^34^

Importantly, we found that, in a mixed co-culture siRNA cells (mixed siRNA) had a higher global protein synthesis compared to siRNA cells in non-mixed control group, whereas mixed control cells (siRNA negative control) showed a relative decrease in protein synthesis compared to their respective non-mixed control group (Figure 5D and Supplementary Figure S5D). This observation is consistent with an intercellular ribosomal transport from translationally unimpaired to translationally impaired cells with a clear impact on protein translation in both donor and acceptor cells.

During these experiments we noticed that Carboxyfluorescein Succinimidyl Ester (CFSE)—a dye that covalently labels intracellular proteins by reacting with amine groups and is detectable in the FITC channel—is partially transferred via TNTs from siRNA-negative control cells (CFSE) to cells co-silenced with siRNA-RPS6 + siRNA-RPL24 (non-stained) (Figure 5D and Supplementary Figure S5D). This observation could potentially affect the interpretation of our results as it blurs the distinction between stained and unstained cells.

We therefore used two different fluorescence-intensity gates to distinguish the two cell populations: in the first instance we defined positive (stained) cells as those with green fluorescence intensity above the maximum of non-stained cells, and as a stricter setting we excluded from the analysis cells, whose green fluorescence intensities fit between the non-stained and stained populations (i.e. cells which may have originated as non-stained cells but received some green-stained protein via TNTs or other means). Since both gating settings yielded similar results we can be confident that the partial intercellular dye transfer did not meaningfully influence the conclusion of this experiment (Figure 5D and Supplementary Figure S5D).

To validate our findings and exclude any possible confounds due to the asymmetric staining, we performed a colour-reversed experiment in which cells co-silenced with siRNA-RPS6 + siRNA-RPL24 with reduced translation were stained with CFSE, while siRNA-negative control cells remained unstained (Figure 5E and Supplementary Figure S5E). In this reverse experiment, we once again used the two gating strategies described above (Figure 5E and Supplementary Figure S5E). The observed changes in overall protein synthesis between mixed and non-mixed control and siRNA cells remained consistent.

In order to analyze these data statistically we used a linear, varying intercepts Bayesian model (see Methods) for posterior predictions of average populations and differences between them (Figure 5F and Supplementary Figure S5F). The main figure shows the posterior predictions from first experimental setup gated using the more strict gating strategy, while the supplementary figure shows posterior prediction from the original experimental setup using the standard gating strategy and from the colour-reversed setup using both gating strategies, as described above. In all posterior predictions the probability densities of differences between mixed vs non-mixed populations are overwhelmingly different from zero, suggesting a significant effect of mixed co-culture on global protein synthesis in PANC-1 cells. We can thus conclude that the spatially close presence of translationally unimpaired control cells can enhance overall protein synthesis in cells with impaired protein translation, presumably via the intercellular transfer of ribosomal components or entire ribosomes via TNTs.

### PDAC cells with gemcitabine-induced TNTs increase ribosome counts in cancer cells with depleted ribosomal proteins S6 and L24

We then asked whether the observed reduction in overall protein synthesis in cells transfected with siRNAs targeting RPS6 and RPL24, compared to siRNA-negative control cells, may be due to an impaired assembly of functional ribosomes following the gene silencing. Thus, if siRNA-negative control cells can transfer ribosomes to siRNA RPS6 + RPL24 knockdown cells via tunnelling nanotubes, we should be able to detect an increase in ribosome counts in siRNA RPS6 + siRNA-RPL24 cells co-cultured with siRNA-negative control cells.

Therefore, we used the proximity ligation assay and confocal microscopy to quantify the number of assembled ribosomes per cell in a parallel setup to the global translation assay described above to see if the observed changes in protein translation will be reflected in ribosome counts per cell. Cells (siRNA-negative non-mixed control group stained with CFSE, siRNA non-mixed control group, the mixed co-culture of control (CFSE) and siRNA cells) were seeded on glass coverslips, treated with gemcitabine for 29 hours, fixed, and subjected to PLA using antibodies against RPS6 and RPL24. Imaging was performed using confocal microscopy across multiple fields of view. Each scan was reconstructed as a maximum intensity projection from 13 Z-stack slices. For each complete, non-overlapping cell, a region of interest (ROI) was defined based on actin boundaries. Within each ROI, assembled ribosomes were counted as distinct PLA-detected fluorescent spots.

The results revealed a significant reduction in the number of fully assembled ribosomes in PANC-1 cells co-silenced with RPS6 and RPL24, compared to siRNA-negative control cells (Figure 6A and Supplementary Figure S6A), suggesting that the observed decrease in global protein synthesis in siRNA cells is indeed due to reduced ribosome assembly. Importantly, the number of fully assembled ribosomes in siRNA cells co-cultured with siRNA-negative control cells increased compared to their respective non-mixed control group, and decreased in mixed siRNA negative control cells compared to non-mixed cells (Figure 6A and Supplementary Figure S6A). For a statistical analysis of ribosome count data we used a Bayesian model (see Methods) and the predicted posterior distributions and differences between average cell populations are shown in Figure 6B. This finding is consistent with the observed changes in translation in these cells and is consistent with the hypothesis that the increase in number of ribosomes results from the intercellular transfer of ribosomes from siRNA-negative control cells.

**Figure 6.**
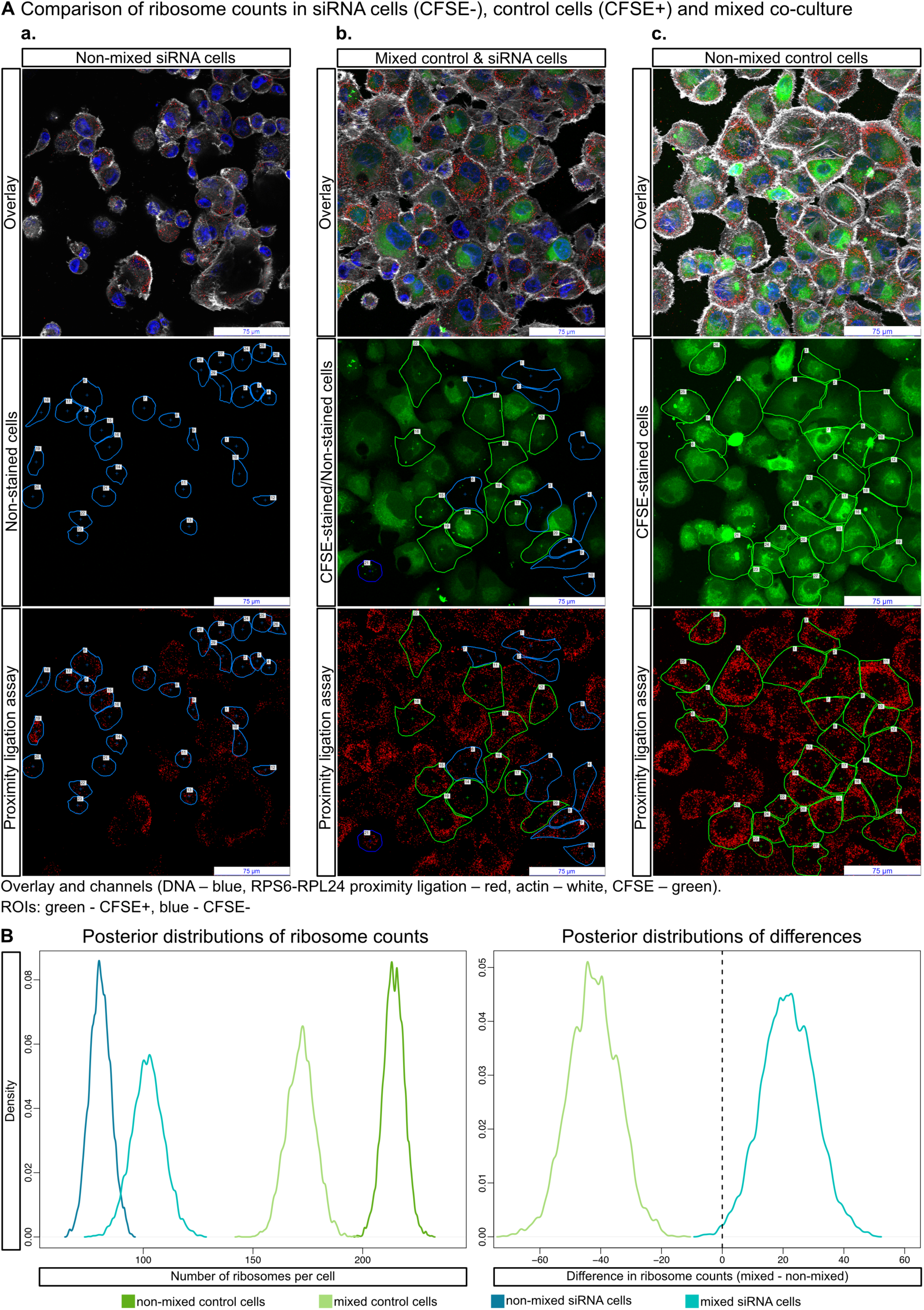
Co-culture of siRNA cells with translation-competent control cells increases the number of fully assembled ribosomes in translationally impaired siRNA cells. **(A)** Microscopic field reconstructed as a maximum intensity projection from 13 Z-stack slices displays: (a) siRNA non-mixed cells (non-stained), (b) mixed co-culture of control cells (CFSE) & siRNA cells (non-stained), (c) control non-mixed cells (CFSE). Regions of interest (ROIs) of non-overlapping cells were defined based on actin boundaries (siRNA cells – blue ROIs; control cells – green ROIs). The copied ROIs were applied to the red channel image, where fluorescent spots represent fully assembled ribosomes detected by PLA. Quantification of fluorescent spots in the defined ROIs was performed using NIS-Elements AR Software. **(B)** Bayesian model posterior predictions for average cell populations in the four experimental groups show clear differences in ribosome counts per cell. Further images in Supplementary Figure S6.

**Video 1. Live cell imaging of HaloTag-RPS9 transport in TNTs between PANC-1 cells.** A movement of the HaloTag-RPS9 labelled with HaloTag-TMR ligand (red) within TNTs formed between PANC-1 cells in culture after a 48 hours exposure to 0.5 µM gemcitabine (3.873 seconds per frame). Nuclei are stained by Hoechst (blue) and actin filaments are stained using the CellMask Green Actin Tracking Stain (green).

## DISCUSSION

Our results show that ribosomal components 5.8S rRNA, RPS9, RPS6, RPL24, and fully assembled ribosomes together with mRNA are present in TNTs between pancreatic cancer cells, and that they are transported between cells. This transport of translational machinery can, furthermore, increase the number of ribosomes and overall protein synthesis rates in translationally impared cells. These findings expand the current knowledge about the nature of intracellular material transported via TNTs and show a possible new direction in cancer biology research.

As stated in the Introduction, ribosomal biogenesis has been shown as one of key determinants of cancer cell survival, proliferation, adaptation and spread leading to the development of specific inhibitors of rRNA synthesis as possible anticancer agents.^8^ Currently used chemotherapeutic agents have also been shown to inhibit rRNA synthesis and ribosome biogenesis, which may contribute to their anticancer effects.^40^ Since the intercellular transfer of mitochondria has been proposed as an adaptive mechanism capable of restoring defective mitochondrial function in acceptor cells,^41–44^ it is perhaps not too far-fetched to see the potential adaptive impact of ribosome transport in cancer cells, which are in many instances dependent on an upregulated rate of ribosome biogenesis and proteosynthesis. The formation of TNTs and their enabling of the transfer of ribosomes and gene transcripts may be part of the response to chemotherapy-induced cellular stress and contribute to the survival of acceptor cells.

Our results show that sub-IC50 concentrations of gemcitabine induce the formation of TNTs between cancer cells. Since TNTs provide the means of redistributing intracellular components from one cell to another, in the case of mitochondria with a proven adaptive impact,^16,24,25^ it is possible to see the induction of TNT formation by gemcitabine as a potential contributor to chemoresistance. The fact, reported in the present study, that ribosomes and ribosomal components are also transferred between cells may further contribute to this pro-survival and pro-proliferative potential of TNT formation.

The phosphorylation of RPS6 via an aberrant activation of the mTOR pathway is a significant factor in the initiation and progression of pancreatic cancer^45^ and is related to metastasis and poor prognosis in other types of cancer,^46^ which presents a possible therapeutic target.^47^ Studies have also shown that the inhibition of mTORC1 decreases the proliferative ability of PDAC cells *in vitro* and *in vivo,*^48^ and the chemosensitivity to gemcitabine potentiated by inhibiting proteosynthesis.^49^ The apparent dependence of PDAC cells on RPS6 phosphorylation and increased rate of proteosynthesis may indicate a further adaptive use for the transfer of ribosomes between cells with different S6 kinase activities and ribosomal pools due to different nutrient status, growth factor signalling cascade activation, cellular stress levels or other factors.

Immunocytochemistry used in this study detects individual ribosomal components with the exception of the proximity ligation assay, which relies on the proximity of two proteins located in the small and large ribosomal subunit and produces signal only if the two subunits are close together. While our data show that complete 80S ribosomes are present in TNTs and exogenous HaloTag-RPS9 subunits are seen in complete 80S ribosomes in non-transfected acceptor cells, we cannot exclude the possibility that in addition to assembled ribosomes TNTs transport individual ribosomal components that are later formed into ribosomes in the acceptor cell.

The TNT-mediated intercellular transfer of cytosolic components other than ribosomes has been previously shown^16,18–20^ but the precise mechanisms driving these movements are still poorly characterised. The importance of the actin cytoskeleton for TNT formation and TNT-mediated transport is clear from numerous studies^13,50^ with recent reports implicating both actin and tubulin-linked motors in the transport.^51^ Whether ribosomes or their components are equipped with suitable adaptors for such cytoskeleton-based transport remains to be shown. While we detected ribosomes and ribosomal components in TNTs we cannot exclude the possibility that other means of intercellular transport may occur simultaneously, e.g. via extracellular vesicles.^52,53^

In this paper we demonstrate that the intercellular transfer of the translational machinery affects ribosome counts and global protein synthesis rates in the donor and acceptor cells in expected ways: the acceptor cells gain ribosome and proteosynthetic capacity, while the donor cells lose them. These findings regarding the intercellular restoration of impaired translation align conceptually with the recent work of Berridge *et al.* (2025), who highlight the therapeutic relevance of targeting intercellular transport mechanisms—such as tunnelling nanotubes— implicated in horizontal mitochondrial transfer that supports tumour cell survival and contributes to cancer progression. The trafficking of ribosomes and RNA and the associated regulation of translation between cancer cells may be another factor contributing to their resilience and adaptability, highlighting the pro-survival strategies that negatively influence cancer treatment. The mechanisms of ribosomal and mRNA transport via TNTs, its regulation and detailed effects of the transferred ribosomes and RNAs in the acceptor cells need to be further investigated in order to expand the possibilities for the treatment of patients with cancer. Since we have observed the formation of TNTs in both stromal cells and cancer cells, a detailed study of the heterotypic exchange of ribosomes within the tumour could provide further insights into the complex relationships between malignant and supporting cells.

## ACKNOWLEDGMENTS

This work was supported by funding by Institutional Research Concept RV067985904, Cooperatio Metabolic Diseases from Charles University; MZE-RO0723; CARDIA, project EXCELES LX22NPO5104 funded by Next Generation EU; and by grants from Charles University GAUK 108624 and GAUK 458225. This work was also supported by donations from Anděl Přerov s.r.o., Aliacem s.r.o., DG Solutions a.s. and Ing. Zdeněk Toman.

## AUTHOR CONTRIBUTIONS

S.M., J.T. and A.S. designed experiments. S.M. cultivated primary patient PDAC cells from biopsy, performed PANC-1 cell viability assay, cytometry, immunocytochemistry, transfection, western blots and fluorescence microscopy scanning of images. D.J. performed and analysed PLA assay, FISH, confocal scanning of images and analysed data from microscopy experiments. J.V. and L.J.L. performed TNT activation and TNT analysis. J.T., L.J.L., J.V. and M.H. analysed data from flow cytometry. P.D. performed cloning and verification of the HaloTag construct. M.B. and J.H. performed biopsies of PDAC patients and collected clinical data. Figures were designed and graphically edited by J.V., S.M., and J.T. S.M. and J.T. wrote the manuscript, all co-authors read and edited the final manuscript.

## RESOURCE AVAILABILITY

### Lead contact

Further information and requests for resources and reagents should be directed to and will be fulfilled by the lead contact, Jan Trnka

### Data and code availability

- All data reported in this paper is available at https://github.com/trnkaj/ribosomes.
- This paper reports original code available at https://github.com/trnkaj/ribosomes.
- Any additional information required to reanalyze the data reported in this paper is available from the lead contact upon request.

## METHODS

### Experimental model and subject details

#### Isolation of patients’ PDAC samples from EUS-guided FNAB

Patient samples were collected using fine-needle aspiration biopsy under endoscopic ultrasound (EUS-FNAB) control, which is the standard method for diagnosing pancreatic cancer. A mandrel aspiration needle was employed for the biopsy and removed after insertion into the lesion. Subsequently, several passes of the needle through the fixed lesion were performed under a vacuum, induced by a syringe with a fixable plunger position. The aspirate obtained by needle aspiration was transferred into a collection container with a transfer medium and into another container for cytological examination by the pathologist. Patient preparation for the procedure was similar to other therapeutic upper gastrointestinal endoscopies or biopsies of other abdominal organs. Patients arrived fasting, were informed about the purpose of the examination and its procedures, and provided their consent by signing an informed consent form. Standard analgosedation was used during the examination, and premedication with antibiotics was administered during the biopsy of cystic lesions. Patients were examined lying on their left side, with the possibility of additional position adjustments. Heart rate, blood pressure, and blood oxygen saturation were monitored throughout the procedure. The conclusion of the results, determined by the diagnostic pathologist, was based on the tissue samples examined in our study.

#### Cultivation of primary PDAC cells from biopsy tissues in 2D culture

Following EUS-FNAB, fresh patient-derived samples were immediately transferred into transfer medium (TM) composed of DMEM/F-12 Ham nutrient mixture, 10% foetal bovine serum, 1% Penicillin/Streptomycin, and 50 µg/mL Primocin at final concentrations, and transported on ice to the laboratory. Tissues were transferred using a pre-wetted pipette and tweezers onto a sterile glass Petri dish containing drops of wash medium with the same composition as the transfer medium. The tissue underwent five rounds of washing, with each round involving the gradual transfer of the tissue to new drops of wash medium. Subsequently, the tissue samples were minced into smaller fragments measuring 1–2 mm³ each using a scalpel in a separate drop of wash medium. These tissue fragments were then placed at the centre of a 12-well plate, pre-coated with rat tail collagen, and submerged in pre-warmed growth Biojepan culture medium (BCM). The BCM culture medium consisted of DMEM/Nutrient Mixture F-12 Ham, 2% foetal bovine serum, 2 mM GlutaMAX, 1% Penicillin/Streptomycin, 5 µg/mL insulin, 10 ng/mL epidermal growth factor, 0.4 µg/mL hydrocortisone, 10 µM Y-27632, and 50 µg/mL Primocin at final concentrations. The plate was transferred to an incubator set at 37°C and 5% CO_2_. The growth BCM medium was refreshed every 2–3 days.

Pancreatic cancer tissue was obtained from patients undergoing tissue biopsies at University Hospital Kralovske Vinohrady in Prague. Written informed consent was obtained from all patients. The study was approved by the Ethics Committee of the University Hospital Kralovske Vinohrady (EK-VP/05/0/2023) and adhered to the Declaration of Helsinki.

### Method details

#### PANC-1 cultivation and cell viability assay

PANC-1 cells were seeded at a density of 6 × 10^2^ viable cells per well in a 96-well white plate and cultured in 50 µL of DMEM supplemented with 10% foetal bovine serum and 1% Penicillin/Streptomycin (complete DMEM medium) at 37°C and 5% CO_2_. After 24 hours of cultivation, 50 µL of medium supplemented with varying concentrations of gemcitabine was added for viability assessment and IC50 determination. Each experiment was set up in technical triplicates with gemcitabine concentration ranging from 1.0 × 10^-8^ to 1.0 × 10^-4^ mol/L. After 3 days of treatment, cell viability was assessed using CellTiter-Glo as per the manufacturer’s instructions, and relative luminescence units were measured using the TECAN Infinite M200Pro microplate reader. The measured values of luminescence were obtained from three independent experiments, and the data were recorded for analysis. The dose-response model was created using the *drc* package^55^ with a four-parameter log-logistic function and the minimum fixed at 0 and the maximum fixed at the untreated control. The dose-response curve is in Supplementary Figure S1B, and data and R script available at https://github.com/trnkaj/ribosomes.

#### PANC-1 TNTs activation and counting

The human PANC-1 cell line was cultured as described above. For the dose-response curve of TNT numbers vs. gemcitabine concentration we exposed cells to 0.25 µM, 0.5 µM, and 1 µM of gemcitabine for 2 days. The F-actin in live cells was stained using CellMask Green Actin Tracking Stain to detect TNTs in both the control and gemcitabine-treated groups. TMRM was used for staining mitochondria at a final concentration of 100 nM. Hoechst 33342 cell-permeant nuclear counterstain was applied at a final concentration of 5 µg/mL for cell counting purposes. The cells were then incubated with the staining solutions for 30 minutes at 37°C in a 5% CO_2_ incubator.

Four random regions of interest (ROIs) were selected, and scanning was performed using a Plan Fluor 4x objective (0.13 NA) in DAPI, FITC and TRITC field modes. Imaging was conducted with a monochrome pco.panda 4.2 camera mounted on the camera port of a Nikon Eclipse Ts2 fluorescence inverted manual microscope. TNTs in both control and treated cells were counted in four independent images. The number of TNTs at each concentration of gemcitabine was normalised to 1 × 10^4^ cells. For nuclear counting and TNT detection, NIS-AR Elements software was used. Measurements were obtained from three independent experiments, and the data were recorded for analysis.

### Statistical analysis

We used a Bayesian linear model to estimate the dose-dependence of TNT formation on gemcitabine concentration. The model:

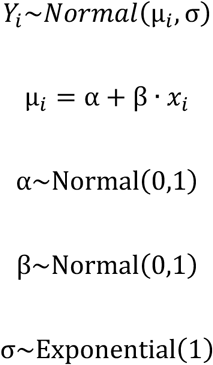

used normalised values of the number of TNTs (Y) and gemcitabine concentration (*x*) and was estimated using the weakly informative priors with STAN via the *rethinking* package by Richard McElreath.^56^ The source data and R code are available at https://github.com/trnkaj/ribosomes

#### Immunocytochemistry of ribosomal components

The PANC-1 cell line was treated for 2 days with 0.5 µM gemcitabine for TNT activation. PDAC primary cells isolated from EUS-FNAB patient samples were grown on a cover glass for immunofluorescence purposes. Patient-derived cells were not exposed to gemcitabine. The samples were fixed in 4% paraformaldehyde (PFA) in phosphate-buffered saline (DPBS) pH 7.4 for 15 minutes at room temperature (RT). The samples were then washed three times in 1% BSA/DPBS, each lasting 5 minutes, and permeabilised for 20 minutes with 1% BSA/DPBS containing 0.1% Triton X-100 at RT. Afterward, cells were washed three times for 5 minutes each in 1% BSA/DPBS and blocked using a solution of 1% BSA/DPBS to prevent non-specific antibody binding. After 1 hour of blocking, primary antibodies were added and diluted in 1% BSA/DPBS. The cells were then incubated with primary antibodies overnight (O/N) in a humidified chamber at 4°C. The following primary antibodies were used: anti-rRNA antibody (Immunogen: whole 5.8S ribosomal RNA), anti-RPS6 antibody, anti-RPS9 antibody, anti-RPL24 antibody, anti-vimentin antibody, and anti-cytokeratin-19 antibody. Subsequently, the samples underwent three 10-minute washes, and were then incubated with the relevant Alexa Fluor 488/546/647 conjugated secondary antibody for 2 hours at RT in the dark. Following the incubation, the samples were rinsed three times with 1% BSA/DPBS for 10 minutes each. ActinGreen™ 488 ReadyProbes or ActinRed™ 555 ReadyProbes were used for F-actin staining. Subsequently, the samples underwent another three washes with 1% BSA/DPBS before being mounted in VECTASHIELD® Antifade Mounting Medium with DAPI.

#### RNA fluorescent in situ hybridization (FISH) – Poly(A) mRNA

Cells with activated TNTs, seeded on cover glass, were fixed for 15 minutes in 4% PFA and permeabilised with 1% Triton X-100 in DPBS. The samples were washed in 900 µL of 1× wash buffer A and incubated O/N at 42°C in a hybridization buffer with 75 nM oligo-d(T) probe and CalFluorRed635. The samples were then washed twice in wash buffer A and twice in 2× SSC. ActinGreen™ 488 ReadyProbes was used for F-actin staining according to the manufacturer’s instructions. Finally, the samples were mounted in VECTASHIELD® Antifade Mounting Medium with DAPI.

#### Negative controls for 5.8S rRNA and for Poly(A) mRNA

RNase treatments were employed as negative controls to validate the specificity of the anti-5.8S rRNA antibody and the FISH Poly(A) assay. These controls were used to confirm the binding affinity of the rRNA monoclonal antibody and the oligo-d(T) probe. PANC-1 cells with activated TNTs were fixed in 4% PFA for 15 minutes at RT, washed twice with DPBS, and incubated with an RNase mix (RNase A, RNase H, and RNase III in 500 µL DPBS) for 2 hours at 37°C. After incubation, the cells were thoroughly washed with DPBS to remove residual RNases. The RNase-treated samples were then stained with immunocytochemistry or RNA FISH. Specificity of the Alexa Fluor™ Plus 647 secondary antibody was assessed using negative controls to differentiate specific binding from non-specific signals. Subtraction of non - specific signals from experimental images enabled accurate quantification of fluorescence corresponding to the binding of primary antibodies to their targets.

#### Cloning of HaloTag-RPS9

For the cloning strategy, total RNA was isolated from MCF-7 breast cancer cells (RRID: CVCL_0031) using the RNeasy Kit according to the manufacturer’s instructions. Two micrograms of total RNA were reverse transcribed by oligo(dT)_18_ in a 20 µL reaction using the Maxima H Minus First Strand cDNA Synthesis Kit following a modified protocol: 50°C for 30 minutes followed by 55°C for 30 minutes. Then, cDNA equivalent to 20 ng of mRNA served as a nested PCR template in a total volume of 50 µL. The first PCR was done with the CLO_hsRPS9_FOR and CLO_hsRPS9_REV pair of primers specific to the 5’ and 3’ untranslated regions of human RPS9. Primers for the second PCR, CLO_hsRPS9Eco_FOR and CLO_hsRPS9Xho_REV were situated to encompass RPS9 open reading frame except the stop codon. Both PCR reactions ran for 35 cycles with Q5 high-fidelity polymerase under similar conditions: initial denaturation at 95°C for 2 minutes, denaturation at 95°C for 20 seconds, annealing 63.8°C for 20 seconds, extension at 72°C for 40 seconds and final extension at 72°C for 3 minutes. The product of the second PCR was cloned via *Eco*RI and *Xho*I sites in frame with the C-terminal HaloTag® to generate pHTC-hsRPS9-HaloTag® CMV-neo vector (Supplementary Figure S8). The insert was sequenced using CMV universal sequencing primer CMV_FOR.

#### Proximity ligation assay (PLA) of 80S ribosome complex, RPS6-RPL24, RPS9-RPL24 and HaloTag-RPS9-RPL24

The PANC-1 cell line with activated TNTs, seeded on cover glass, was fixed for 15 minutes in 4% PFA/DPBS and permeabilised for 20 minutes in 0.1% Triton X-100 in DPBS. The PLA Duolink kit blocking solution was then added to the sample. For the three types of PLA experiments, three different combinations of the primary antibody were used. For the first variant, the cells were incubated with primary antibodies rabbit anti-RPL24 (1:100) and mouse anti-RPS6 (1:150) at 4°C O/N. For the second variant, anti-RPL24 (1:100) and mouse anti-RPS9 (1:150) were used. For the third variant, anti-RPL24 (1:100) and Anti-HaloTag® Monoclonal Antibody (1:250) were used. The samples were washed in DPBS and then in Wash Buffer A. They were incubated with 240 μL of reaction mixture (48 μL PLA probe MINUS stock, 48 μL PLA probe PLUS stock and 144 μL DPBS) in a chamber for 1 hour at 37°C. The cells were washed in 1× Wash Buffer A for 6 × 2 minutes, followed by ligation performed in 240 μL of reaction mixture containing 6 μL of ligase, 48 μL of ligation buffer, and 186 μL of molecular H_2_O. Samples were incubated in the ligation reaction mixture for 30 minutes at 37°C and then washed 6 × 2 minutes in Wash Buffer A. Next, 240 μL of amplification reaction mixture (3 μL polymerase and 48 μL amplification solution) was added to each sample before incubation at 37°C for 100 minutes. The samples were washed in 500 μL of Wash Buffer B for 3 × 5 minutes and 1 × 2 minutes in 0.01× Wash Buffer B. ActinRed™ 555 ReadyProbes was used for F-actin staining according to the manufacturer’s instructions. Finally, the samples were mounted using VECTASHIELD® Mounting Medium with DAPI. For the PLA assays, three combinations were used: one detecting the interaction between RPL24 and RPS6, the second detecting the interaction between RPL24 and RPS9, and the third aimed at detecting the interaction between RPL24 and HaloTag-RPS9 expressed from the pHTC-hsRPS9-HaloTag® CMV-neo vector. As a negative control for the PLA assay, PANC-1 cells with activated TNTs were treated with puromycin at a final concentration of 10 µg/mL for 4 hours to cause ribosome disassembly. After treatment, the cells were fixed in 4% PFA/DPBS for 15 minutes at RT, washed twice with DPBS, and the PLA experiment using primary antibodies anti-RPS6 and anti-RPL24 was performed.

#### Transfection of HaloTag-RPS9

The PANC-1 cell line was cultured in complete DMEM medium at 37°C in a humidified atmosphere with 5% CO₂. Upon reaching approximately 70% confluence in a 24-well plate, the cells were transfected with the pHTC-hsRPS9-HaloTag® CMV-neo vector (donor cells) using Lipofectamine™ 3000 Transfection Reagent, following the manufacturer’s instructions. Two days post-transfection, the transfection reagent was removed by washing the cells three times with DPBS, and the medium was replaced with fresh complete DMEM medium.

### Transport of HaloTag-RPS9 to acceptor cells

Two stained variants of PANC-1 acceptor cells were used: one with the nucleus stained using the CellLight™ Nucleus-GFP BacMam 2.0, and the other with the entire cell stained using the covalent CellTrace™ Violet Cell Proliferation Kit, both following the manufacturer’s instructions. Expressed RPS9 (in donor cells) was visualised by labelling live cells with the HaloTag-TMR ligand for 30 minutes in an incubator at 5% CO₂ before the co-culture experiment. The wells containing donor and acceptor cells were washed with pre-warmed DPBS at 37°C prior to trypsinisation and centrifugation. Subsequently, the centrifuged cells were washed twice separately with pre-warmed DPBS and resuspended in complete DMEM medium. Donor and acceptor cells were then mixed in equal proportions and seeded into a 24-well plate. One day post-seeding, the cells were treated with 0.5 µM gemcitabine to activate TNTs for two days. After the three-day co-culture, expressed RPS9 was visualised by re-labelling live cells with the HaloTag-TMR ligand and imaged using a fluorescence microscope. For live-cell imaging of HaloTag-RPS9 transport, cells expressing RPS9 were seeded onto ibidi µ-Dishes with a coverslip bottom and treated with 0.5 µM gemcitabine one day post-seeding to induce TNT formation over two days. Following TNT induction, live cells were labelled with HaloTag-TMR ligand to detect RPS9 expression from the vector, actin was visualised using CellMask™ Green Actin Tracking Stain, and nuclei were counterstained with Hoechst 33342. Live-cell videos were acquired using a Leica Stellaris 8 confocal microscope.

### Incorporation of HaloTag-RPS9 into ribosomes of acceptor cells

To demonstrate the incorporation of HaloTag-RPS9 into the ribosomes of acceptor cells, acceptor cells stained with CellTrace™ Violet were co-cultured with non-stained donor cells, transfected with the pHTC-hsRPS9-HaloTag vector, as mentioned in the methods above. One day post-seeding, the cells were treated with 0.5 µM gemcitabine to activate TNTs for two days. After the three-day co-culture, cells were labelled with the HaloTag-TMR ligand and fixed in 4% PFA/DPBS for 15 minutes at RT before undergoing a proximity ligation assay (PLA) using Anti-HaloTag® Monoclonal Antibody and anti-RPL24 primary antibodies.

The PLA (anti-HaloTag and anti-RPL24) staining and TMR ligand labelling were also performed on cells lacking the HaloTag-RPS9 transgene, co-cultured with acceptor CellTrace™ Violet-stained cells, which served as the negative control. Actin was stained using CellMask™ Deep Red Actin. VECTASHIELD® Antifade Mounting Medium without DAPI was used for mounting, and the cells were subsequently imaged using a confocal microscope.

#### Flow cytometry for the dual cell labelling experiment

For the dual cell labelling experiment, PANC-1 cells were cultured to 70% confluency and then transfected with the HaloTag-RPS9 plasmid using Lipofectamine™ 3000 Transfection Reagent. 48 hours post-transfection, cells expressing HaloTag-RPS9 were labelled with TMR ligand to verify transfection efficiency. Before mixing donor and acceptor cells, donor cells (HaloTag-RPS9) and acceptor cells (CellTrace Violet) were washed with pre-warmed DPBS, centrifuged, washed twice separately with pre-warmed DPBS and resuspended in complete DMEM medium. Cells were combined and seeded into a 24-well plate at equal numbers. Two conditions were set up for the mixed culture: an untreated control and a group treated with 0.5 µM gemcitabine for 24 hours. Following incubation, cells expressing HaloTag-RPS9 were labelled with TMR ligand for 30 minutes at 37°C to ensure accurate signal detection. All cells were then fixed in the dark with 4% PFA/DPBS for 15 minutes at RT, centrifuged, washed twice with DPBS, and resuspended in 500 µl of DPBS for flow cytometry.

Cytometry data were analysed using an R script available at https://github.com/trnkaj/ribosomes. Data. fcs files generated by BD FACSuite™ Software (BD Biosciences, CA, USA) were read using the *flowCore* and *ggcyto* packages by Bioconductor.^57,58^ All measured events for each experimental condition were gated using FSC-W values to remove doublets, which could present themselves as double-stained cells. Minimum fluorescence intensity values for selecting HaloTag TMR and CellTrace Violet positive events were chosen using fluorescence values of fluorescence minus one (FMO) control cells. Specifically, for the HaloTag/CellTrace Violet mixed population experiments the DAPI channel signal of cells containing HaloTag-RPS9 with TMR ligand was used to set the CellTrace Violet positive gate as any value over the maximum DAPI channel value in these control cells, and PE channel fluorescence of cells stained with CellTrace Violet was used to set the HaloTag TMR positive gate as any value over the maximum PE channel value in these control cells. Events in FMO controls with fluorescence intensity over ten standard deviations above the log-mean were ignored in the setting of fluorescence intensity gates. As double-positive cells we considered cells with fluorescence values in both channels higher than the respective fluorescence minus one control. Since the doublet gate (FSC-W) is likely to have a substantial effect on the total number of double-stained cells observed, we performed a sensitivity analysis by varying the value of this gate over the range of available events and observing the changes in the resulting number of double-positive cells (Supplementary Figure S4F).

#### Preparation of cells for western blotting, flow cytometric analysis of global translation, and microscopic quantification of ribosomes

PANC-1 cells were seeded in 24-well plates at a density of 5 × 10^4^ cells per well. On the following day, cells were transfected with 9 pmol Silencer Select negative-control siRNA or with 9 pmol siRNA directed against RPS6 and 9 pmol siRNA directed against RPL24 (all Thermo Fisher Scientific) using Lipofectamine™ 2000 (Thermo Fisher Scientific) in accordance with the manufacturer’s instructions. Transfection was allowed to proceed for 24, 48, or 72 hours for Western blot experiments, and for 48 hours for flow cytometry-based global translation and ribosome quantification experiments. Transfection efficiency was assessed with Silencer™ Cy3-stained negative-control siRNA (Supplementary figure S9).

### Western blotting

On the first, second, and third day following transfection with either negative control siRNA or siRNA targeting RPS6 +RPL24, cells were harvested on ice using a cell scraper and resuspended in ice-cold DPBS supplemented with protease inhibitors (cOmplete™ Mini, EDTA-free, Roche). The cell suspension was centrifuged at 600 × g for 5 minutes at 4°C. The supernatant was discarded, and the resulting pellet was resuspended in ice-cold RIPA buffer containing protease inhibitors. Cell lysates were incubated overnight at −80°C to ensure complete lysis. The lysates were subsequently centrifuged at 15,000 rpm for 15 minutes at 4°C to remove cellular debris. The supernatant, containing soluble proteins, was carefully transferred to new low protein-binding tubes (Eppendorf). Protein concentrations were determined using the BCA protein assay (Thermo Fisher Scientific), according to the manufacturer’s instructions. **SDS-PAGE and Immunoblotting** - protein samples were mixed with 2× Laemmli buffer, preheated at 95°C for 5 minutes, and loaded onto a 12% SDS– polyacrylamide gel (according to Hoefer manuals) at a final concentration of 15 µg protein per lane. After electrophoretic separation, proteins were transferred onto PVDF membranes (Cytiva Amersham™ Protran) using Towbin transfer buffer. Total protein staining was performed using the No-Stain™ Protein Labelling Reagent (Thermo Fisher Scientific) for normalisation purposes. Membranes were imaged using the Alliance Q9-series imaging system (Uvitec). **Blocking and Antibody Incubation** - membranes were blocked in 3% non-fat dried milk dissolved in TTBS (Tris-buffered saline containing 0.1% Tween-20) for 1 hour at RT. Subsequently, membranes were incubated overnight at 4°C under mild agitation with primary antibodies against RPS6, RPL24, and vinculin (loading control), each diluted in 3% non-fat dried milk in TTBS. Following primary incubation, membranes were washed three times with 1% milk in TTBS (20 minutes per wash) and then incubated for 2 hours at RT with HRP-conjugated secondary antibodies (Promega) diluted in the same buffer. A further three washes of 20 minutes each were performed. **Signal Detection and Quantification** - protein bands were visualised using enhanced chemiluminescence (ECL) reagents. Chemiluminescent signals were detected using the Alliance Q9-series imaging system and quantified using Bio-Rad Image Lab software. Densitometric analysis was used to assess changes in protein expression levels following siRNA treatment, with normalisation to total protein.

### Flow cytometric analysis of global translation, and microscopic quantification of ribosomes (Table M1)

48 hours after siRNA transfection, cells were washed with pre-warmed DPBS. Following trypsinisation, trypsin activity was neutralised by adding complete DMEM medium. The cell suspension was then centrifuged at 600 × g for 5 minutes. The siRNA-negative control cells (or siRNA-RPS6 + RPL24 treated cells in the colour-reversed experimental setup) were resuspended in serum-free DMEM and stained with the cell proliferation dye CFSE (1 µL/mL; Thermo Fisher Scientific) for 30 minutes at 37 °C in a dark incubator. After staining, the cells were centrifuged at 600 × g for 5 minutes at RT and washed twice with 10 mL of complete DMEM medium containing FBS to remove any residual staining solution, thereby preventing its binding following mixing with non-stained siRNA RPS6+RPL24 treated cells (or siRNA-negative control cells in the reverse-colour experimental setup). The cells were resuspended in complete DMEM and seeded into the following experimental groups, which were labelled for methodological clarity: **control cells** – siRNA-negative control PANC-1 cells; **siRNA cells** – PANC-1 cells co-transfected with siRNA-RPS6 and siRNA-RPL24; **mixed control & siRNA cells** – a 1:1 mixture of siRNA-negative control PANC-1 cells and PANC-1 cells co-transfected with siRNA-RPS6 and siRNA-RPL24; **NTC** – non-transfected, non-stained negative control cells. All cells were subsequently treated with 0.5 µM gemcitabine for 24 hours to induce tunnelling nanotube formation.

### Click-iT global translation assay (CFSE, HPG-Azide) for flow cytometry experiment

24 hours after gemcitabine treatment, cells were washed, and 600 µL of DMEM without methionine, supplemented with 10% foetal bovine serum dialyzed, 2 mM GlutaMAX, 1% Penicillin/Streptomycin, 1mM Sodium Pyruvate and 200 µM freshly prepared L-Cysteine (complete DMEM without methionine medium) was added for 15 minutes to allow cell equilibration at 37°C in a 5% CO_2_ incubator in the dark. After equilibration, fresh complete DMEM medium without methionine, containing HPG (Thermo Fisher Scientific) 50 µM at final concentration, was added to experimental groups control cells, siRNA cells, and mixed control & siRNA cells mentioned above. Following 5 hours of HPG treatment, cells were washed with pre-warmed DPBS and trypsinised. After adding complete DMEM to inhibit trypsin, the cells were gently harvested and centrifuged at 600 × g for 5 minutes. Cells in suspension were fixed in 4% PFA/DPBS for 15 minutes at RT in the dark, followed by centrifugation at 600 × g for 5 minutes. The cell pellet was subsequently washed with DPBS. For permeabilisation, cells were treated with 0.002% Triton X-100/DPBS for 5 minutes at RT in the dark, then centrifuged and washed with DPBS before the Click-iT reaction. Cells were resuspended and incubated in a freshly prepared Click-iT reaction mix (1 µM Azide 647 from Vector Laboratories, 10 mM sodium L-ascorbate, and 2 mM CuSO₄ in DPBS) for 30 minutes at RT in the dark. After the reaction, cells were centrifuged, and the pellet was resuspended in a DPBS, followed by a second centrifugation. Finally, the pellet was resuspended in 500 µL of DPBS and stored at 4°C in a dark tube for cytometric analysis. For flow cytometric analysis, NTC-non-stained negative control was first used to define the population of non-fluorescent cells. Permeabilised negative controls stained with Azide 647 (Vector Laboratories) were included to assess and adjust for background fluorescence. Laser excitation at 488 nm detected CFSE stained siRNA negative control cells, while laser excitation at 647 nm detected the translation signal from Click-iT/Azide 647. These experiments were performed in six replicates.

Gates for flow cytometry analysis were set in the following way: cell debris and doublets were excluded using the FSC-A vs FSC-H plot. Events with FITC-H of ≤ 0 were also removed. For the one-gate setting events we classified as CFSE+ (green) if their FITC-H values exceeded the geometric mean + 5×S.D. of the unstained control cells. In the stricter two-gate setting events with FITC-H values less then the geometric mean + 2×S.D. of the unstained control cells were considered CFSE-(non-green) cells, and cells with FITC-H values more than the geometric mean - 2×S.D. of the unmixed CFSE-stained cells were considered as CFSE+ (green). The two-gate setting was designed to exclude partially stained cells formed due to the putative transfer of CFSE between stained and unstained cells.

### Statistical analysis

Data were analysed using R. Relevant data were extracted from flow cytometry .fcs files using the *flowCore* package and analysed using a Bayesian varying-intercepts linear model^59^:

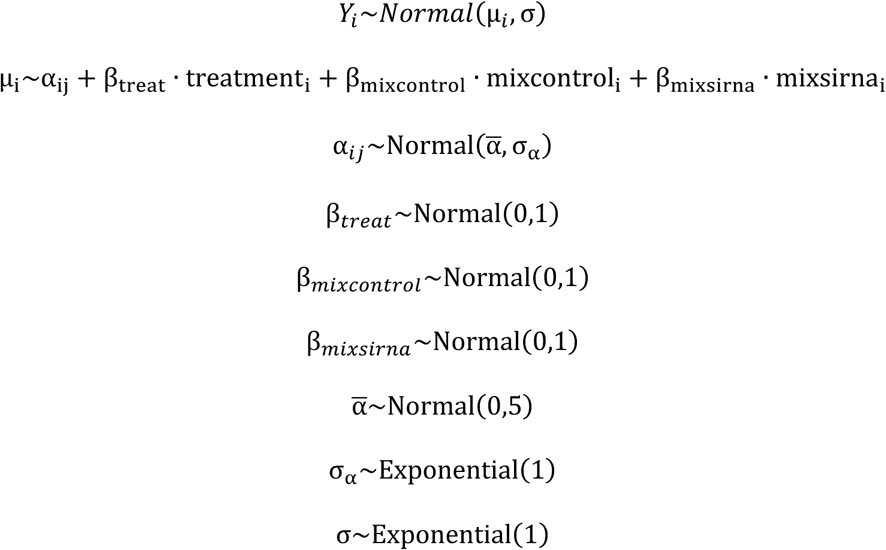

where Y_i_ are standardised, log-transformed fluorescence intensities at 647 nm, α_ij_ is the varying intercept indexed by experiment and corresponding to the fluorescence of control cells, treatment is a binary indicator of treatment with RPS6+RPL24 siRNA (0 for non-treated, 1 for treated), β_treat_ is the coefficient of siRNA treatment effect, mixcontrol is a binary indicator of control cells in mixed culture with siRNA treated cells (0 for non-mixed control and siRNA cells and for mixed siRNA cells, 1 for control cells in mixed culture), β_mixcontrol_ is the coefficient of the effect of mixing on control cells, mixsirna is a binary indicator of siRNA-treated cells in mixed culture with control cells (0 for non-mixed control and siRNA cells and for mixed control cells, 1 for siRNA-treated cells in mixed culture), β_mixsirna_ is the coefficient of the effect of mixing on siRNA-treated cells, alpha bar is the hyperprior for α_ij_. After estimation with STAN using the *rethinking* package in R, samples were drawn from the posterior distribution, de-standardised, exponentiated and used to predict fluorescence intensity distributions for average populations of non-treated, treated, non-mixed and mixed cells, and for differences between mixed and non-mixed cells to show the average effect of mixing. The source data and R code are available at https://github.com/trnkaj/ribosomes.

**Table M1.**
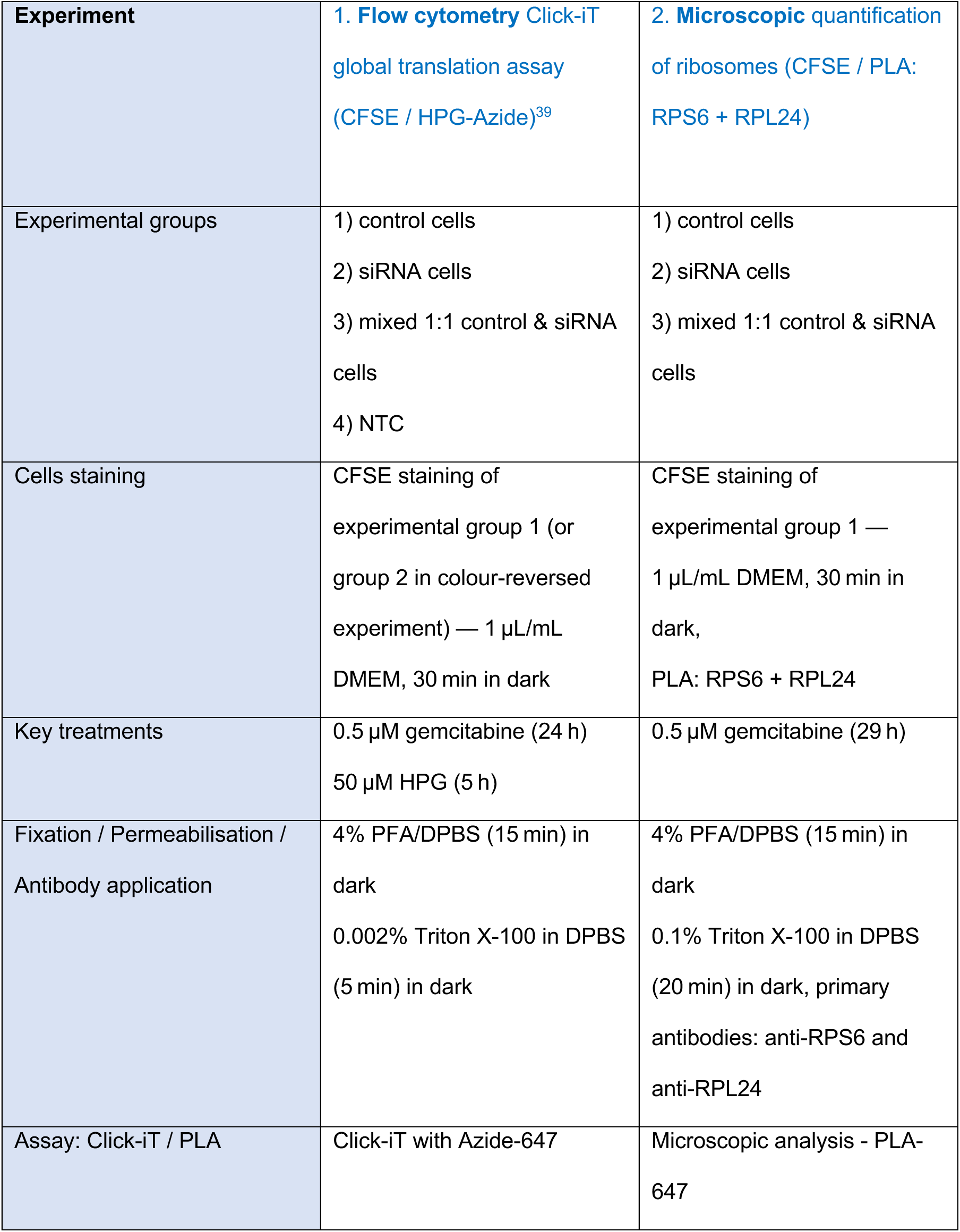
Flow cytometry for global translation and microscopic quantification of ribosomes. Table M1 summarise methodology workflow for Flow cytometry Click-iT global translation assay and for microscopic quantification of ribosomes. Experimental groups are **control cells** – siRNA-negative control PANC-1 cells; **siRNA cells** – PANC-1 cells co-transfected with siRNA-RPS6 + siRNA-RPL24; **mixed control & siRNA cells** – a 1:1 mixture of siRNA-negative control PANC-1 cells and PANC-1 cells co-transfected with siRNA-RPS6 + siRNA-RPL24; **NTC** – non-transfected, non-stained negative control cells.

### Microscopic quantification of ribosomes

Cells in experimental groups control cells, siRNA cells, and mixed control & siRNA cells were seeded on cover glasses for immunofluorescence. Twenty-four hours after gemcitabine treatment, the cells were washed with pre-warmed DPBS and fixed in 4% PFA/DPBS for 15 minutes at RT in the dark. Cells were permeabilised for 20 minutes in 0.1% Triton X-100 in DPBS at RT in the dark, then washed with DPBS. For immunofluorescence-based ribosome quantification, the Duolink PLA 647 Kit was used, following the previously described protocol. Primary antibodies against RPS6 (1:150) and RPL24 (1:100) were applied for ribosome detection. Samples were mounted in VECTASHIELD® Antifade Mounting Medium with DAPI. CFSE stained siRNA-negative control cells were detected using 488 nm laser excitation, while assembled ribosomes were detected using 647 nm laser excitation. ActinRed 555 ReadyProbes was used for F-actin staining. Microscopy scanning was performed using a Leica Stellaris 8 confocal microscope with a 40× objective and zoom factor of 1.08×. A total of 13 z-stack slices were acquired, and their maximum intensity projection was subsequently analysed.

Ribosome quantification was performed using NIS-Elements AR software, based on the detection of ribosomal PLA spots within individual cells. ROIs, corresponding to the entire visible area of each cell, were defined either automatically or manually using the polygonal ROI tool. Actin-stained scans were used to ensure accurate delineation of cell boundaries.

Only cells that were fully captured in the field of view and did not overlap with neighbouring cells were included in the analysis, in order to ensure reliable per-cell quantification. Cells located at the image edges or visibly overlapping with adjacent cells were excluded from the analysis.

ROIs were colour-coded for improved clarity—siRNA-RPS6 + siRNA-RPL24 treated cells (blue), siRNA-negative control cells (green). ROIs were copied onto the corresponding scans with ribosomes. Spot detection was performed using fixed parameters: a spot diameter of 4px for 1.08× zoom with a contrast threshold consistently set at 9. These parameters were maintained throughout the entire experiment to minimise bias and ensure methodological consistency. This standardised approach also ensured uniform error characteristics across all analysed images. Spots were quantified using the measurement module in NIS-Elements, and exported to MS Excel. A total of 17 scans (No. of cells 412) from the siRNA-negative control cells, 22 scans (No. of cells 412) from the siRNA-RPS6 + siRNA-RPL24 treated cells and 17 scans (No. of siRNA-negative control cells 215/ No. of siRNA-RPS6 + siRNA-RPL24 treated cells 170) from the co-cultures experimental groups were analysed.

### Statistical analysis

Ribosome counts were analysed using a linear Bayesian model:

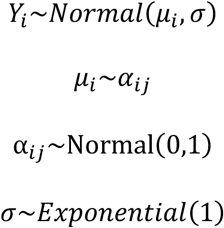

Where Y_i_ are standardised ribosome counts for each cell, α_ij_ are average counts indexed by treatment (1 for non-mixed control cells, 2 for non-mixed siRNA-treated cells, 3 for mixed control cells and 4 for mixed siRNA-treated cells). After estimation with STAN using the *rethinking* package in R, samples were drawn from the posterior distribution, de-standardised and used to predict ribosome count distributions for average populations of non-mixed and mixed control and siRNA cells, and for differences between mixed and non-mixed cells to show the average effect of mixing. The source data and R code are available at https://github.com/trnkaj/ribosomes

#### Live cell microscopy transfer of HaloTag-RPS9 protein via TNTs

For live-cell microscopy, cells transfected with the pHTC-hsRPS9-HaloTag® CMV-neo vector were labelled with HaloTag-TMR ligand to visualise RPS9 expression. Control cells were stained with the CellTrace™ CFSE Cell Proliferation Kit, as described in the Methods section below, mixed in a 1:1 ratio with HaloTag-RPS9 transfected cells, and treated with 0.5 µM gemcitabine for 2 days to induce tunnelling nanotubes. A live-cell video capturing the transport of HaloTag-RPS9 (red) within TNTs formed between PANC-1 cells was recorded using a Leica Stellaris 8 confocal microscope (3.873 seconds per frame). Nuclei are stained by Hoechst (blue) and actin filaments are stained using the CellMask Green Actin Tracking Stain (green).

#### Fluorescence and confocal microscopy

For scanning fluorescently stained samples, a Leica Stellaris 8 confocal microscope (indicated by * in the respective images) and a Nikon Eclipse Ts2 fluorescence inverted manual microscope (indicated by # in the respective images) were used. An inverted routine fluorescence microscope (Nikon Eclipse Ts2) was equipped with Plan Fluor 4x objective (0.13 NA), Plan Fluor 10x objective (0.30 NA), S Plan Fluor ELWD 20x objective (0.45 NA) and S Plan Fluor ELWD 40x objective (0.60 NA). Next, an inverted confocal microscope (Leica Stellaris 8) with a Glycine 40x objective (1.25 NA) was used for sample visualization. The inverted microscope, equipped with a controlled chamber system (Temp Controller 2000–2, Pecon and a CO_2_ Controller, Pecon), was used for time-lapse recordings. The images were processed using NIS-Elements Advanced Research, Leica Application Suite (LAS X), and ImageJ/FIJI.

#### Graphical abstract and experimental scheme

For the drawing of the graphical abstract and the experimental scheme in Figure 5 we used pictures from NIH Bioart^60^ with some modifications, and from Servier Medical Art (https://smart.servier.com/), licensed under CC BY 4.0 (https://creativecommons.org/licenses/by/4.0/) with some modifications.

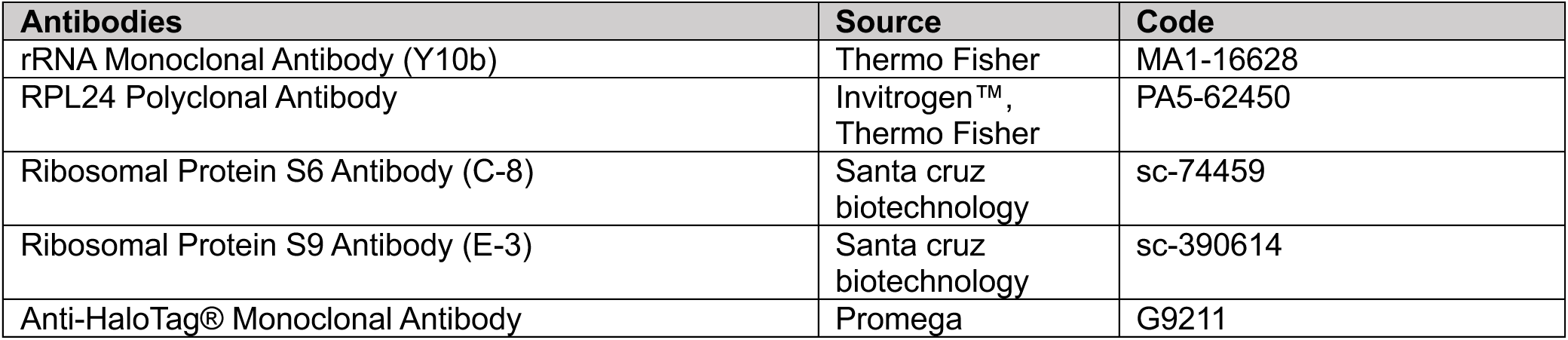
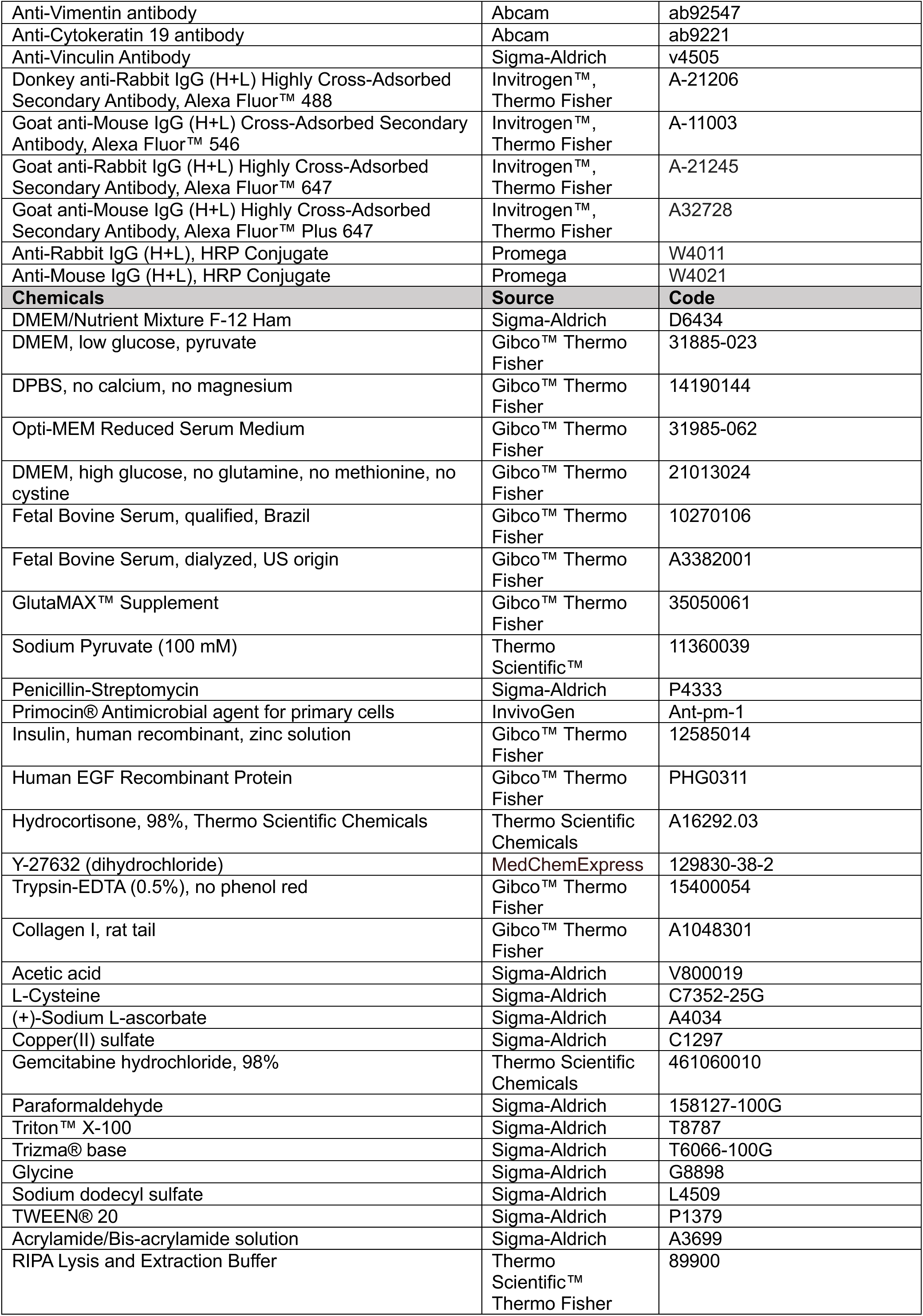
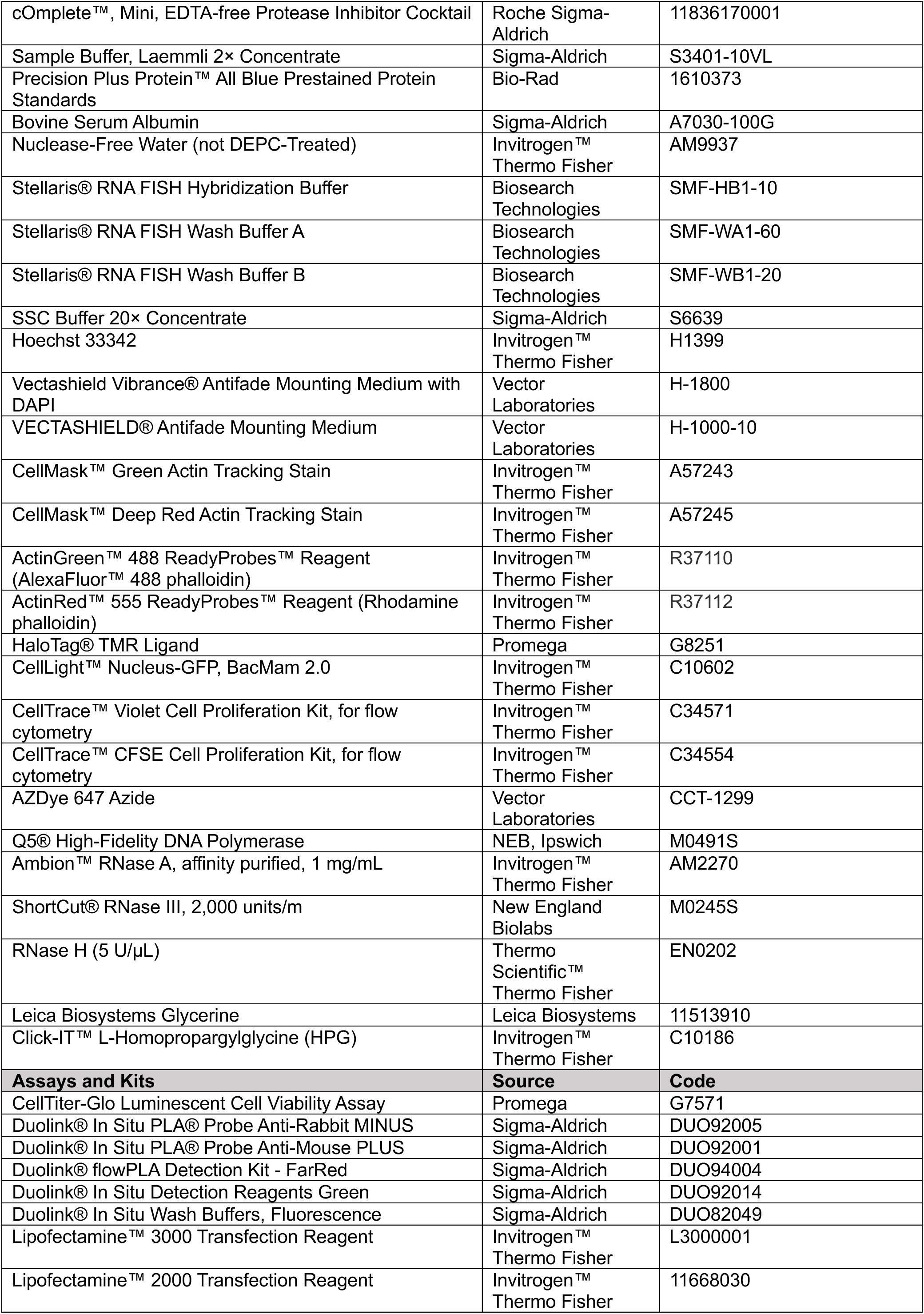
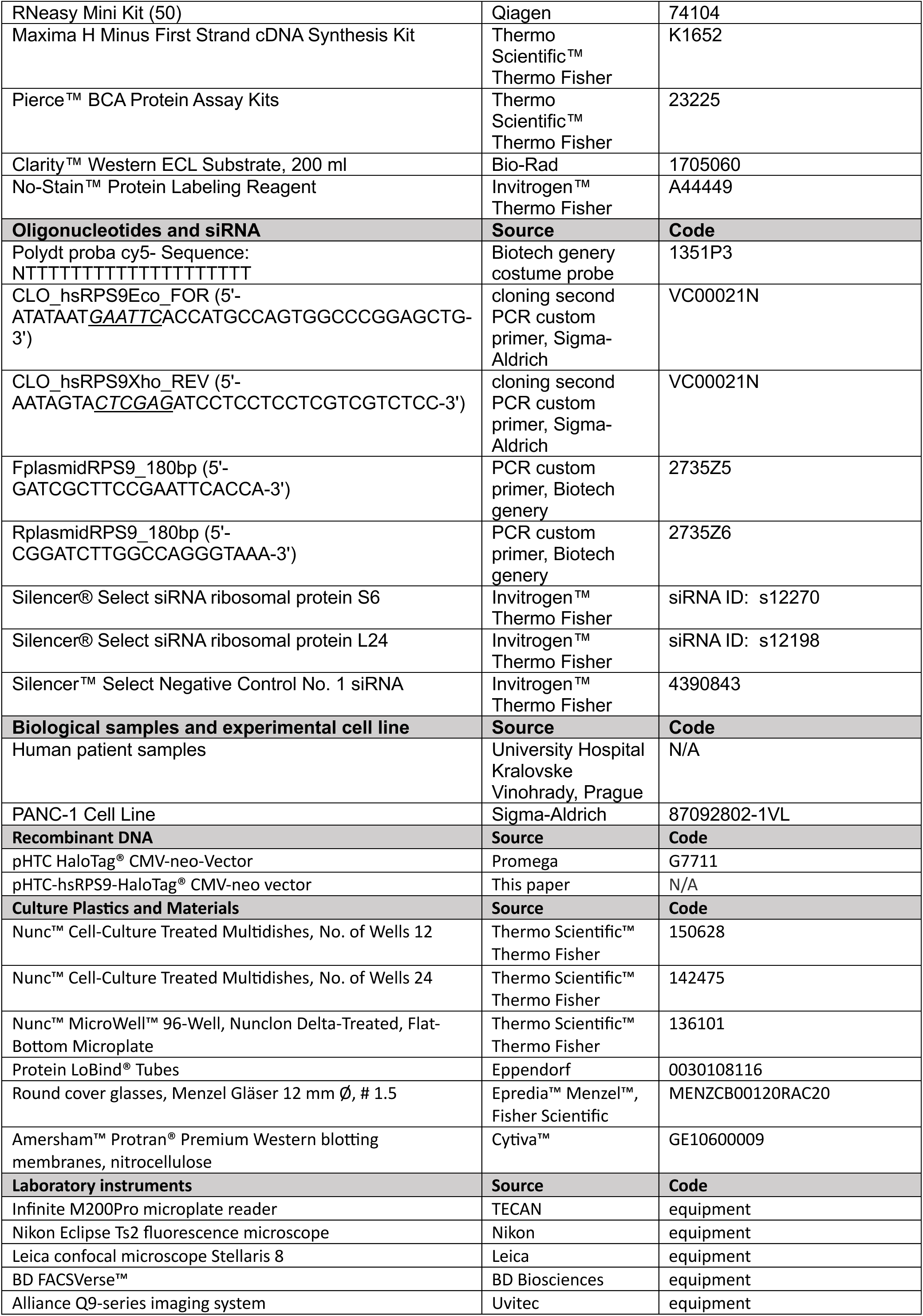
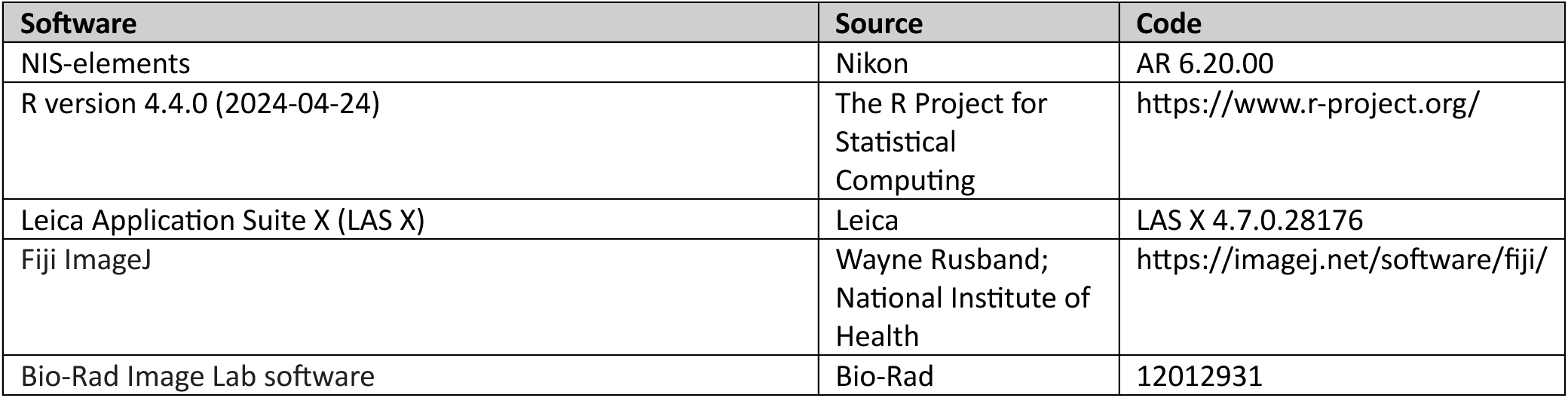

