## Supplemental figures for "Ribosome transfer via tunnelling nanotubes rescues protein synthesis in pancreatic cancer cells"

### TABLE S1.

**Table S1. Summary of patient data and clinical outcomes.** This table presents anonymised patient identifiers, demographic information, detailed diagnoses, and treatment plans for patients diagnosed with pancreatic ductal adenocarcinoma, summarising the patients used in our study. The diagnoses were confirmed by a diagnostic pathologist using imaging methods such as CT scans and EUS-guided FNAB. Treatment modalities listed include palliative chemotherapy, neoadjuvant therapy, and one instance of non-attendance at treatment.

| Patient ID anonymised for study purposes | Gender | Age | Diagnosis determined by diagnostic pathologist and imaging methods (CT,EUS) | Treatment determined by oncologist |
|---|---|---|---|---|
| BJPN15 | M | 60 | Invasive adenocarcinoma of the ductal type in the head of pancreas | Palliative chemotherapy |
| BJPN18 | M | 74 | Moderately differentiated ductal adenocarcinoma in head of the pancreas | Palliative chemotherapy |
| BJPN19 | M | 72 | Invasive, poorly differentiated adenocarcinoma of the pancreatobiliary type in body of the pancreas | Palliative chemotherapy |
| BJPN24 | F | 43 | Tumour lesion visible in the head of the pancreas on a CT scan. The biopsy, was labelled non-diagnostic by the examining pathologist due to the presence of blood coagula, small connective tissue fragments, and sparse sections of duodenal smooth muscle with ganglionic cells, no pancreatic structures were identified in the sample | Patient failed to attend treatment |
| BJPN26 | M | 71 | Ductal adenocarcinoma in the head of the pancreas | Neoadjuvant therapy |

### FIGURE S1A

**Figure S1A.** Cells in 2D cultures of fine-needle biopsy samples from a PDAC patients develop TNTs.

#### Patient sample BJPN18 #

Cells in 2D cultures of fine-needle biopsy samples from a PDAC patient BJPN18 develop TNTs (Day 2).

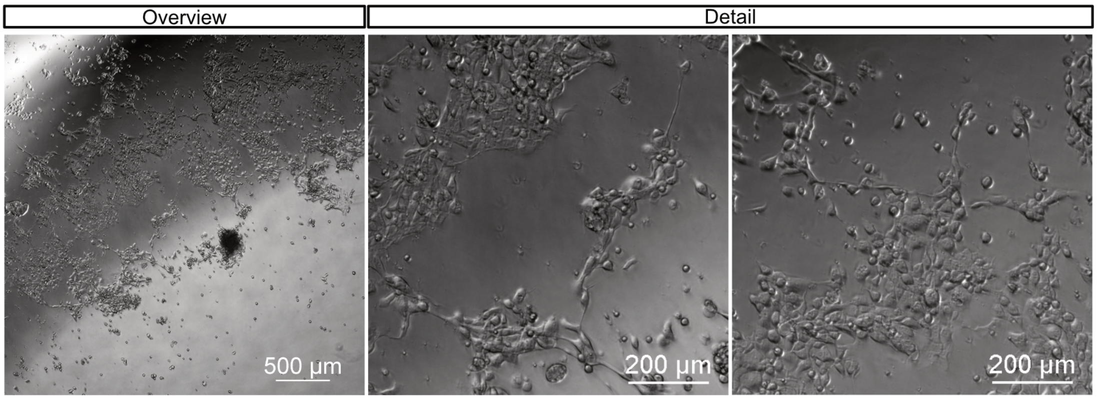


#### Patient sample BJPN19 #

Cells in 2D cultures of fine-needle biopsy samples from a PDAC patient BJPN19 develop TNTs (Day 15)

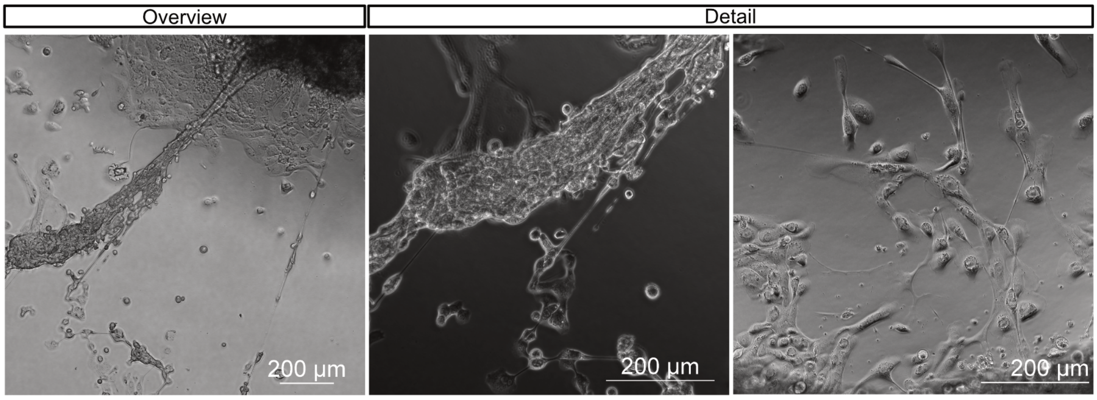


#### Patient sample BJPN26 #

Cells in 2D cultures of fine-needle biopsy samples from a PDAC patient BJPN26 develop TNTs (Day 7)

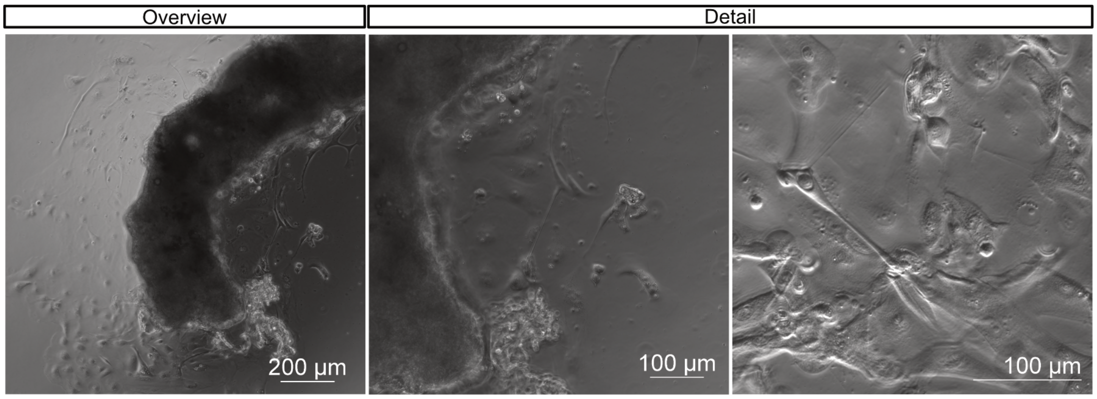

### FIGURE S1B

**Figure S1B.** The dose-response curve: IC50 for gemcitabine in PANC-1 cells is 1.6 μM.

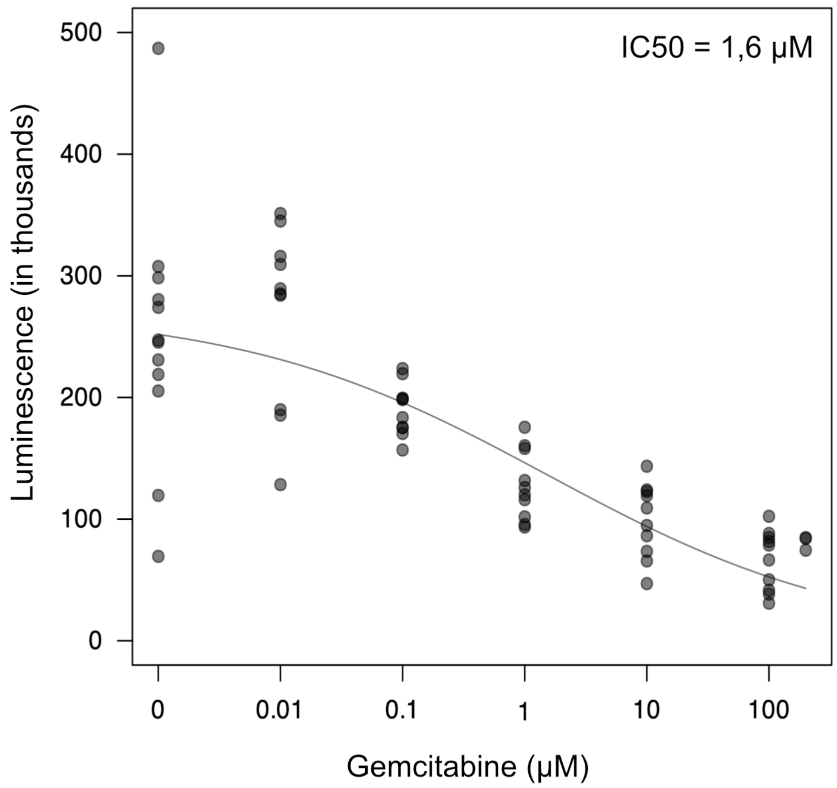


### FIGURE S1C

**Figure S1C.** Posterior distribution of slope in linear model of dose-dependent TNT formation

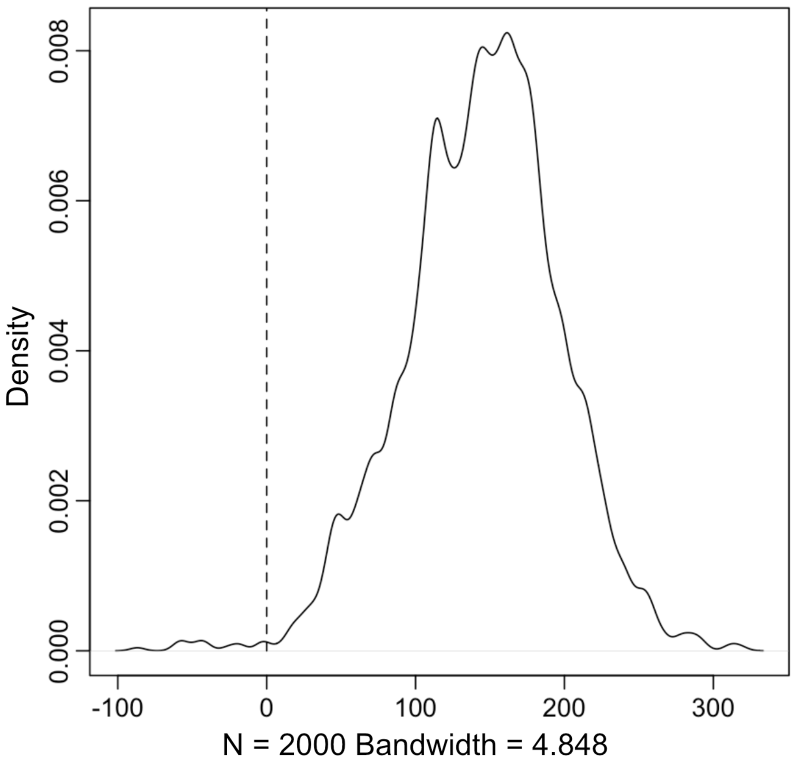

### FIGURE S2A

**Figure S2A.** Ribosomal 5.8S rRNA detected in tunnelling nanotubes (a, b, c).

**a.** PANC-1 5.8S rRNA *

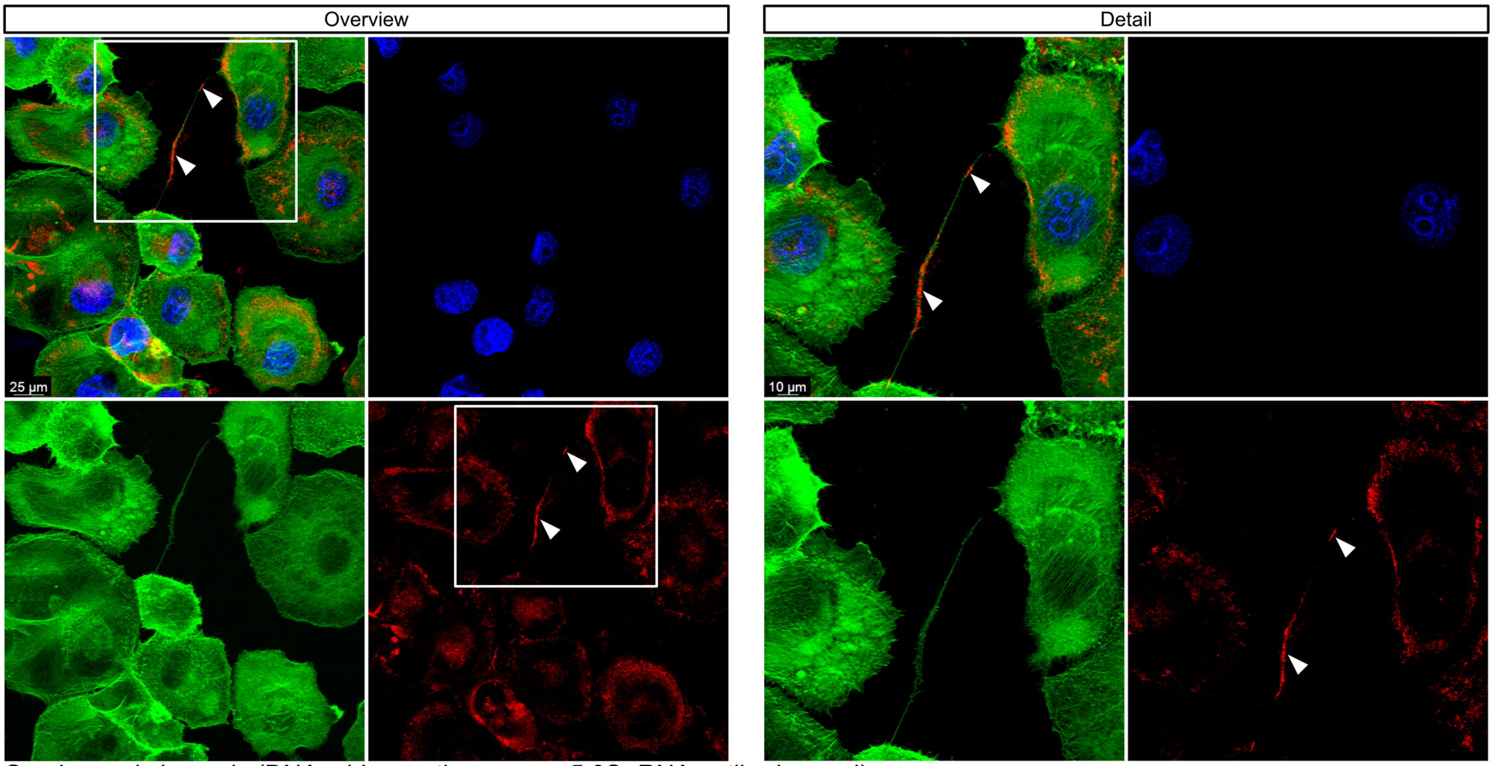


Overlay and channels (DNA – blue, actin – green, 5.8S rRNA antibody – red)

**b.** PANC-1 5.8S rRNA #

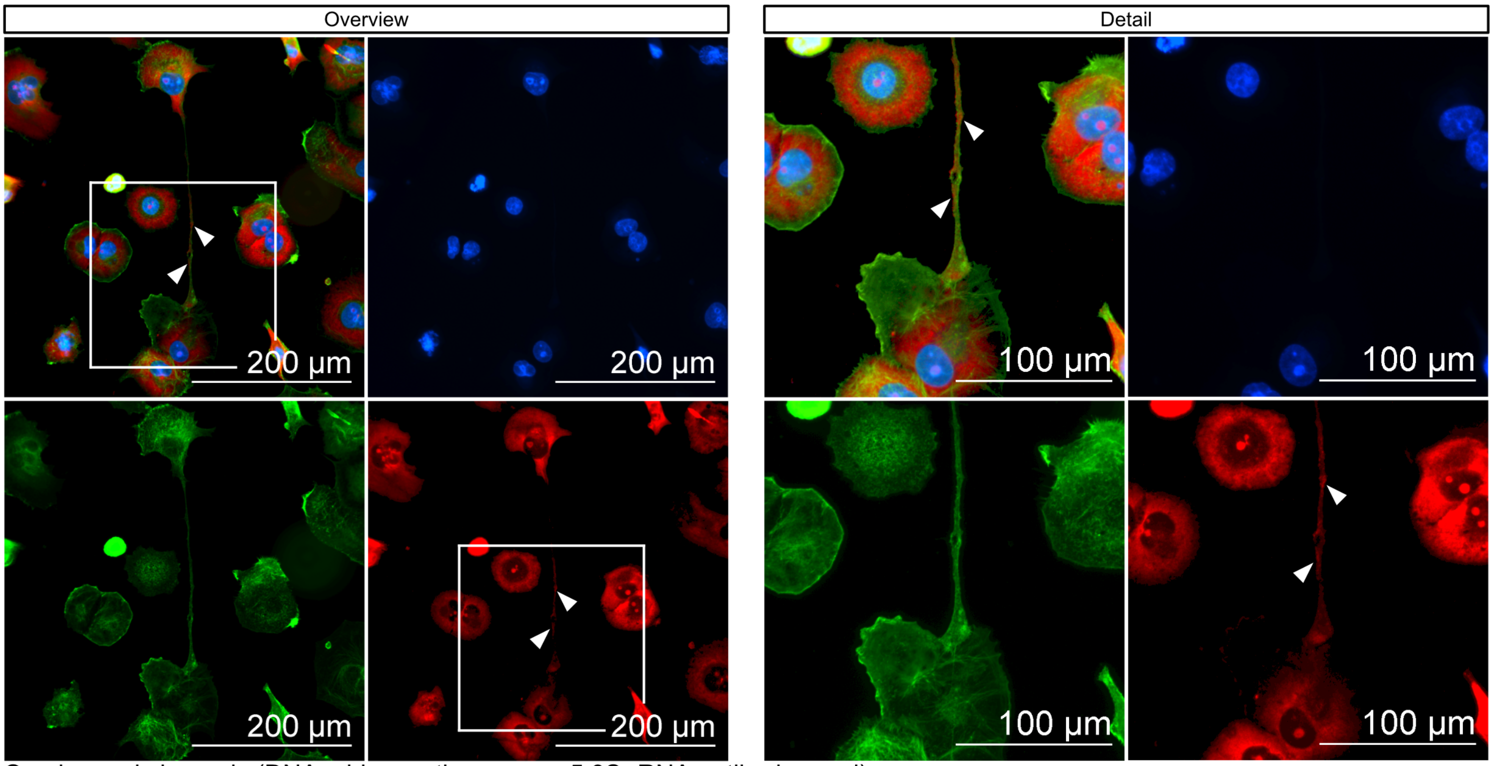


Overlay and channels (DNA – blue, actin – green, 5.8S rRNA antibody – red)

### FIGURE S2A cont.

**Figure S2A.** Ribosomal 5.8S rRNA detected in tunnelling nanotubes (a, b, c).

**c.** PANC-1 5.8S rRNA *

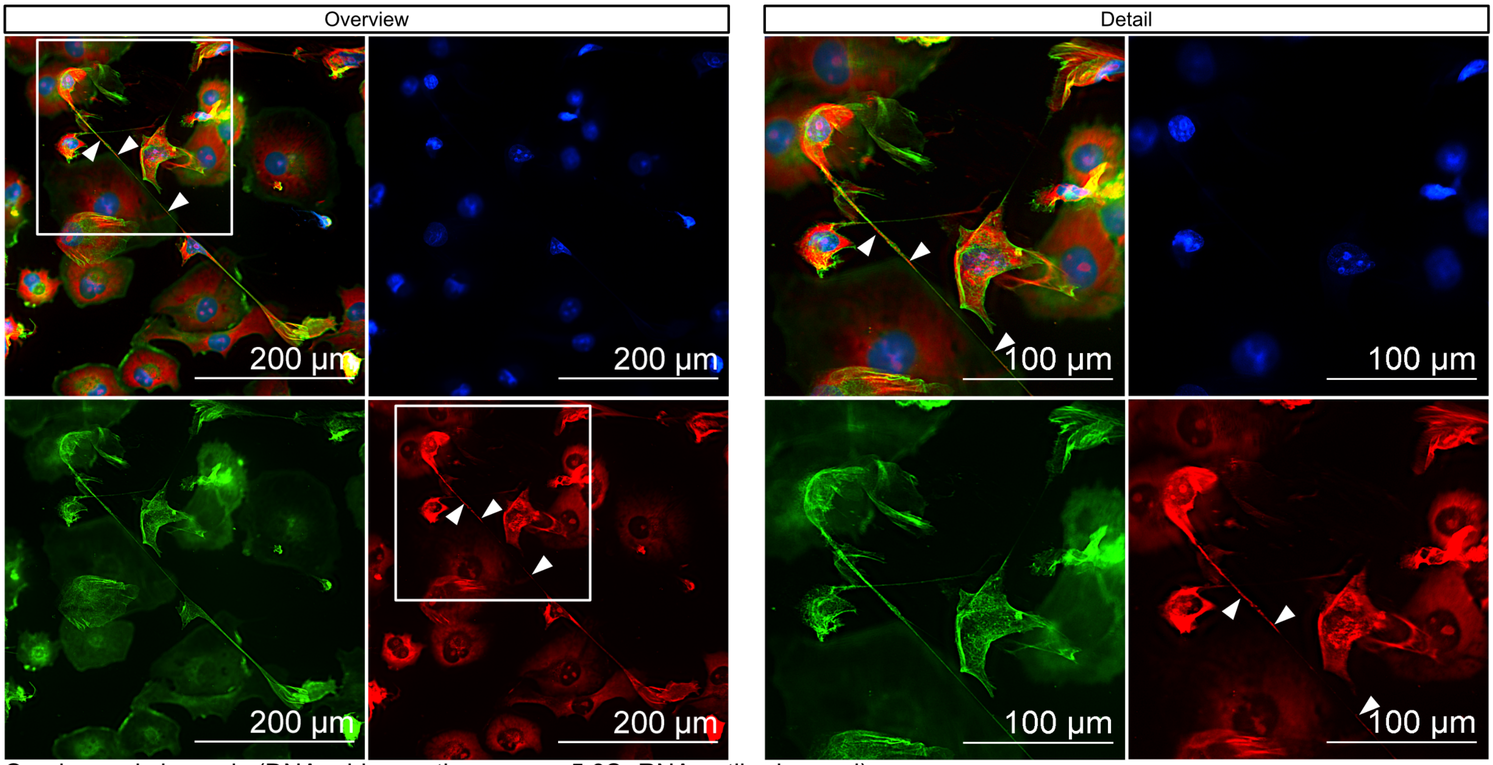


Overlay and channels (DNA – blue, actin – green, 5.8S rRNA antibody – red)

### FIGURE S2B

**Figure S2B.** Ribosomal protein S6 detected in tunnelling nanotubes (a, b, c).

**a.** PANC-1 RPS6 *

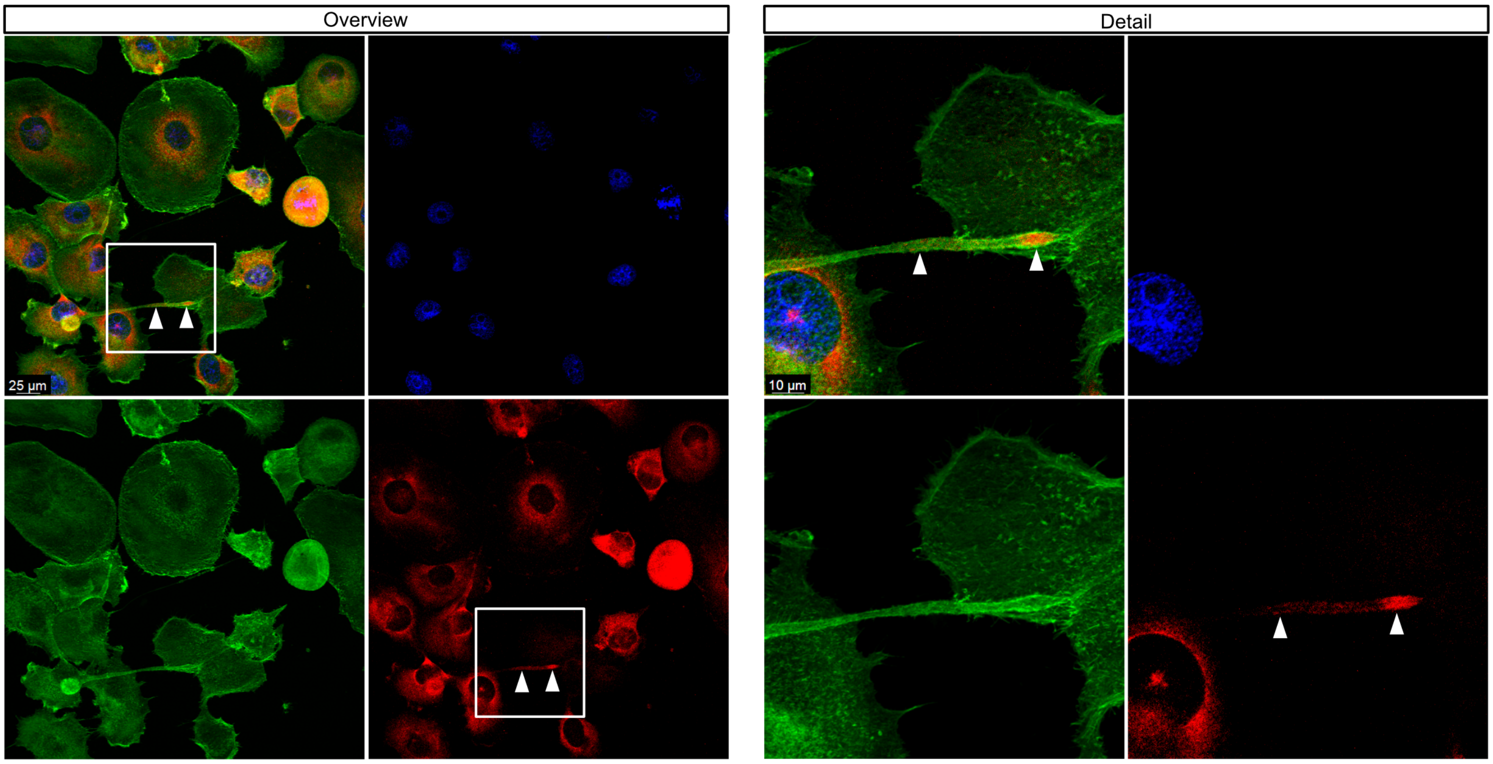


Overlay and channels (DNA – blue, actin – green, RPS6 antibody – red)

**b.** PANC-1 RPS6 #

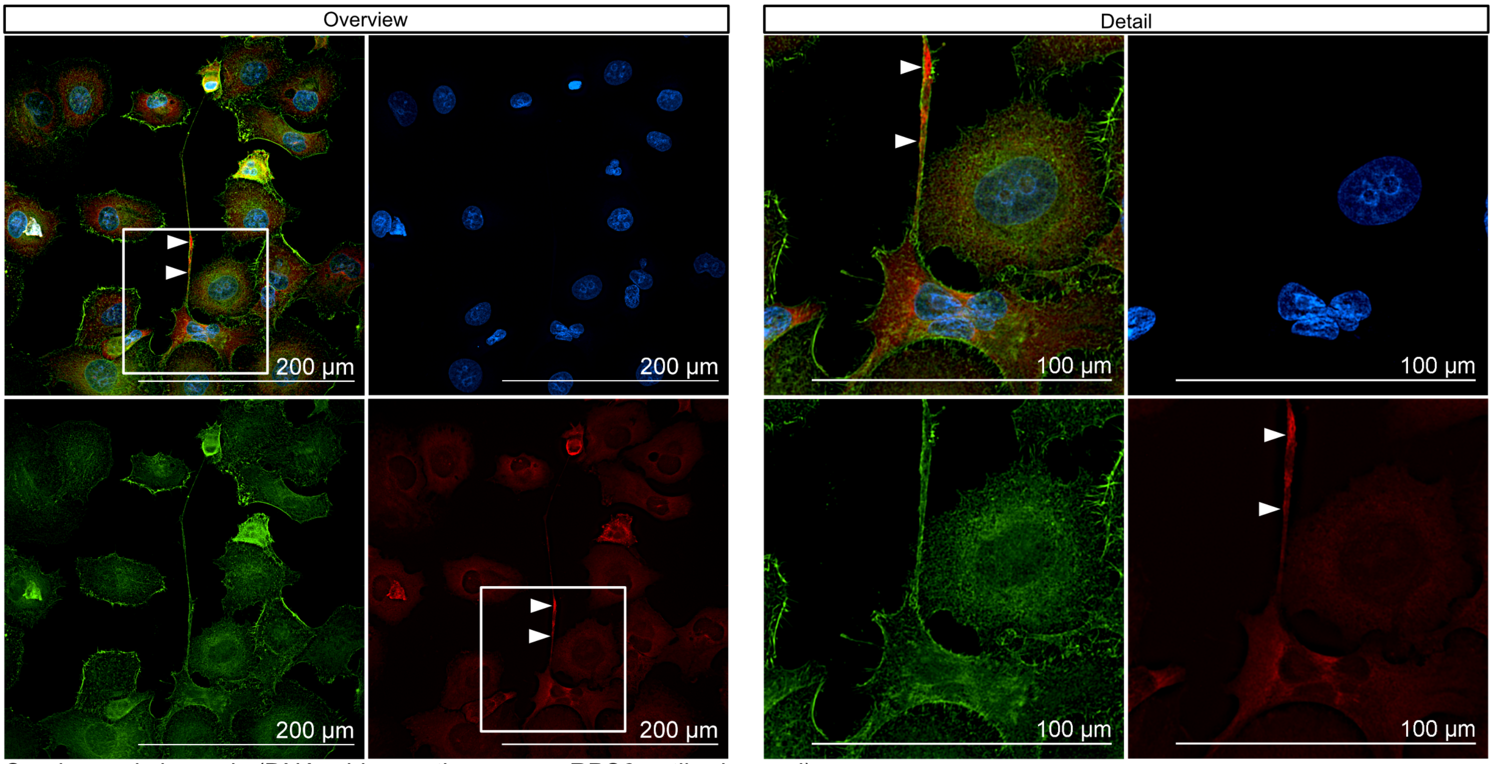


Overlay and channels (DNA – blue, actin – green, RPS6 antibody – red)

### FIGURE S2B cont.

**Figure S2B.** Ribosomal protein S6 detected in tunnelling nanotubes (a, b, c).

**c.** PANC-1 RPS6 #

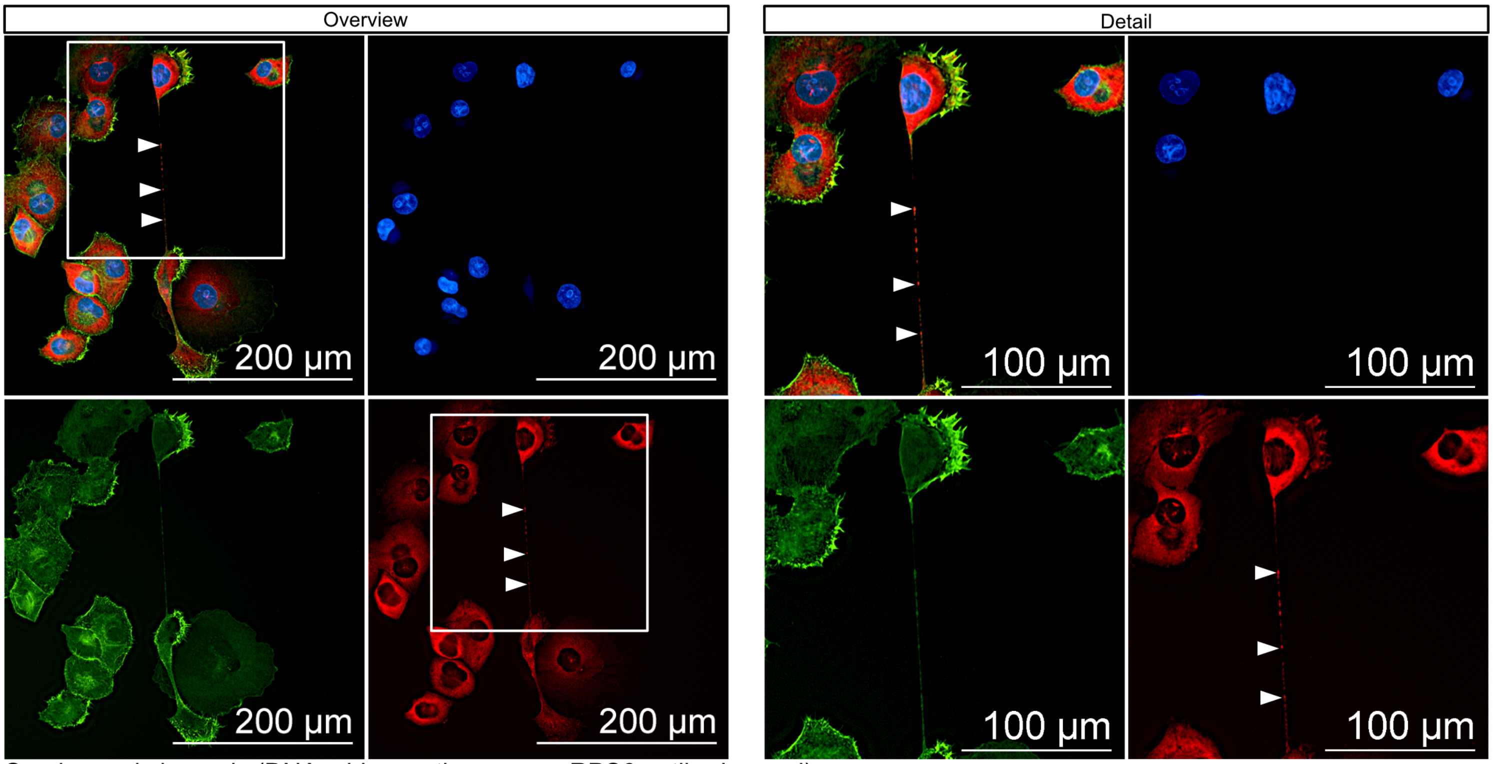


Overlay and channels (DNA – blue, actin – green, RPS6 antibody – red)

### FIGURE S2C

**Figure S2C.** Ribosomal protein L24 detected in tunnelling nanotubes (a, b, c).

**a.** PANC-1 RPL24 *

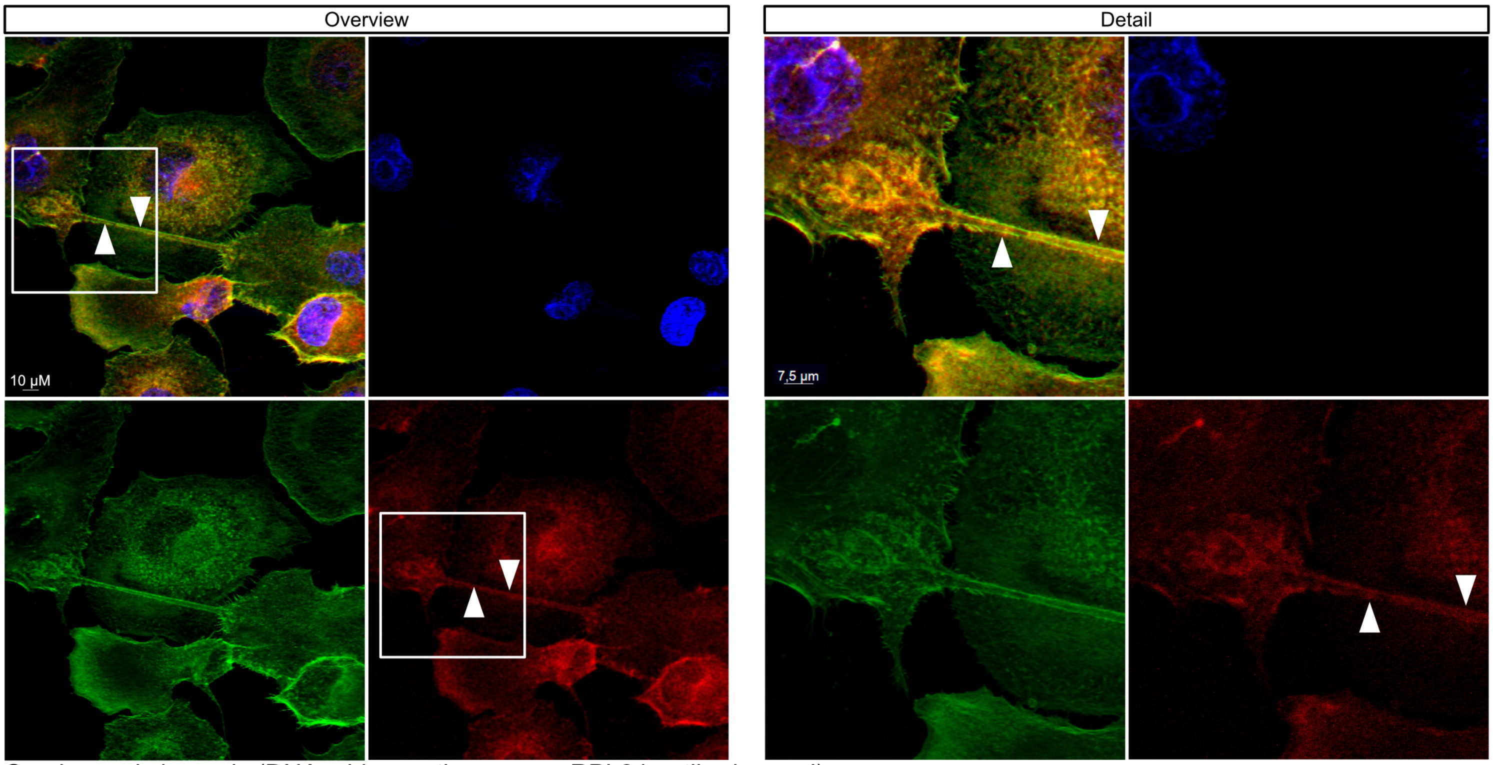


Overlay and channels (DNA – blue, actin – green, RPL24 antibody – red)

**b.** PANC-1 RPL24 *

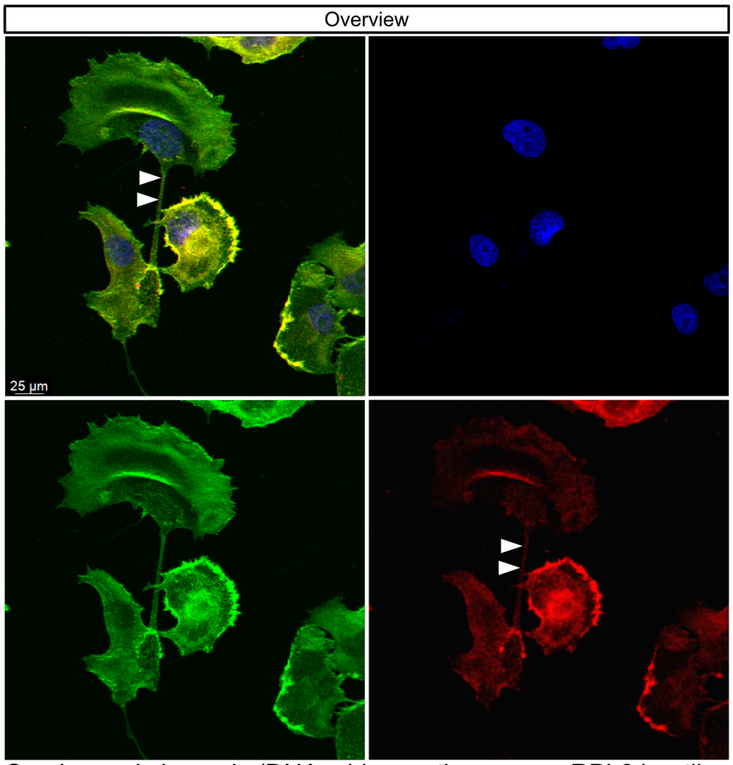


**c.** PANC-1 RPL24 *

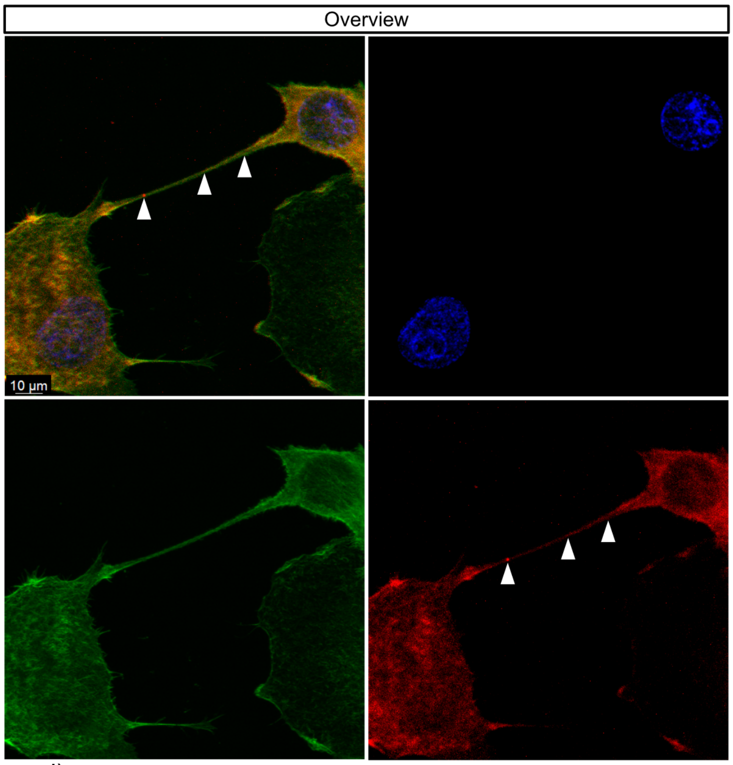


Overlay and channels (DNA – blue, actin – green, RPL24 antibody – red)

### FIGURE S2D

**Figure S2D.** Poly-A mRNA detected in tunnelling nanotubes (a, b, c).

**a.** PANC-1 mRNA *

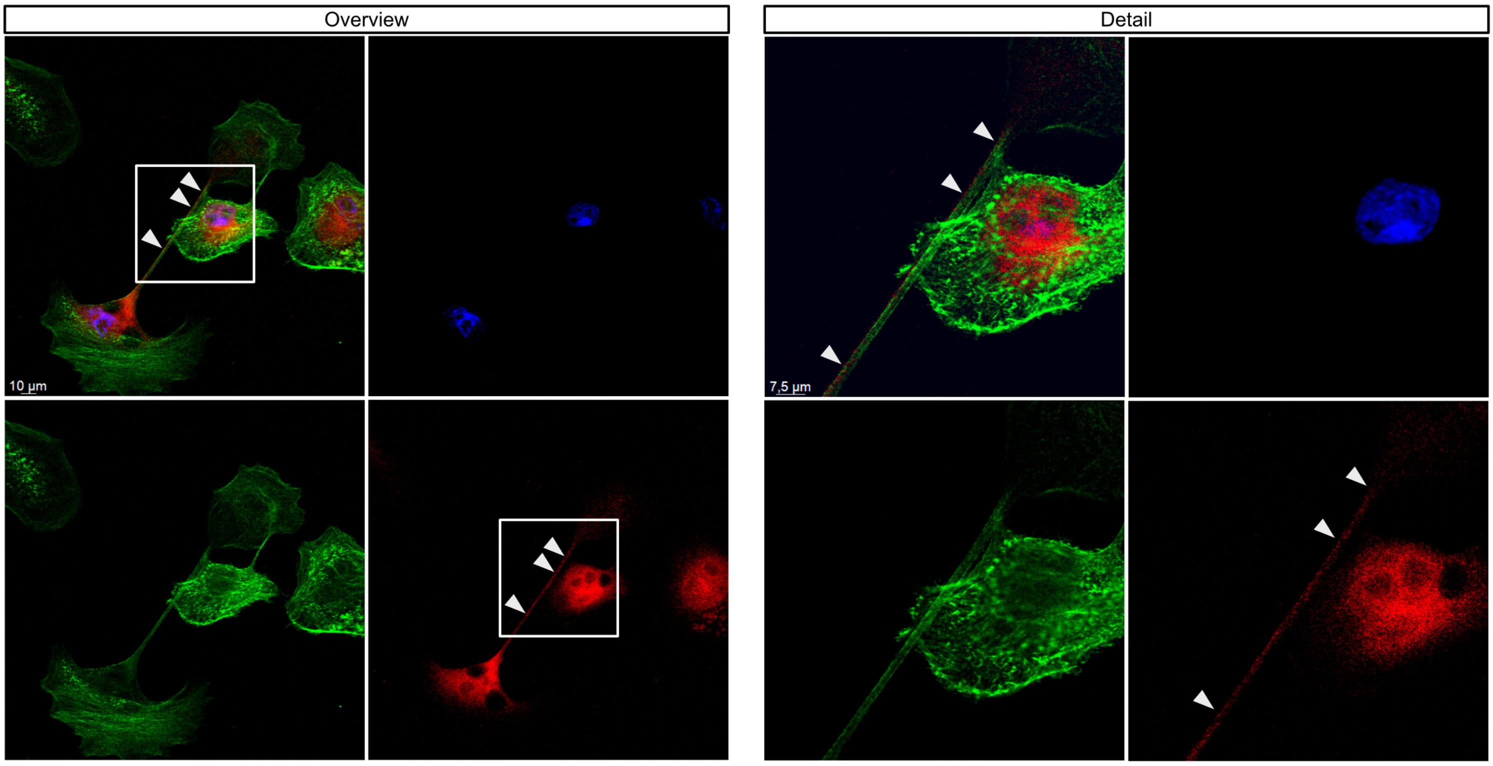


Overlay and channels (DNA – blue, actin – green, polyA probe – red)

**b.** PANC-1 mRNA *

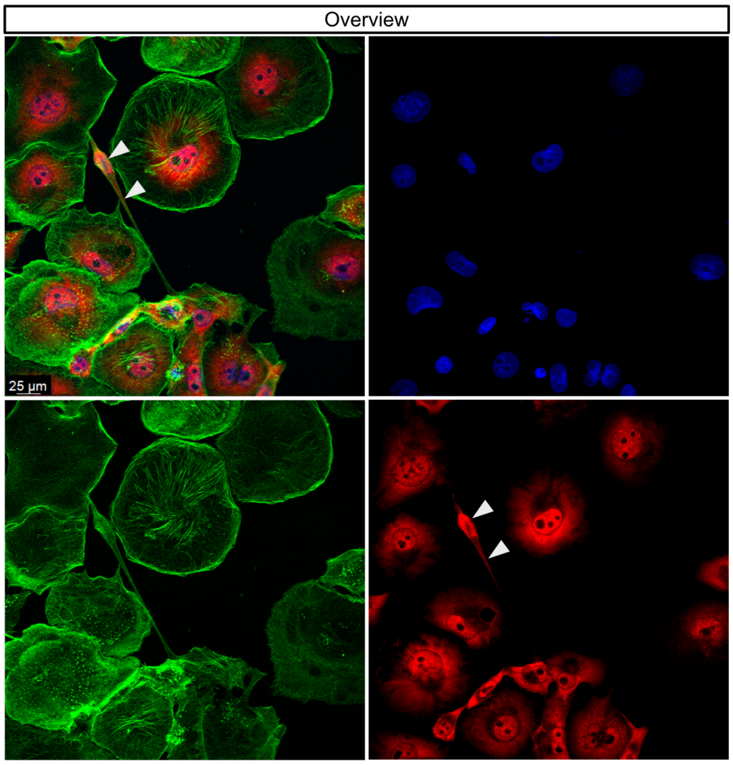


**c.** PANC-1 mRNA *

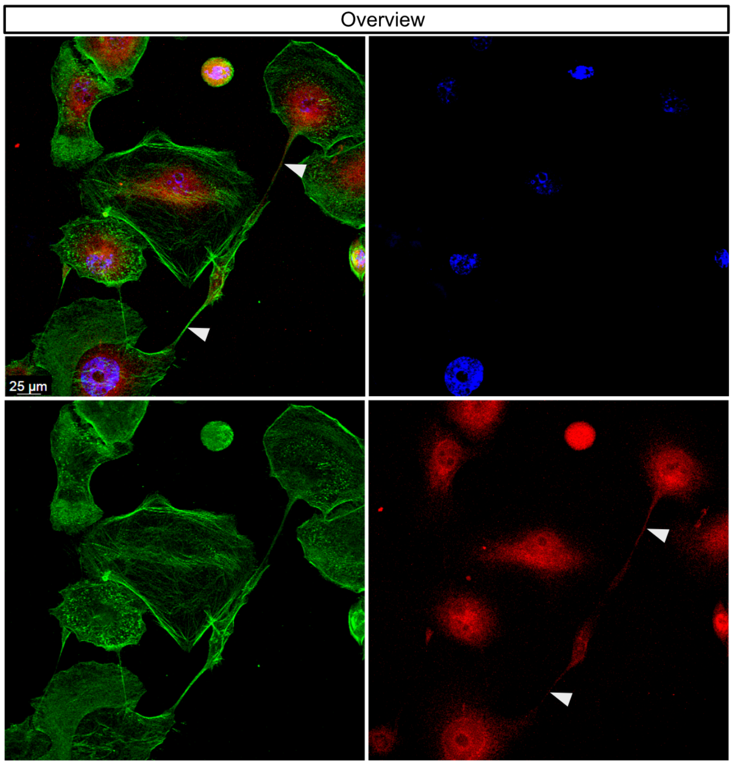


Overlay and channels (DNA – blue, actin – green, polyA probe – red)

### FIGURE S2E

**Figure S2E.** Complete ribosomes detected in tunnelling nanotubes (a, b, c).

**a.** PANC-1 Complete ribosomes (proximity ligation) *

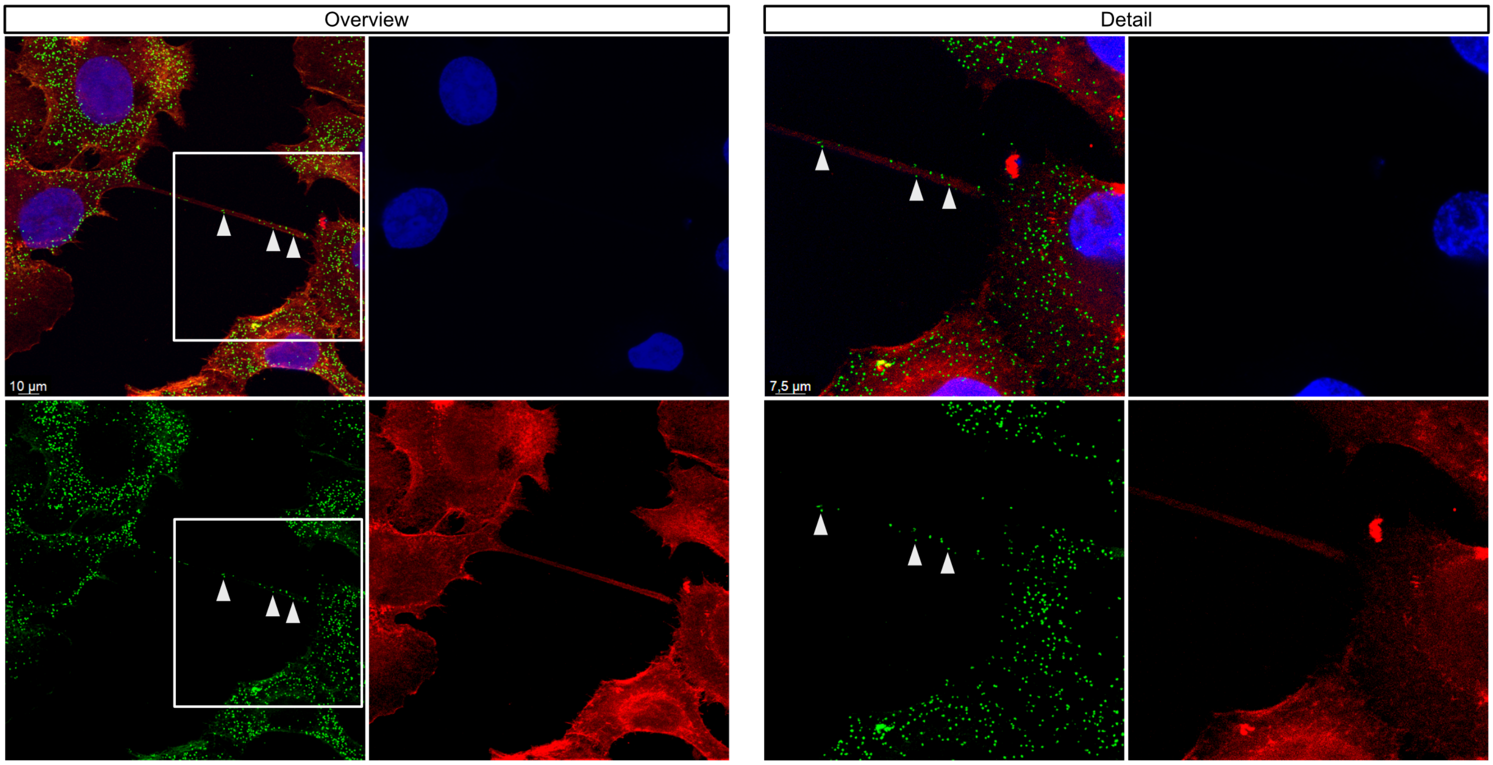


Overlay and channels (DNA – blue, RPS6-RPL24 proximity ligation– green, actin – red)

**b.** PANC-1 Complete ribosomes (proximity ligation) *

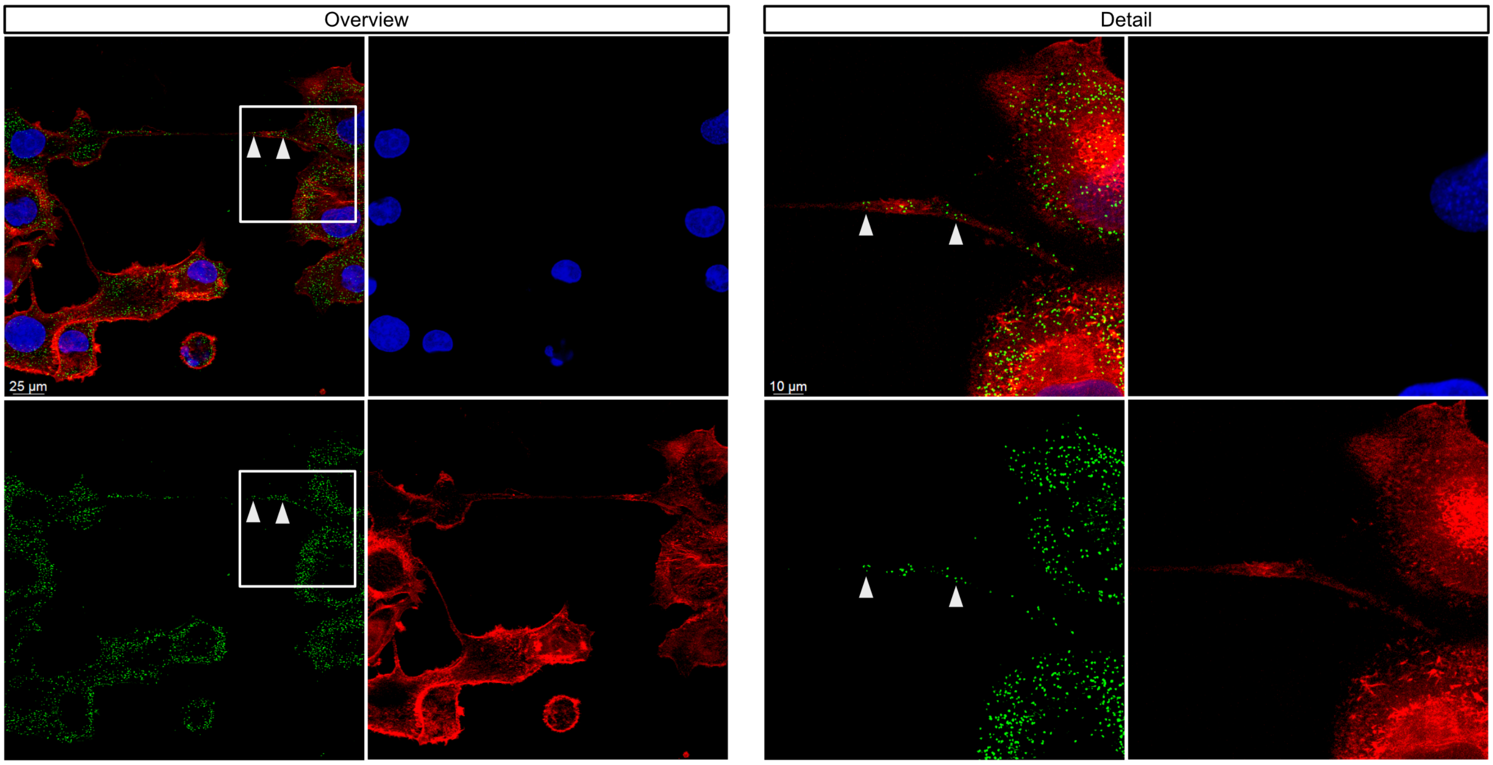


Overlay and channels (DNA – blue, RPS6-RPL24 proximity ligation– green, actin – red)

### FIGURE S2E cont.

**Figure S2E.** Complete ribosomes detected in tunnelling nanotubes (a, b, c).

**c.** PANC-1 Complete ribosomes (proximity ligation) *

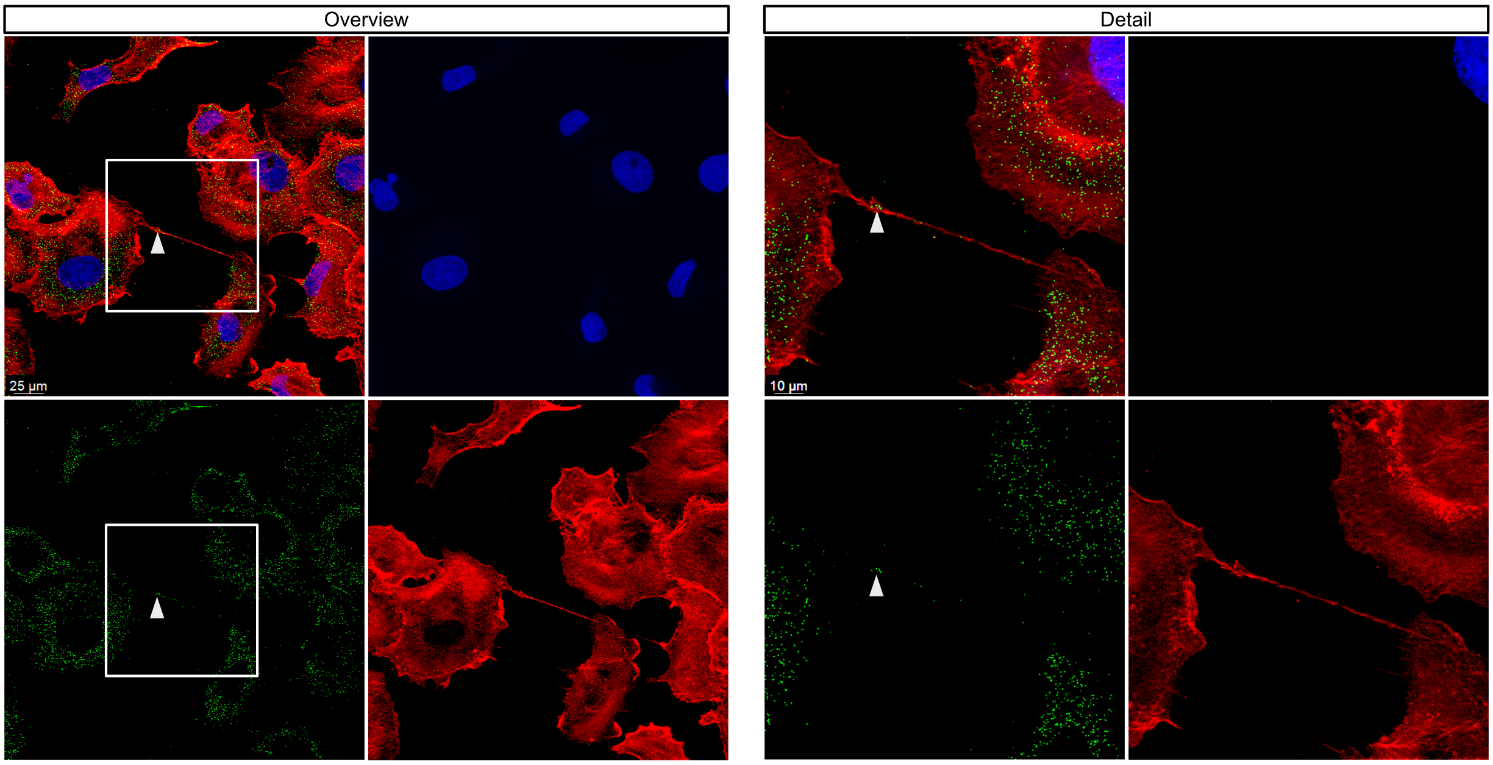


Overlay and channels (DNA – blue, RPS6-RPL24 proximity ligation– green, actin – red)

### FIGURE S2F

**Figure S2F.** Ribosomal protein S9 (a).

**a.** PANC-1 RPS9 *

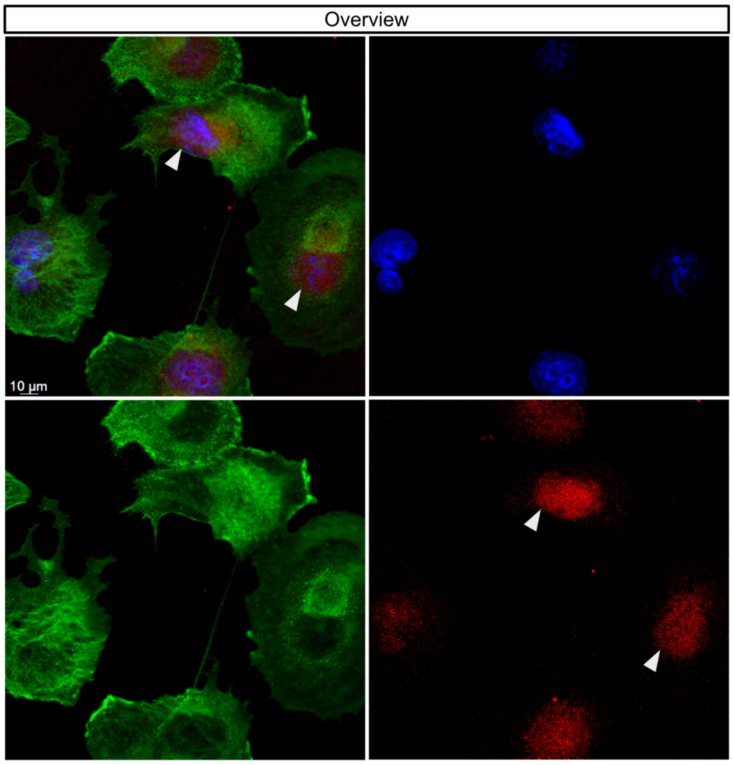


Overlay and channels (DNA – blue, actin – green, RPS9 antibody – red)

### FIGURE S3A

**Figure S3A.** Ribosomal 5.8S rRNA detected in patient BJPN26 tunnelling nanotubes (a, b).

**a.** Patient sample BJPN26 5.8S rRNA #

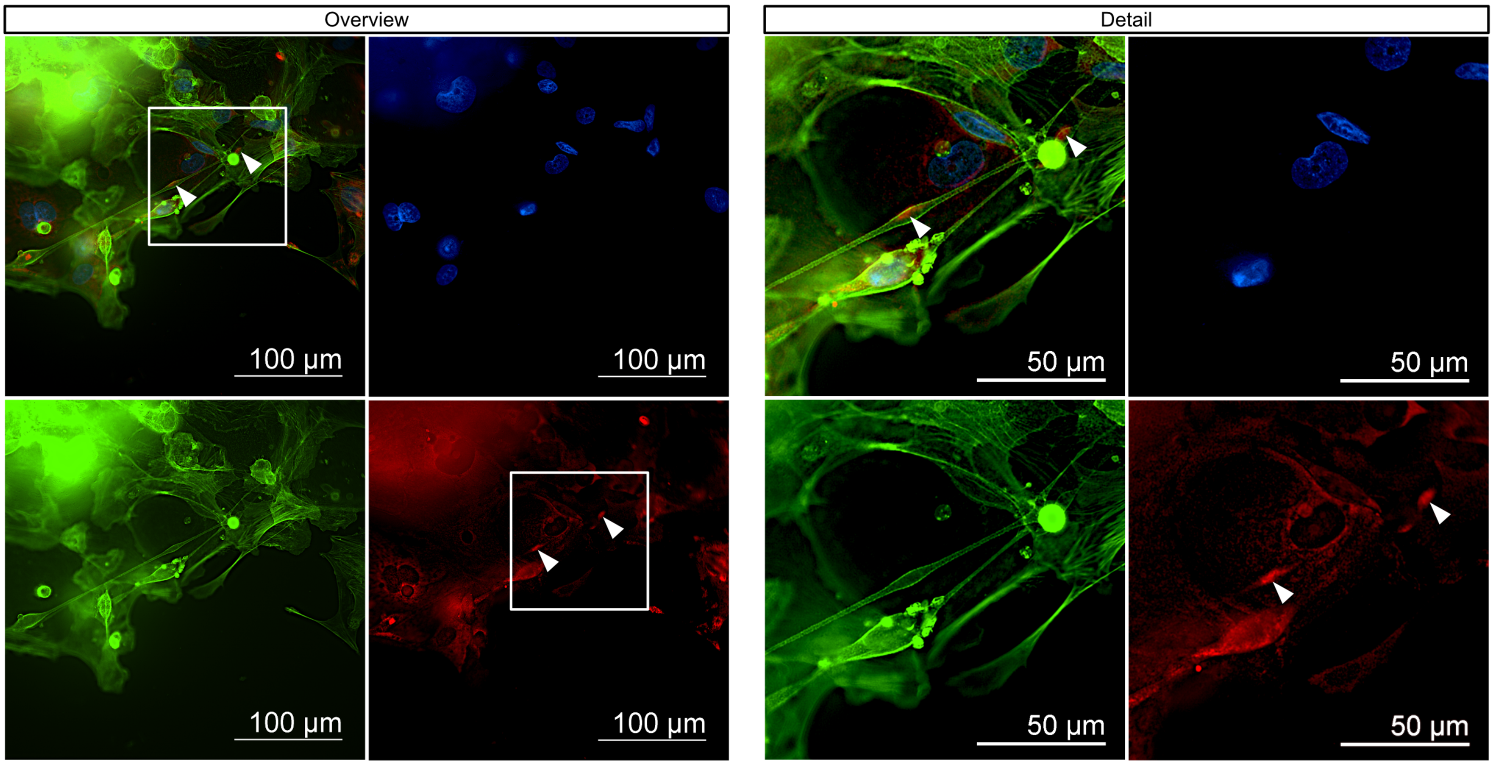


Overlay and channels (DNA – blue, actin – green, 5.8S rRNA antibody – red)

**b.** Patient sample BJPN26 5.8S rRNA #

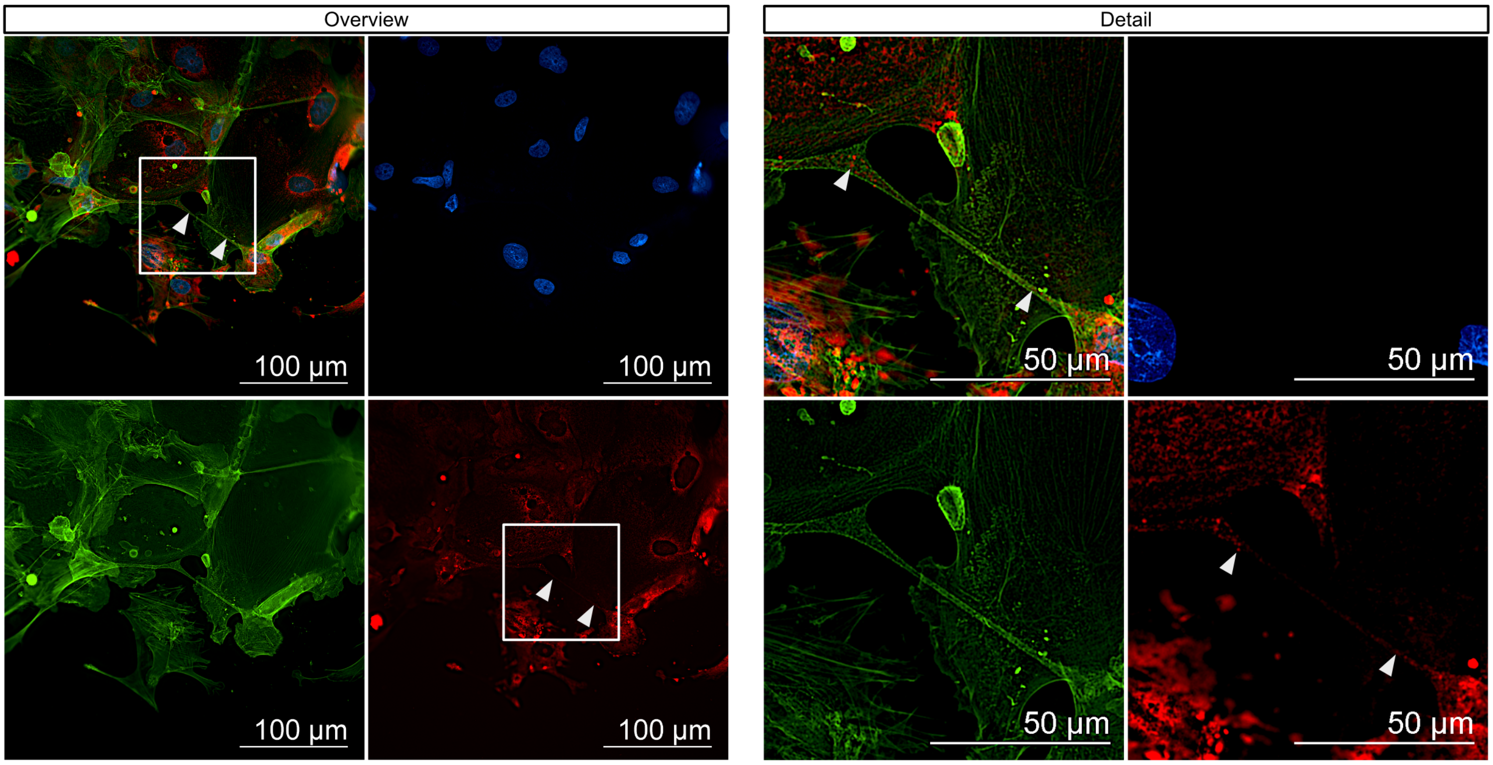


Overlay and channels (DNA – blue, actin – green, 5.8S rRNA antibody – red)

### FIGURE S3B

**Figure S3B.** Ribosomal 5.8S rRNA detected in patient BJPN24 tunnelling nanotubes (a, b).

**a.** Patient sample BJPN24 5.8S rRNA #

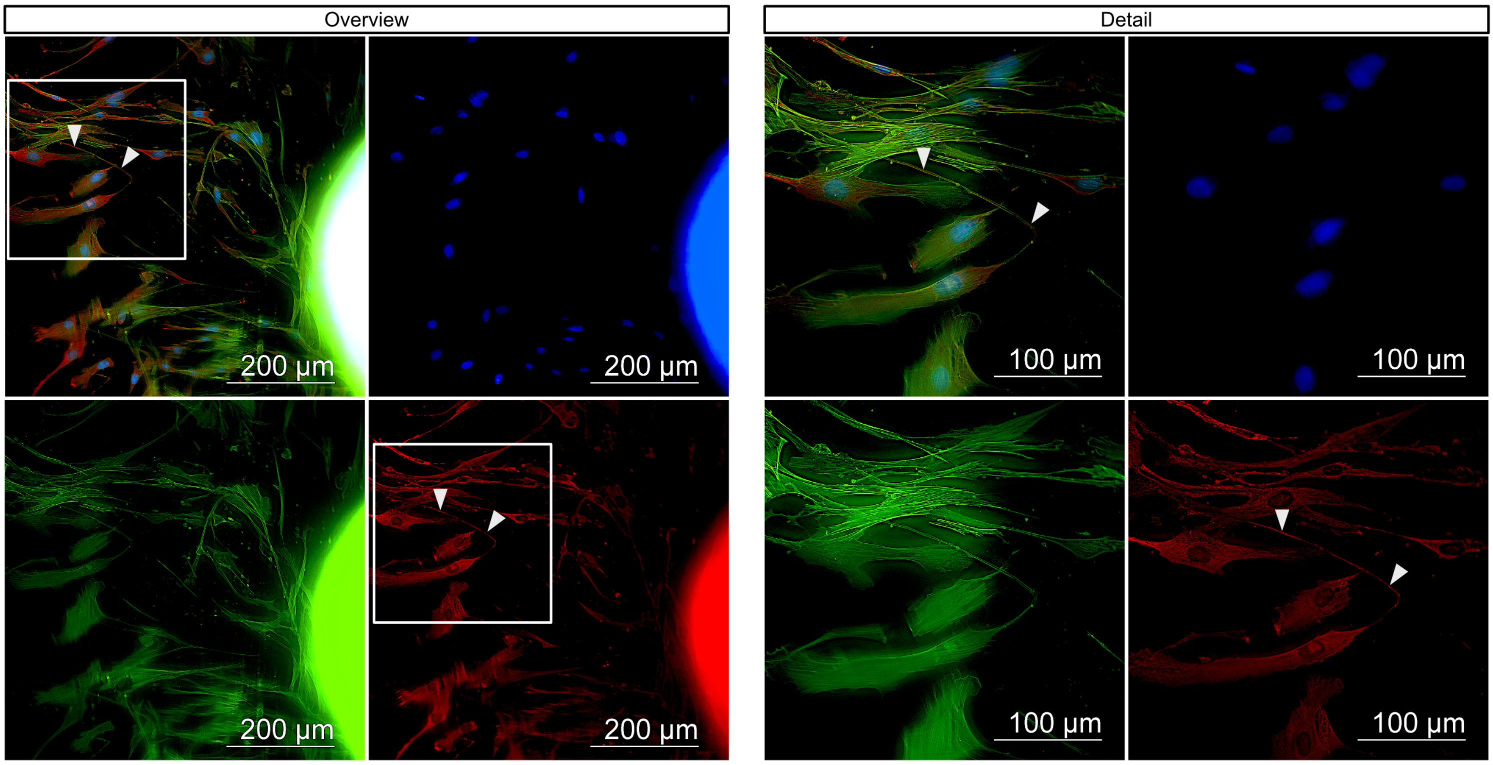


Overlay and channels (DNA – blue, actin – green, 5.8S rRNA antibody – red)

**b.** Patient sample BJPN24 5.8S rRNA #

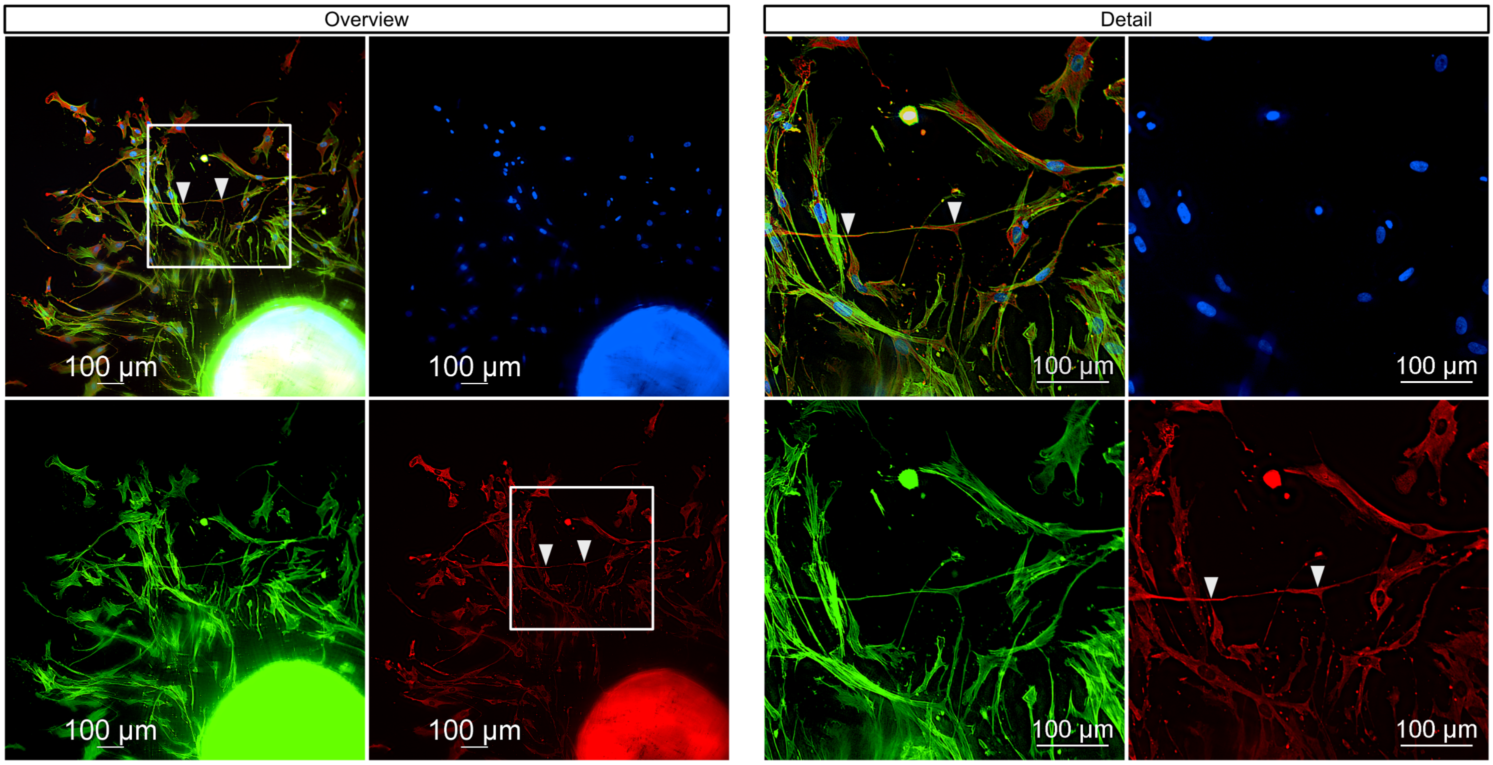


Overlay and channels (DNA – blue, actin – green, 5.8S rRNA antibody – red)

### FIGURE S3C

**Figure S3C.** Presence of cytokeratin-19 (CK-19) in tunnelling nanotubes of patient BJPN26 (a, b, c).

**a.** Patient sample BJPN26 cytokeratin-19 in TNTs #

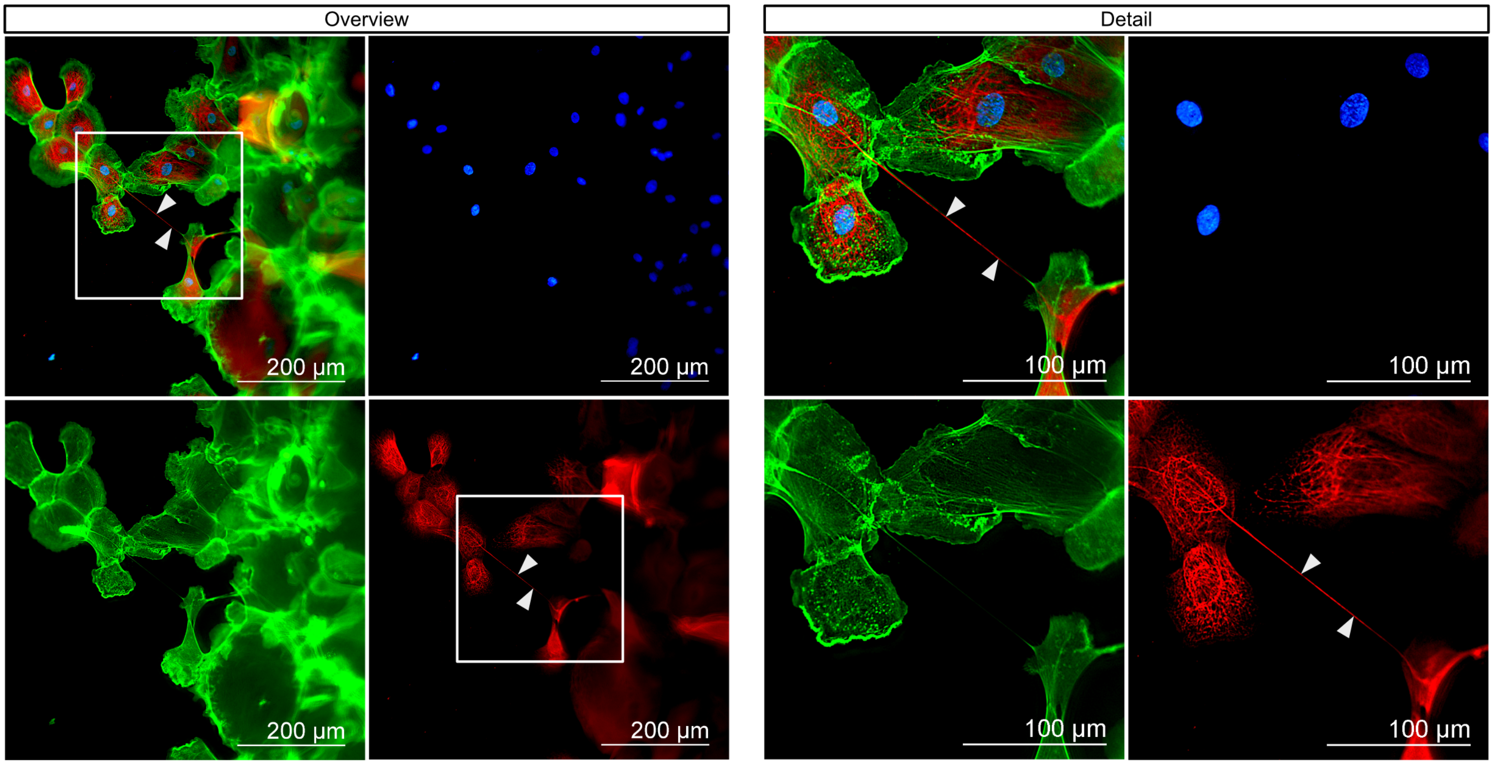


Overlay and channels (DNA – blue, actin– green, cytokeratin-19 (CK-19) antibody – red)

**b.** Patient sample BJPN26 cytokeratin-19 in TNTs #

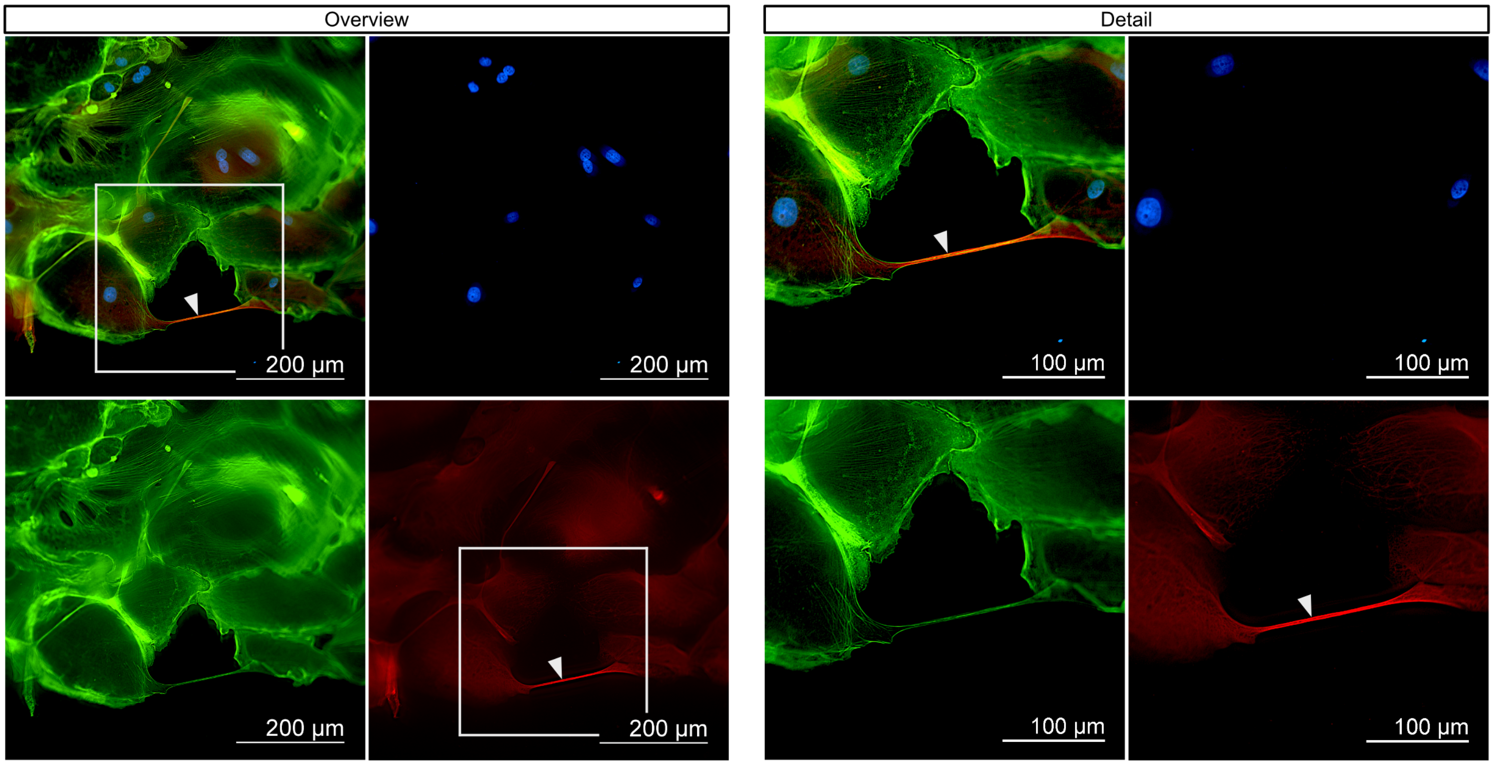


Overlay and channels (DNA – blue, actin– green, cytokeratin-19 (CK-19) antibody – red)

### FIGURE S3C cont.

**Figure S3C.** Presence of cytokeratin-19 (CK-19) in tunnelling nanotubes of patient BJPN26 (a, b, c).

**c.** Patient sample BJPN26 cytokeratin-19 in TNTs #

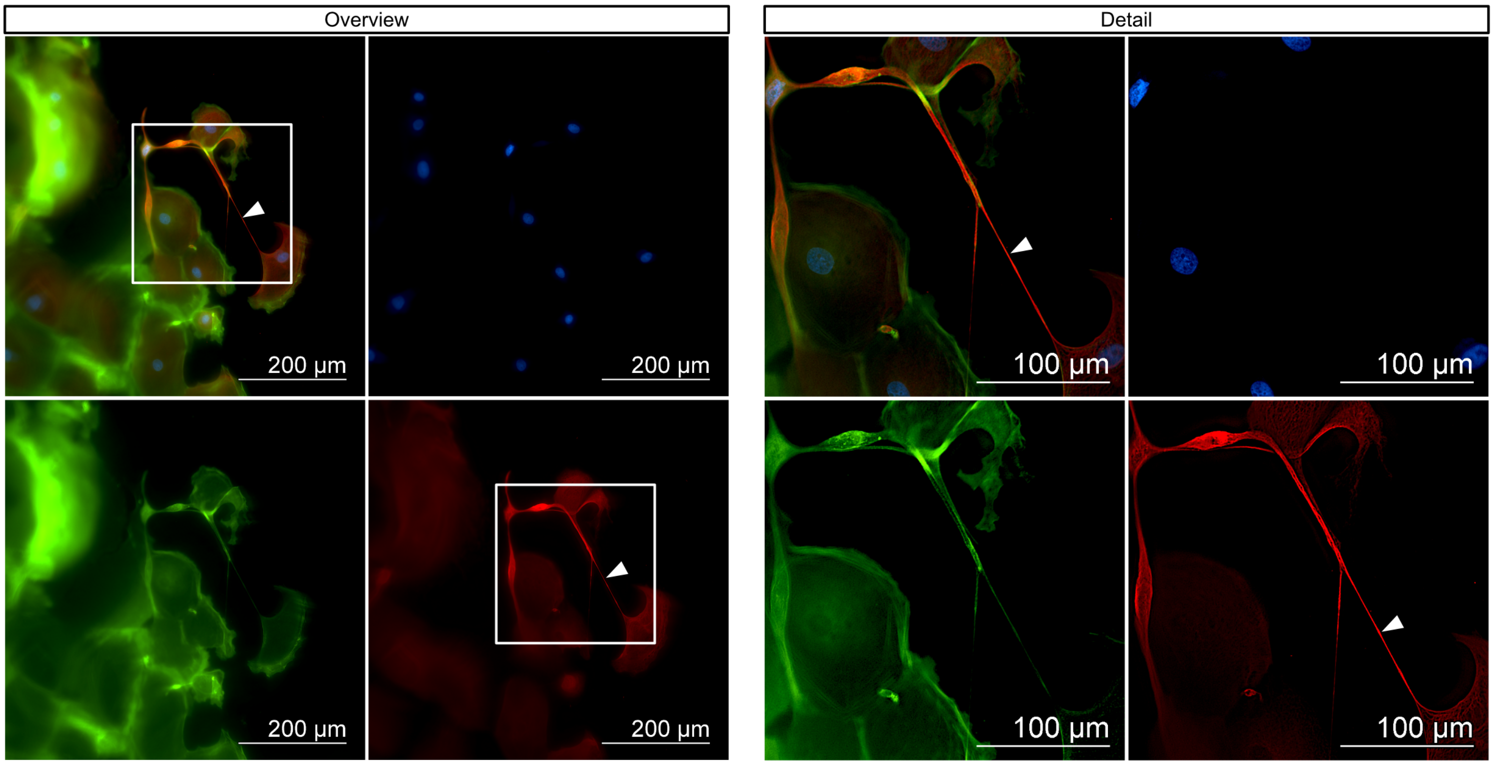


Overlay and channels (DNA – blue, actin– green, cytokeratin-19 (CK-19) antibody – red)

### FIGURE S3D

**Figure S3D.** Presence of vimentin (a, b, c) in tunnelling nanotubes of patient BJPN24, whereas cytokeratin-19 (CK-19) is absent (d).

**a.** Patient sample BJPN24 vimentin in TNTs #

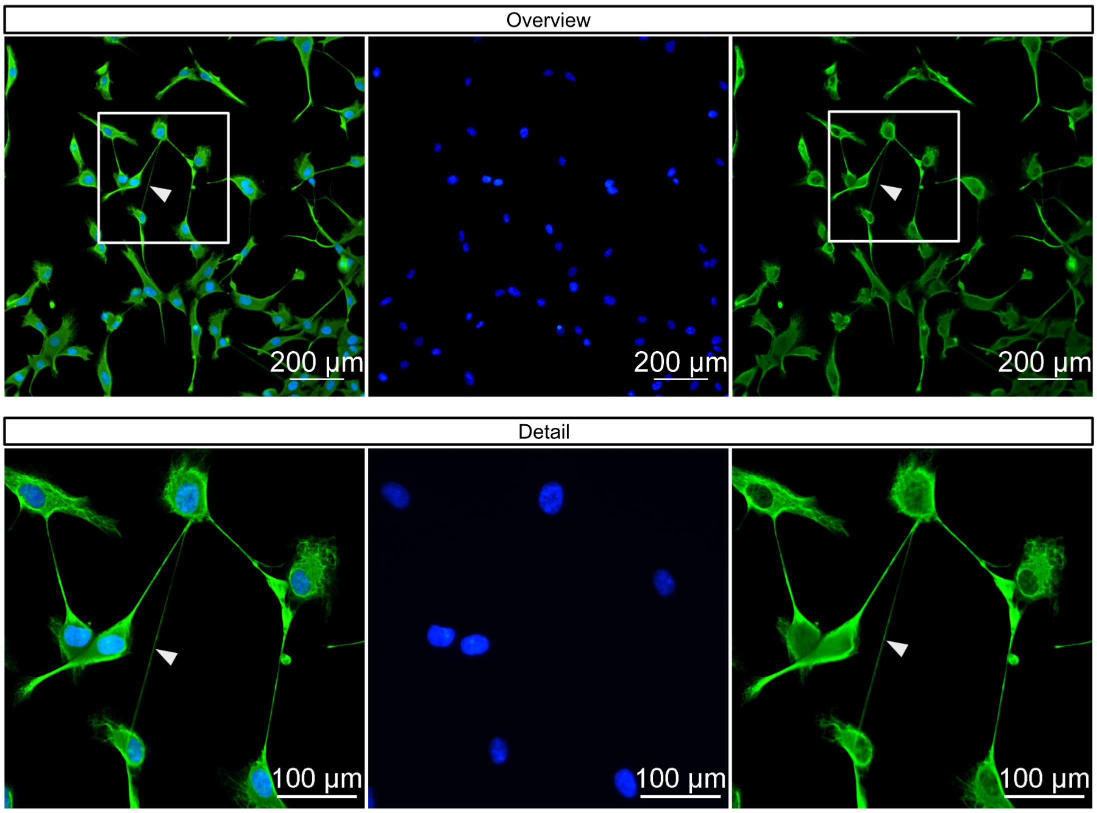


Overlay and channels (DNA – blue, vimentin antibody – green)

**b.** Patient sample BJPN24 vimentin in TNTs #

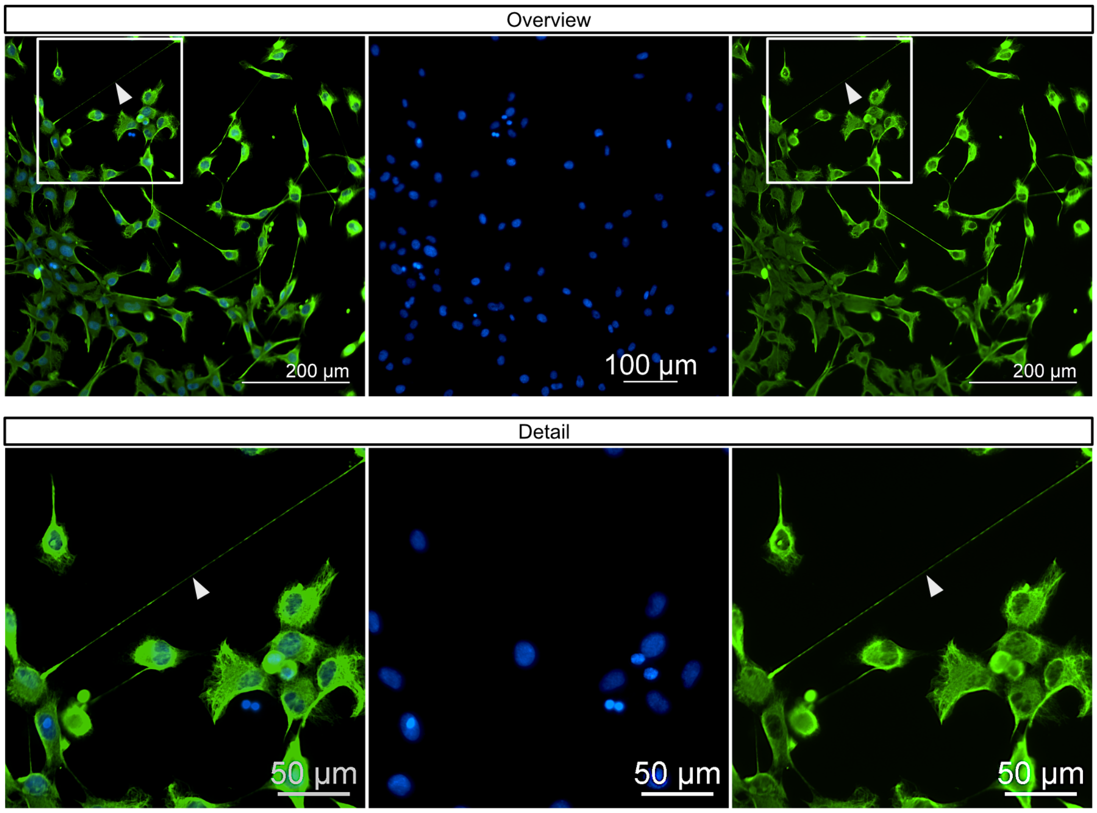


Overlay and channels (DNA – blue, vimentin antibody – green)

### FIGURE S3D cont.

**Figure S3D.** Presence of vimentin (a, b, c) in tunnelling nanotubes of patient BJPN24, whereas cytokeratin-19 (CK-19) is absent (d).

**c.** Patient sample BJPN24 vimentin in TNTs #

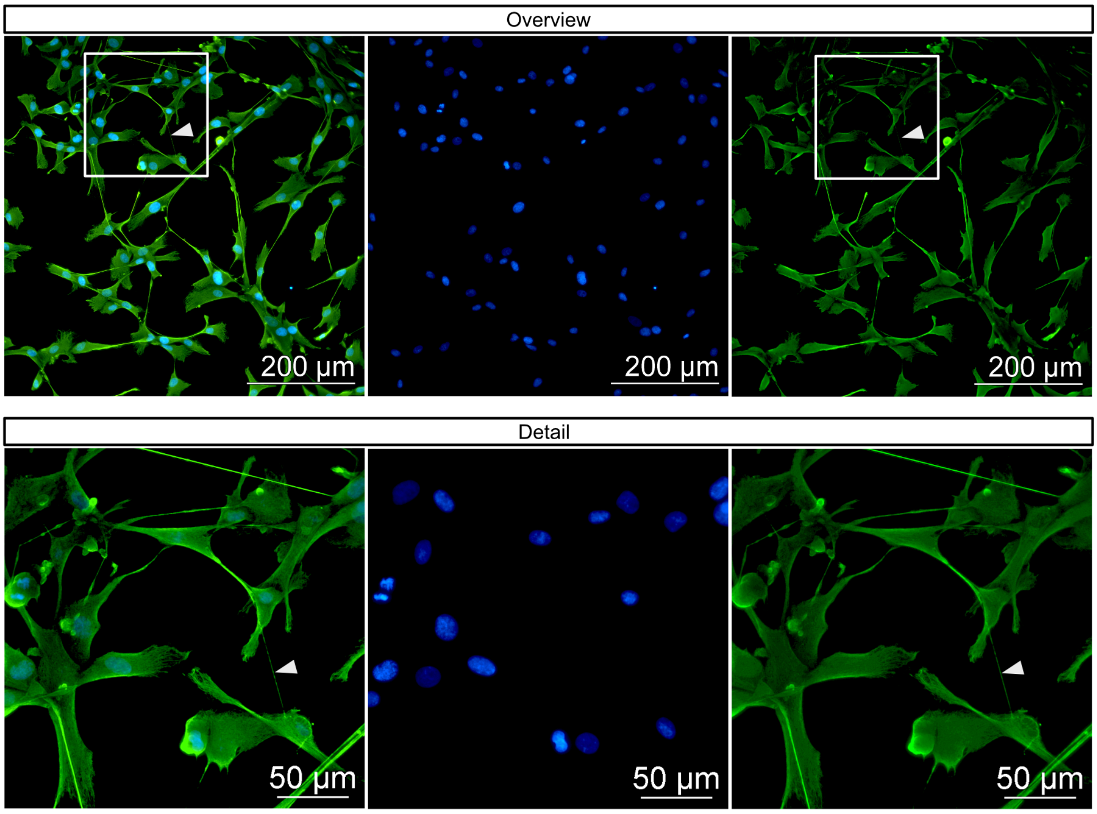


Overlay and channels (DNA – blue, vimentin antibody – green)

**d.** Patient sample BJPN24 vimentin in TNTs #

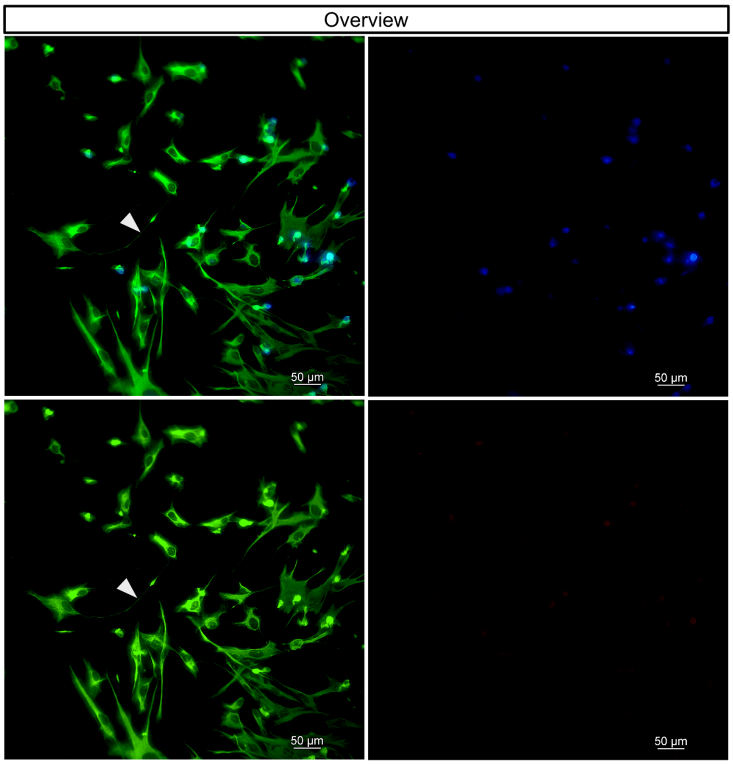


Overlay and channels (DNA – blue, vimentin antibody – green, cytokeratin-19 (CK-19) antibody – red)

### FIGURE S3E

**Figure S3E.** Complete ribosomes detected in patient BJPN26 tunnelling nanotubes (a, b, c).

**a.** Patient BJPN26 sample complete ribosomes *

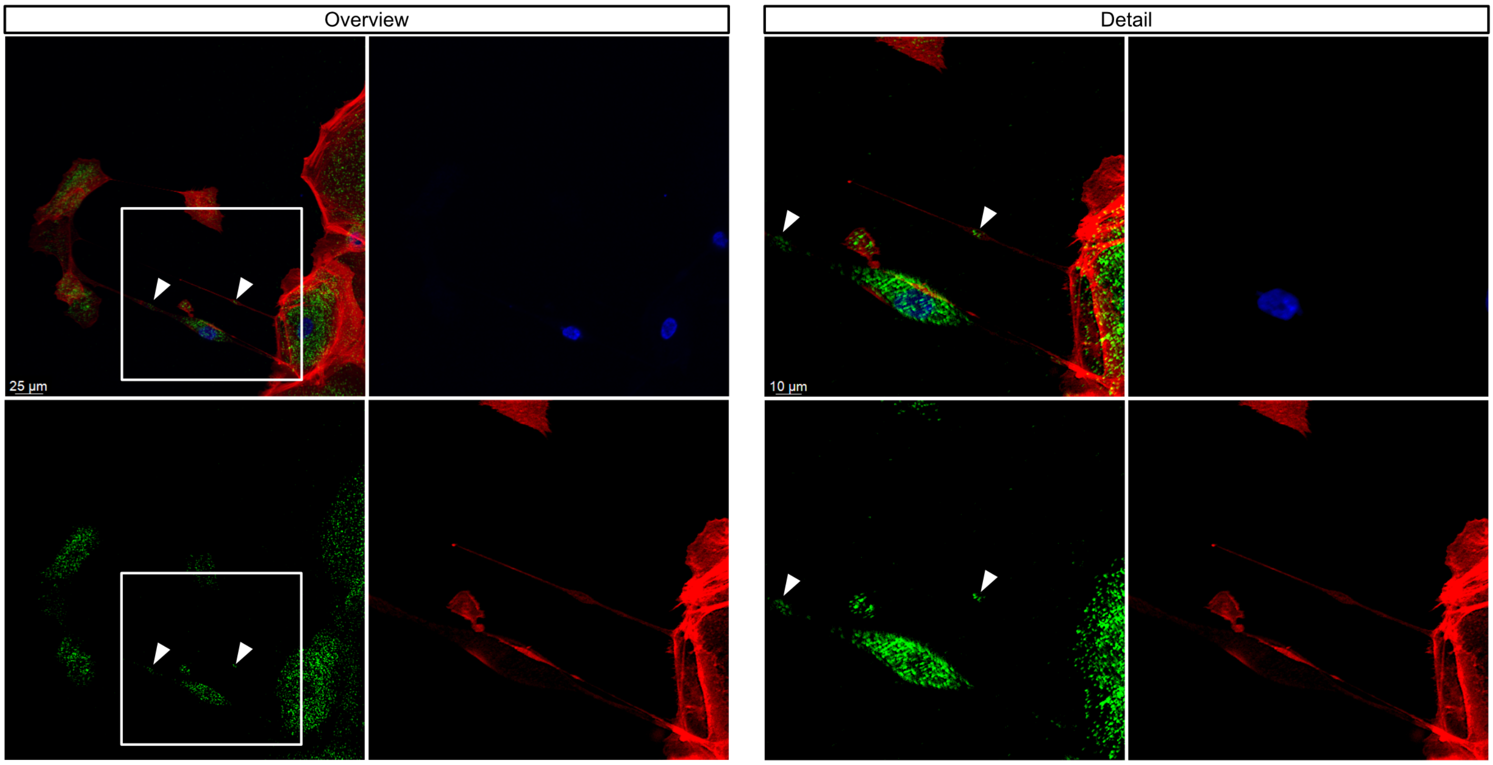


Overlay and channels (DNA – blue, RPS6+RPL24 proximity ligation– green, actin – red)

**b.** Patient BJPN26 sample complete ribosomes *

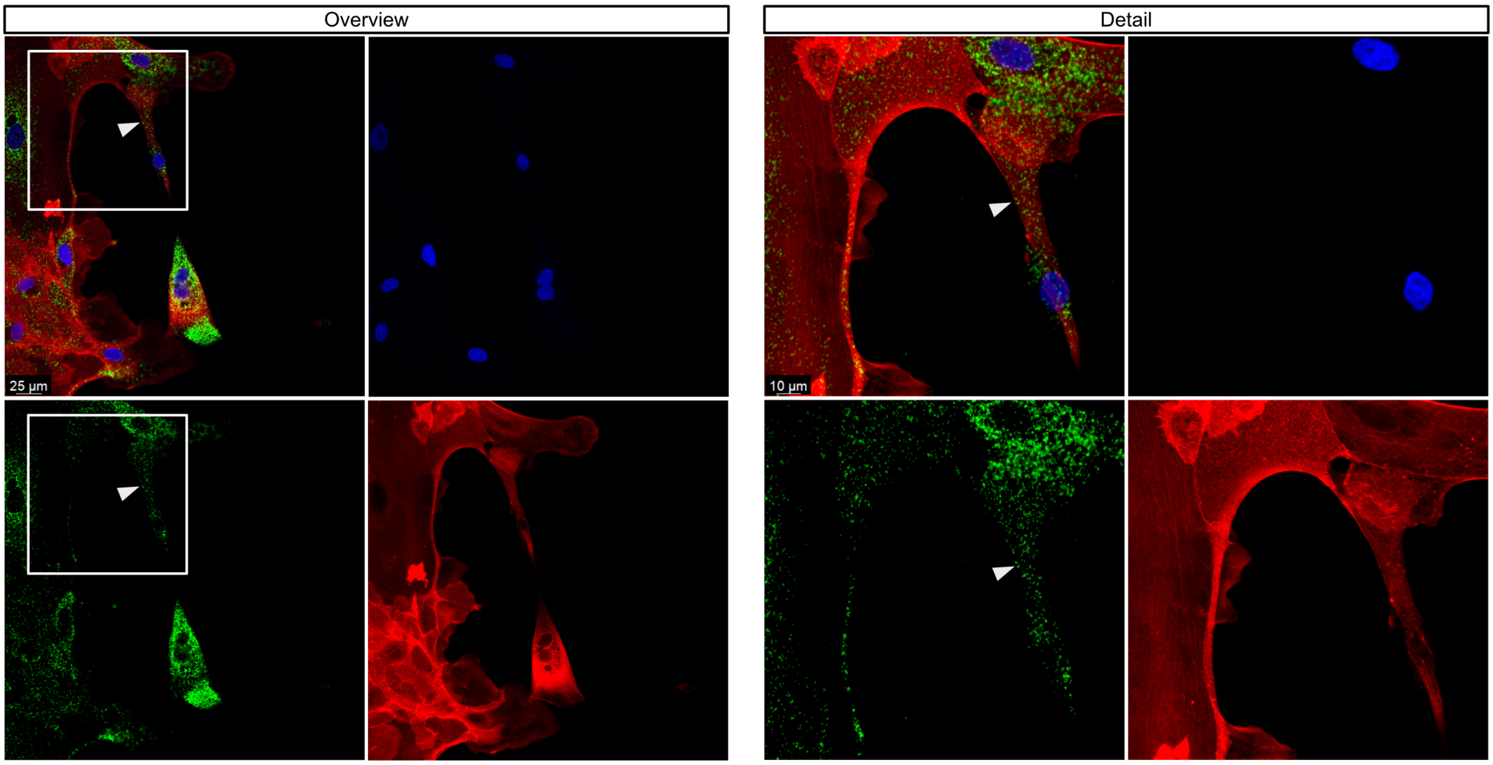


Overlay and channels (DNA – blue, RPS6+RPL24 proximity ligation– green, actin – red)

### FIGURE S3E cont.

**Figure S3E.** Complete ribosomes detected in patient BJPN26 tunnelling nanotubes (a, b, c).

**c.** Patient BJPN26 sample complete ribosomes *

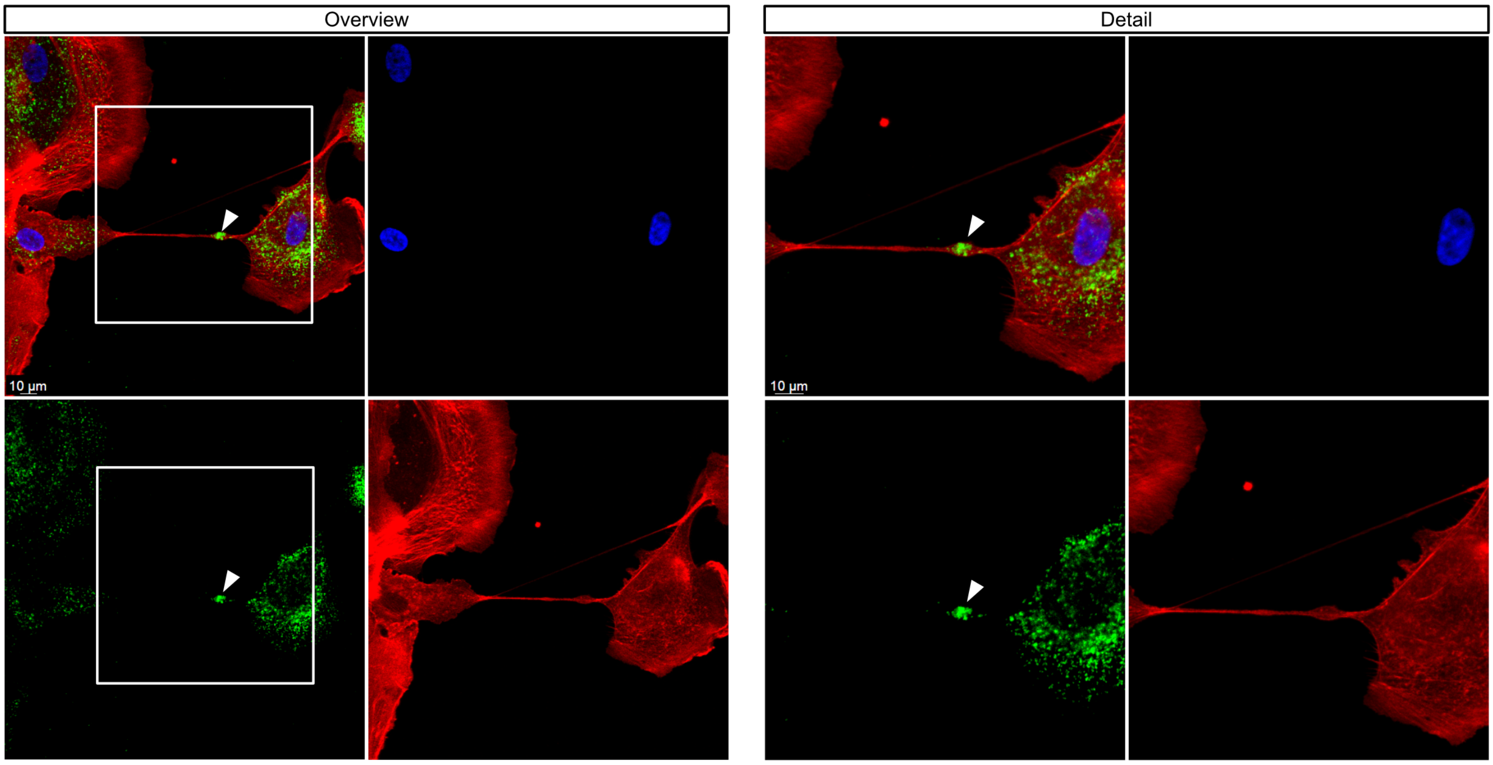


Overlay and channels (DNA – blue, RPS6+RPL24 proximity ligation– green, actin – red)

### FIGURE S3F

**Figure S3F.** Complete ribosomes detected in patient BJPN24 tunnelling nanotubes (a, b).

**a.** Patient BJPN24 sample complete ribosomes *

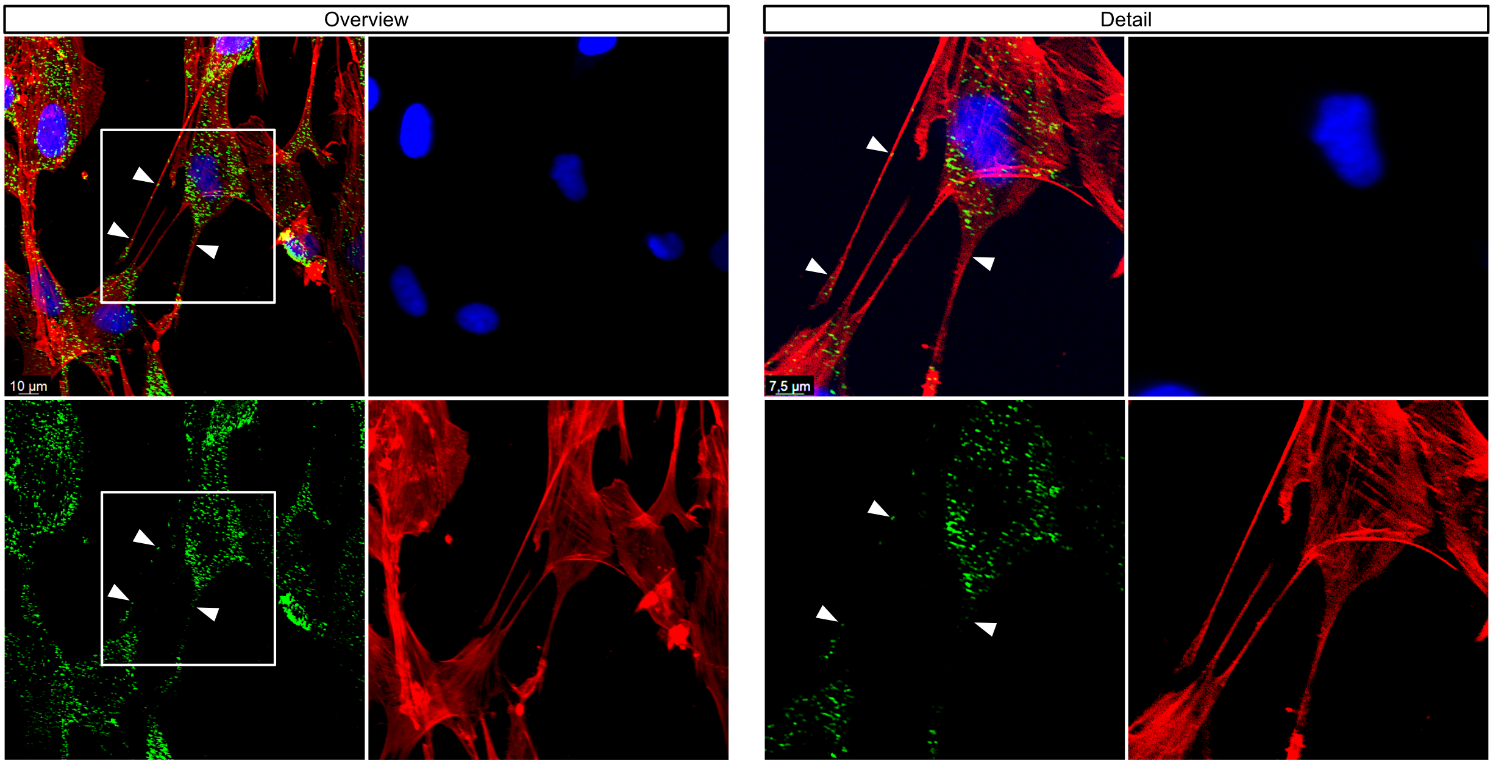


Overlay and channels (DNA – blue, RPS6-RPL24 proximity ligation– green, actin – red)

**b.** Patient BJPN24 sample complete ribosomes *

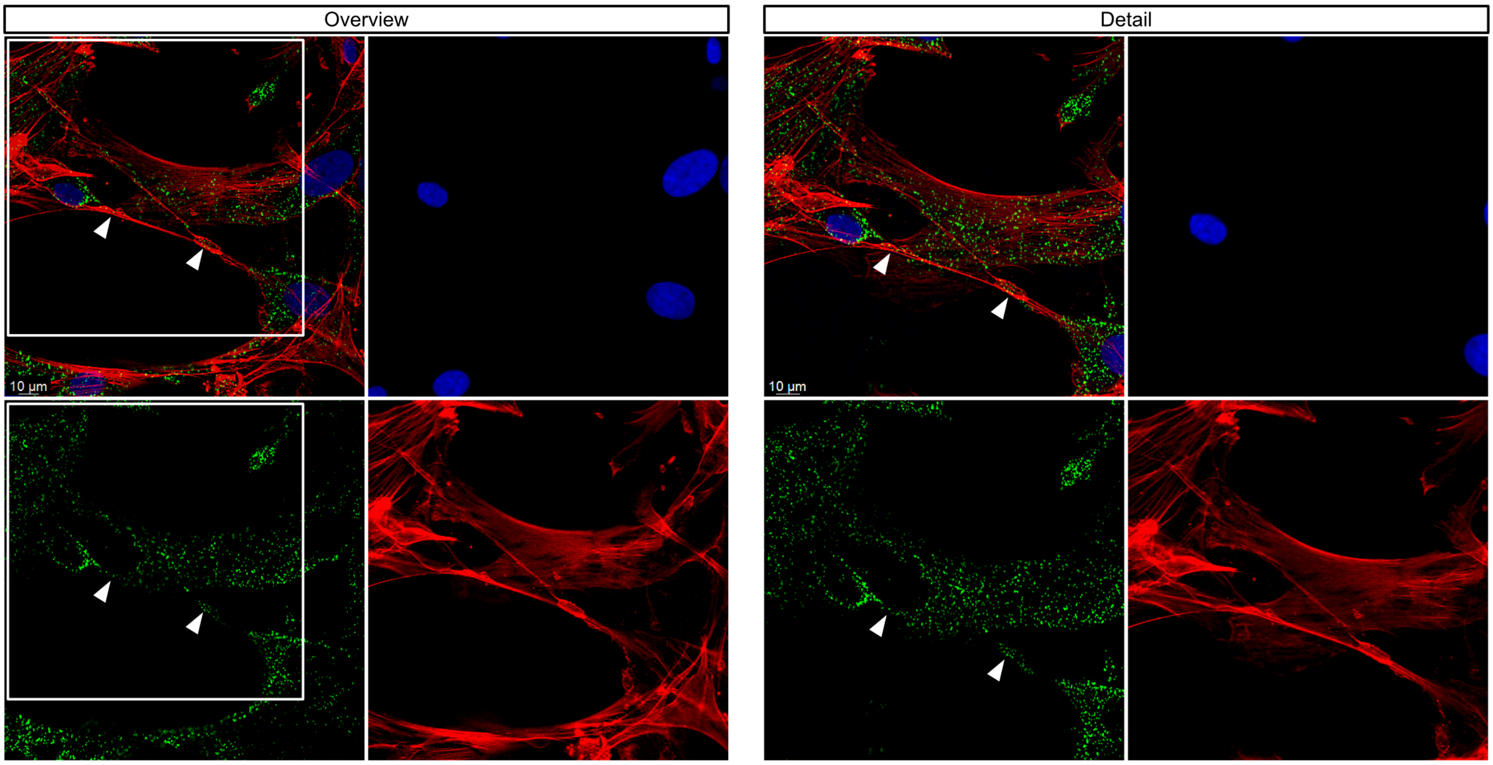


Overlay and channels (DNA – blue, RPS6-RPL24 proximity ligation– green, actin – red)

### FIGURE S4A

**Figure S4A.** HaloTag-RPS9 transfected PANC-1 cells (a) # and transport of HaloTag-RPS9 (red) to non-transfected (blue) PANC-1 cells (b) #.

**a.** HaloTag-RPS9 transfected PANC-1 cells #

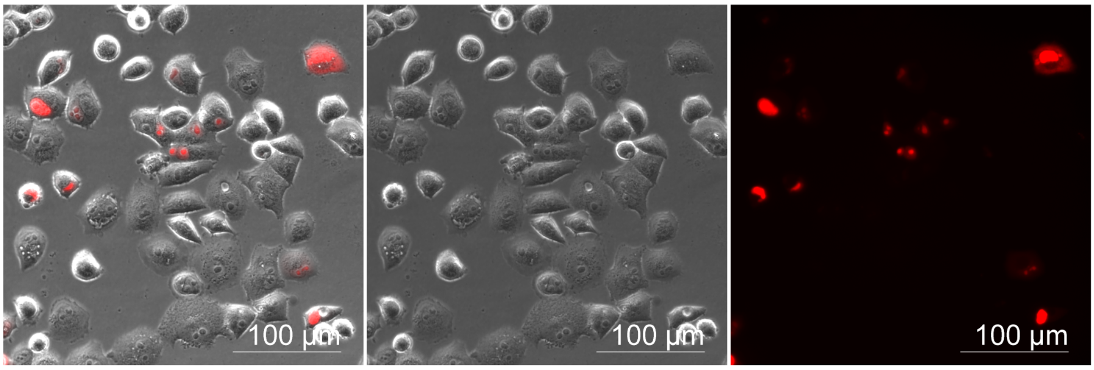


Overlay and channels (Bright-field, HaloTag-TMR ligand – red)

**b.** Transport of HaloTag-RPS9 (red) to non-transfected (blue) PANC-1 cells #

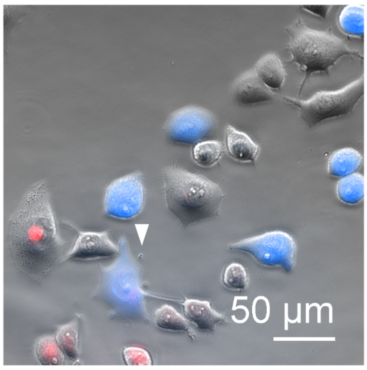


Overlay (CellTrace Violet stain – blue, HaloTag-TMR ligand – red)

### FIGURE S4B

**Figure S4B.** Transport of HaloTag-RPS9 (red) to nuclear GFP transfected (green) PANC-1 cells (a) #.

**a.** HaloTag-RPS9 transport to nuclear GFP transfected PANC-1 cells #

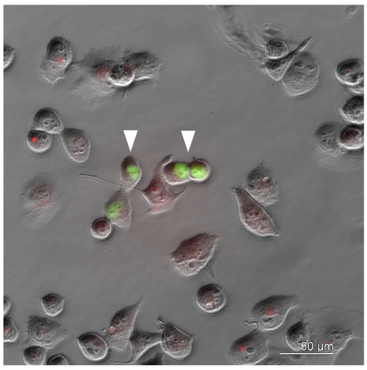


Overlay (GFP expressed in the nucleus - green, HaloTag-TMR ligand – red)

### FIGURE S4C

**Figure S4C.** Complete ribosomes detected in TNTs between PANC-1 cells by the RPS9-RPL24 proximity ligation assay *.

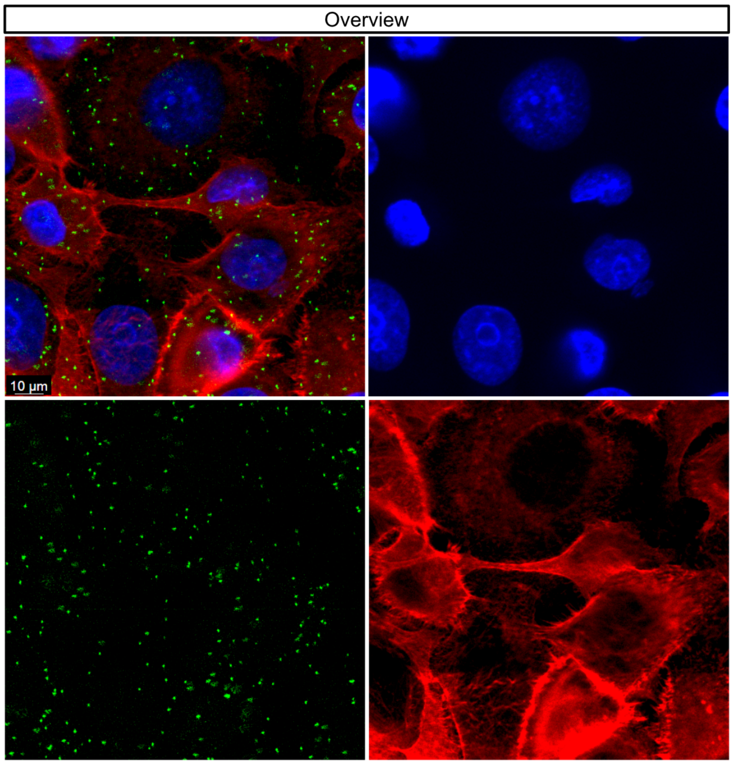


Overlay and channels (DNA – blue, RPS9-RPL24 proximity ligation – green, actin – red)

### FIGURE S4D

**Figure S4D.** HaloTag-RPS9-RPL24 proximity ligation assay, showing the incorporation of the labelled RPS9 into complete ribosomes in PANC-1 cells *.

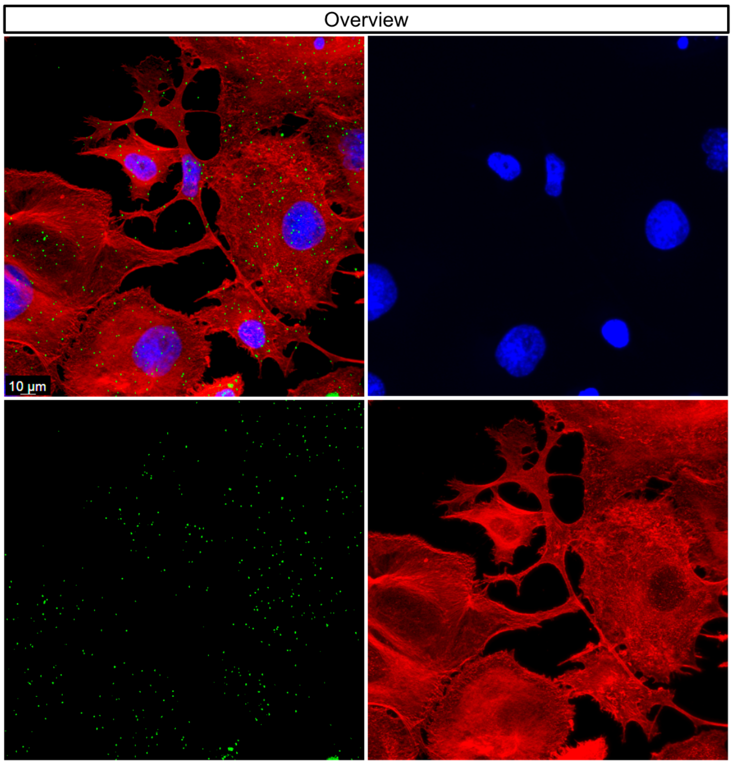


Overlay and channels (DNA – blue, HaloTag-RPS9-RPL24 proximity ligation – green, actin– red)

### FIGURE S4E

**Figure S4E.** Incorporation of HaloTag-RPS9 from donor cells into ribosomes in acceptor cells (a, b, c, d) * and negative controls without the HaloTag-RPS9 transgene (e, f) *.

**a.** Microscopic area 1

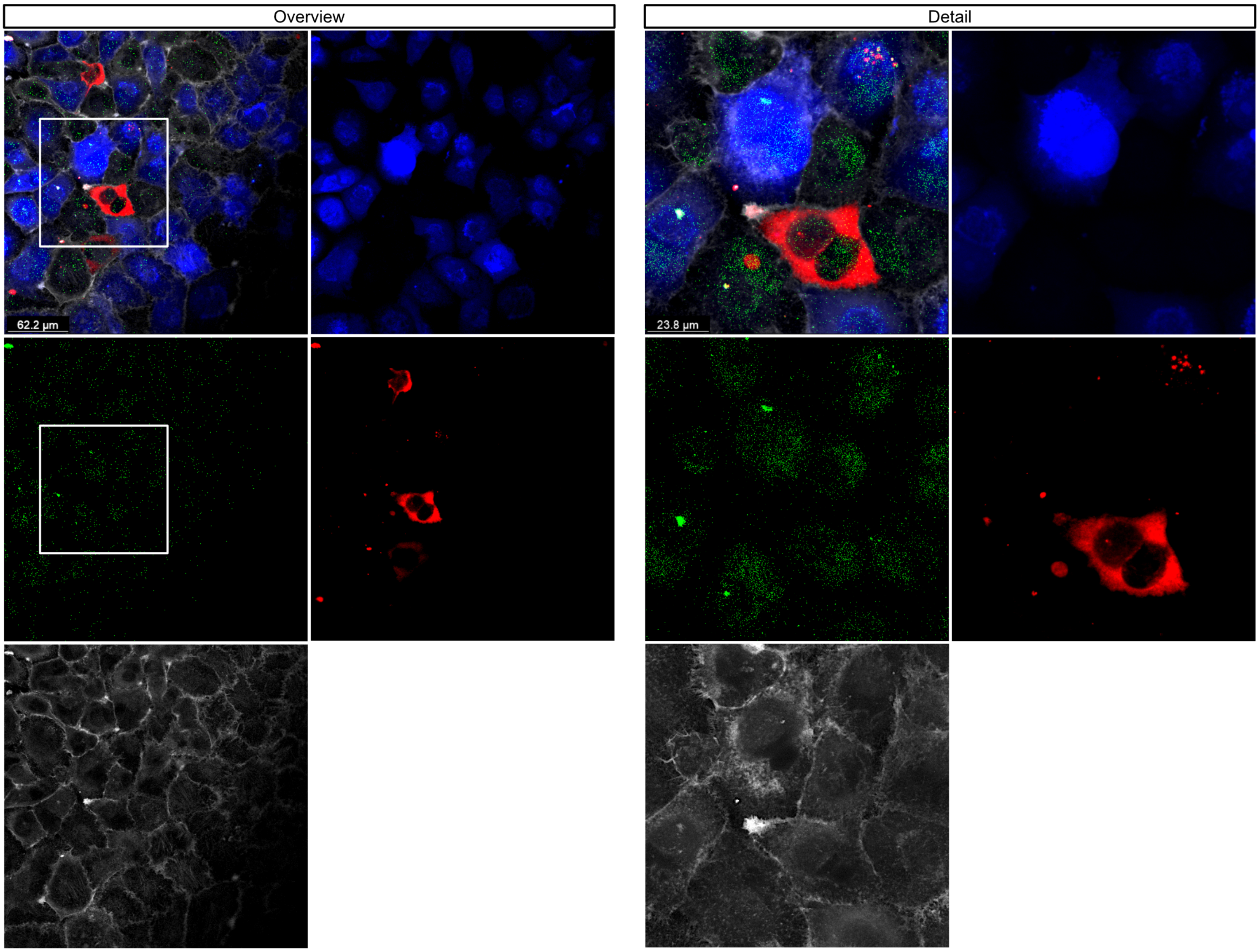


Overlay and channels (DNA – blue, HaloTag-RPS9-RPL24 proximity ligation– green, HaloTag-TMR ligand – red, actin – white)

### FIGURE S4E cont.

**Figure S4E.** Incorporation of HaloTag-RPS9 from donor cells into ribosomes in acceptor cells (a, b, c, d) * and negative controls without the HaloTag-RPS9 transgene (e, f) *.

**b.** Microscopic area 2

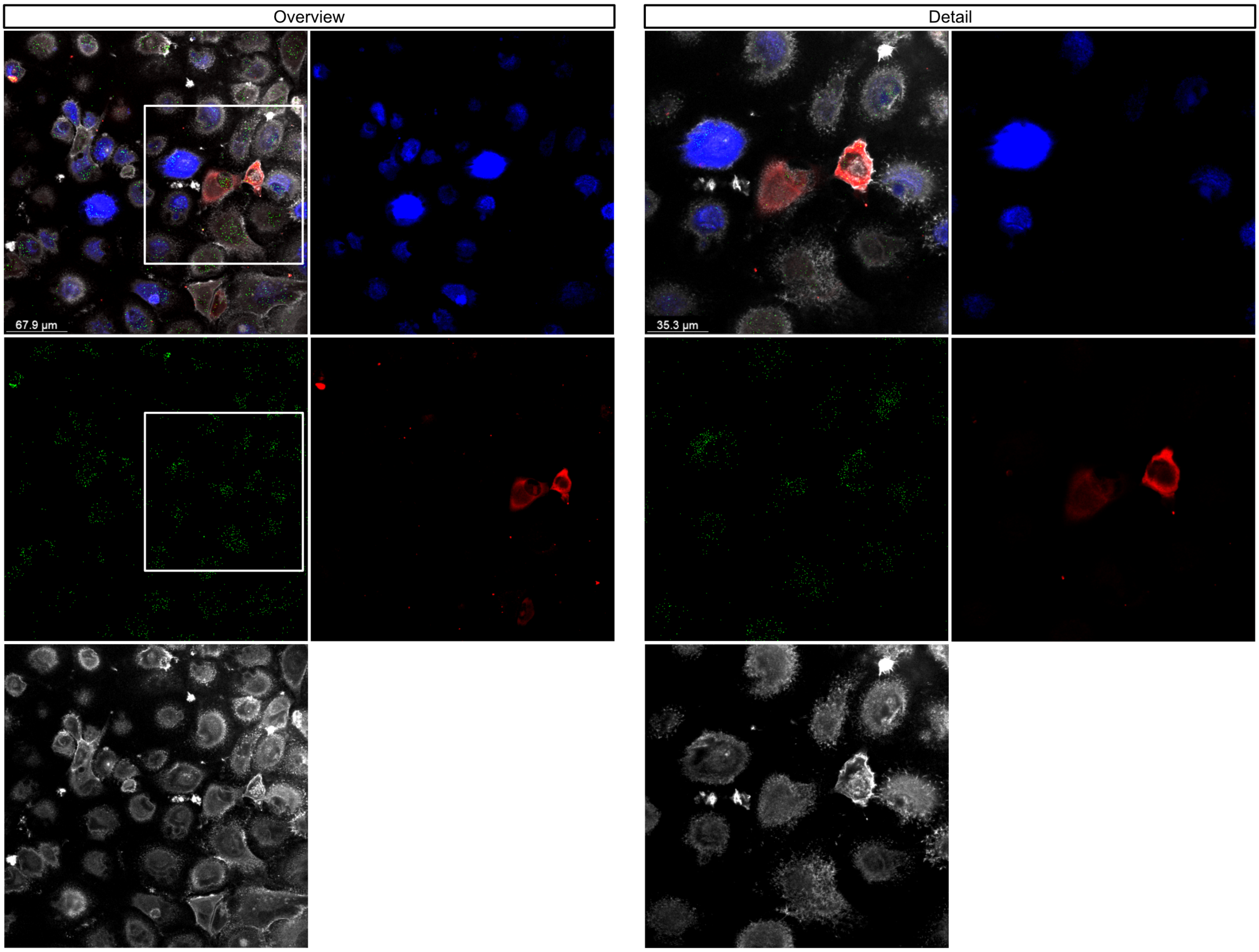


Overlay and channels (DNA – blue, HaloTag-RPS9-RPL24 proximity ligation– green, HaloTag-TMR ligand – red, actin – white)

### FIGURE S4E cont.

**Figure S4E.** Incorporation of HaloTag-RPS9 from donor cells into ribosomes in acceptor cells (a, b, c, d) * and negative controls without the HaloTag-RPS9 transgene (e, f) *.

**c.** Microscopic area 3

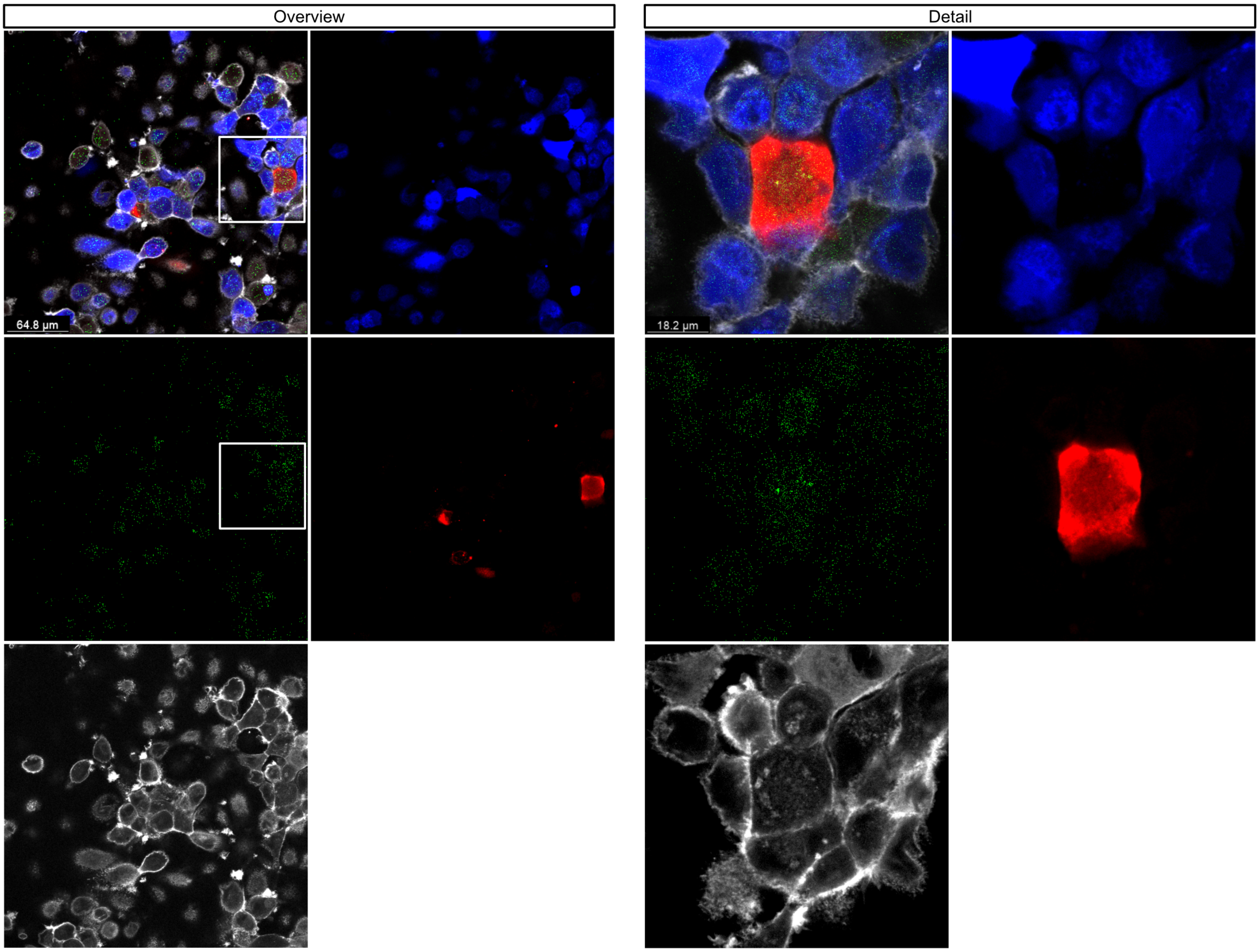


Overlay and channels (DNA – blue, HaloTag-RPS9-RPL24 proximity ligation– green, HaloTag-TMR ligand – red, actin – white)

### FIGURE S4E cont.

**Figure S4E.** Incorporation of HaloTag-RPS9 from donor cells into ribosomes in acceptor cells (a, b, c, d) * and negative controls without the HaloTag-RPS9 transgene (e, f) *.

**d.** Microscopic area 4

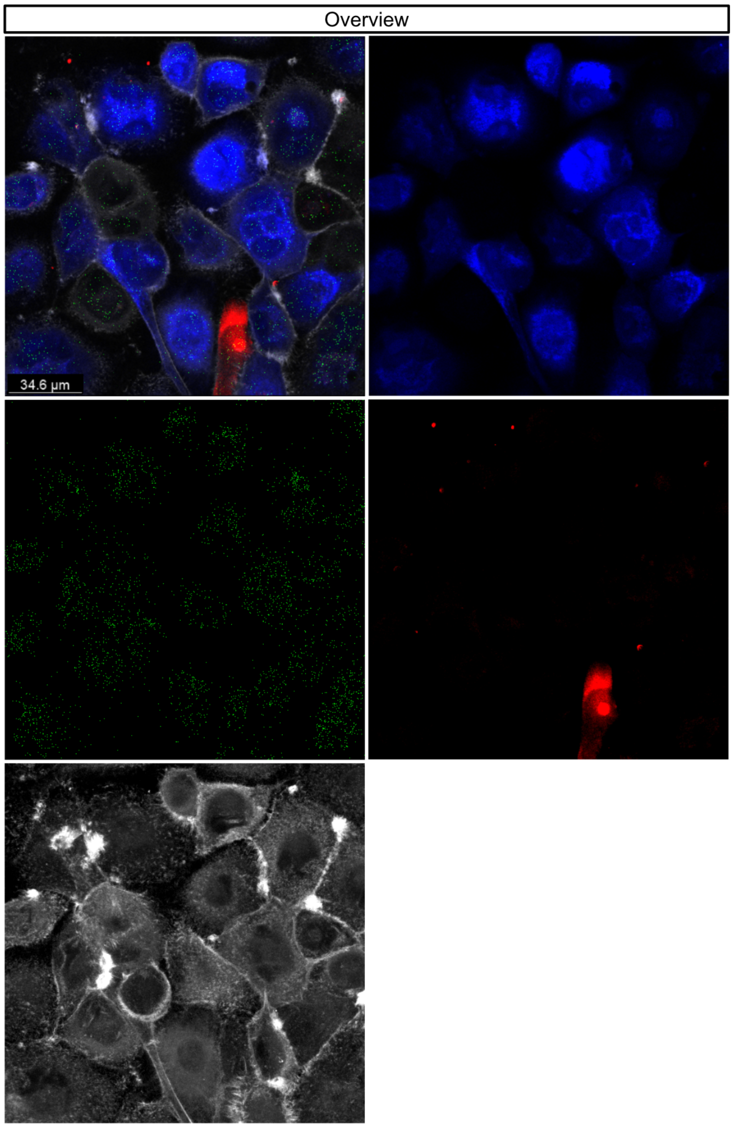


Overlay and channels (DNA – blue, HaloTag-RPS9-RPL24 proximity ligation– green, HaloTag-TMR ligand – red, actin – white)

### FIGURE S4E cont.

**Figure S4E.** Incorporation of HaloTag-RPS9 from donor cells into ribosomes in acceptor cells (a, b, c, d) * and negative controls without the HaloTag-RPS9 transgene (e, f) *.

**e.** Microscopic negative control area 1

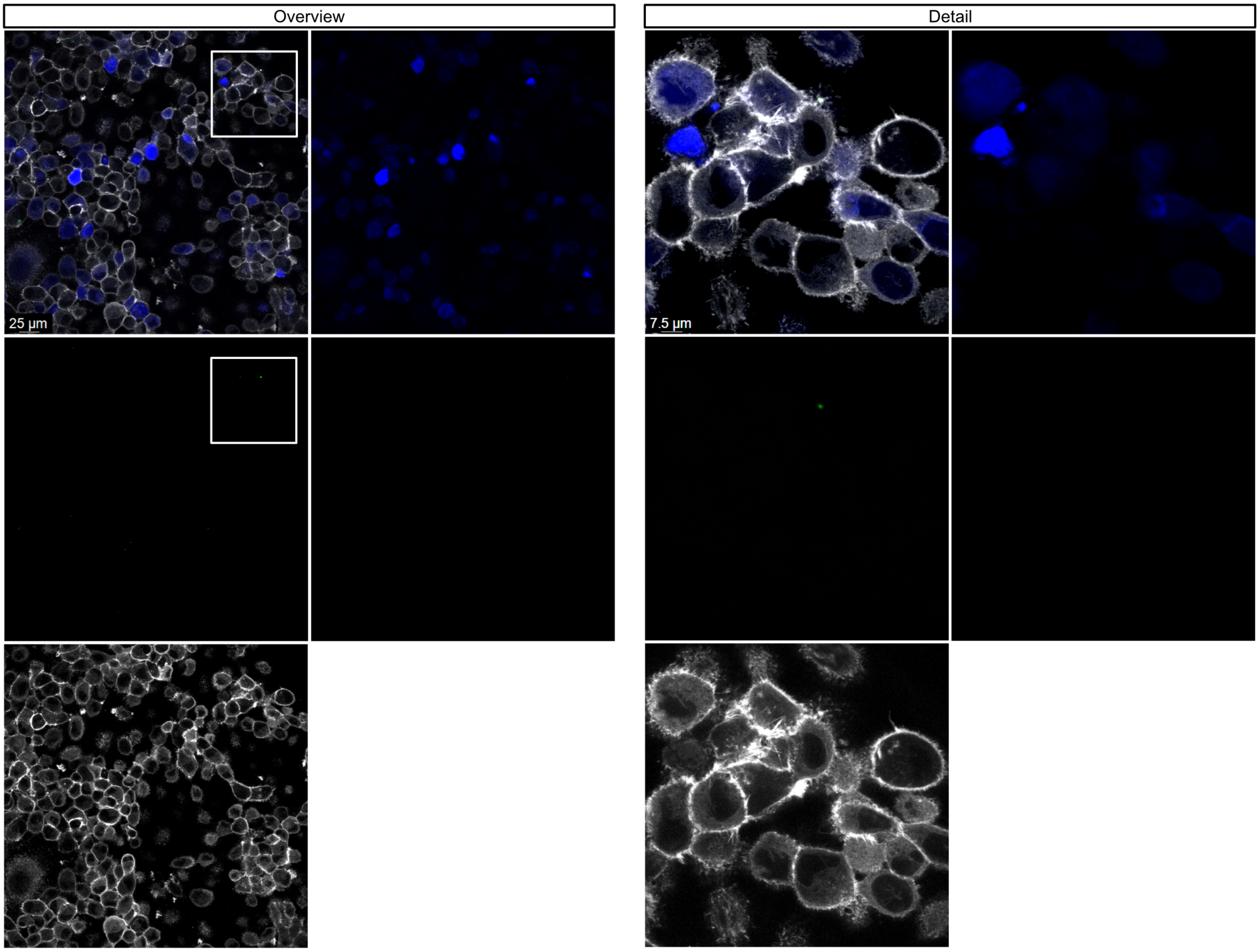


Overlay and channels (DNA – blue, HaloTag-RPS9-RPL24 proximity ligation– green, HaloTag-TMR ligand – red, actin – white)

### FIGURE S4E cont.

**Figure S4E.** Incorporation of HaloTag-RPS9 from donor cells into ribosomes in acceptor cells (a, b, c, d) * and negative controls without the HaloTag-RPS9 transgene (e, f) *.

**f.** Microscopic negative control area 2

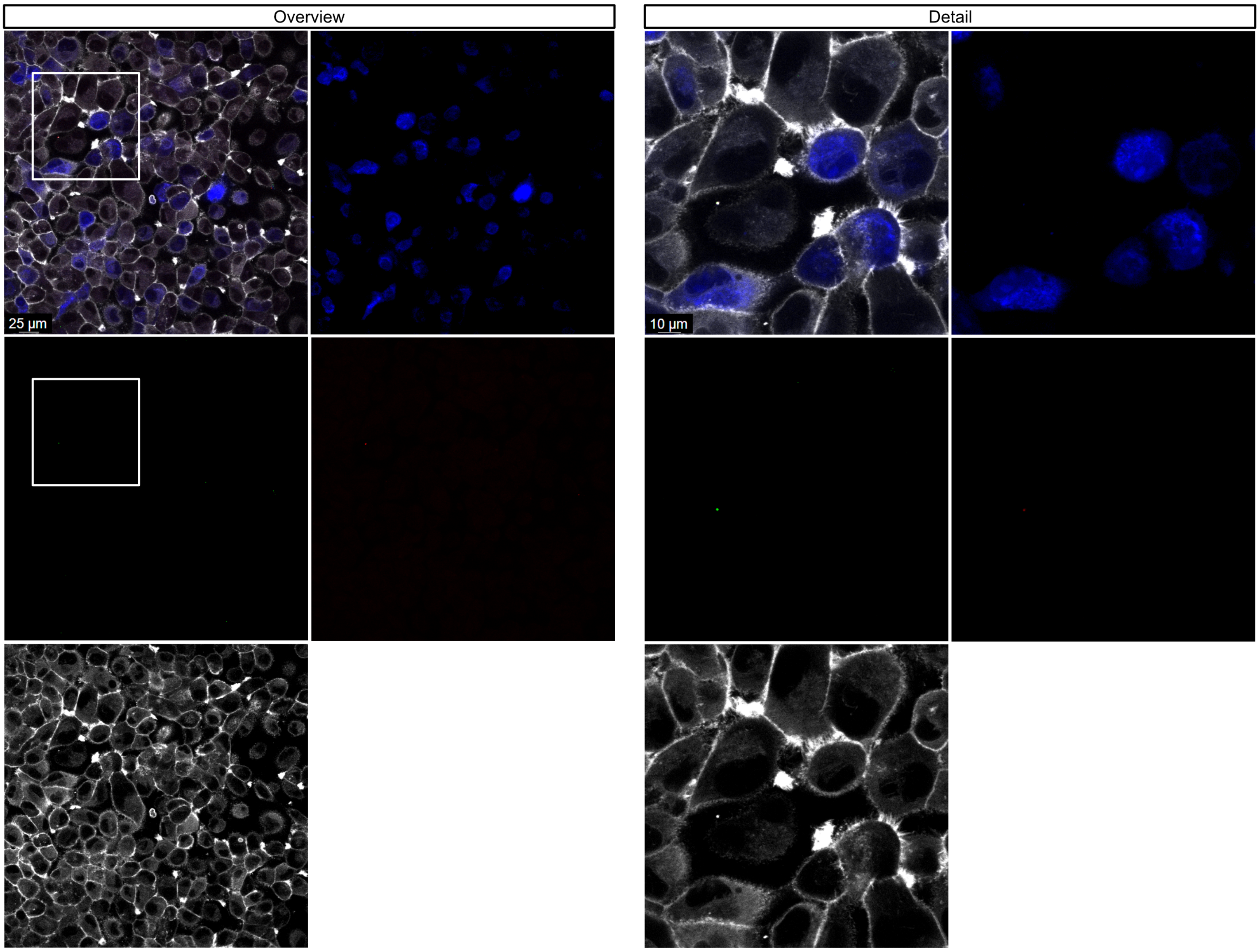


Overlay and channels (DNA – blue, HaloTag-RPS9-RPL24 proximity ligation– green, HaloTag-TMR ligand – red, actin – white)

### FIGURE S4F

**Figure S4F.** Double positive events  HaloTag TMR + (PE-A) and CellTrace Violet + (DAPI-A) and singlets gate sensitivity analysis for each of the three mixed population experiments. Untreated experiments (a, b, c) and gemcitabine treated experiments (d, e, f).

**a.** Untreated first experiment

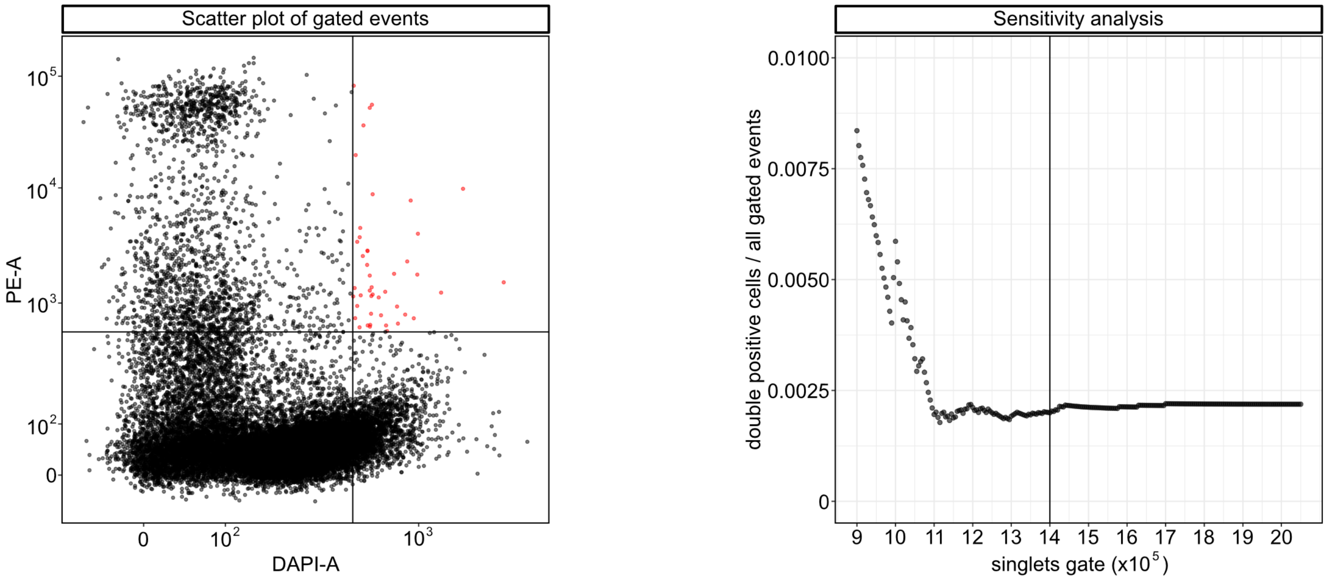


**b.** Untreated second experiment

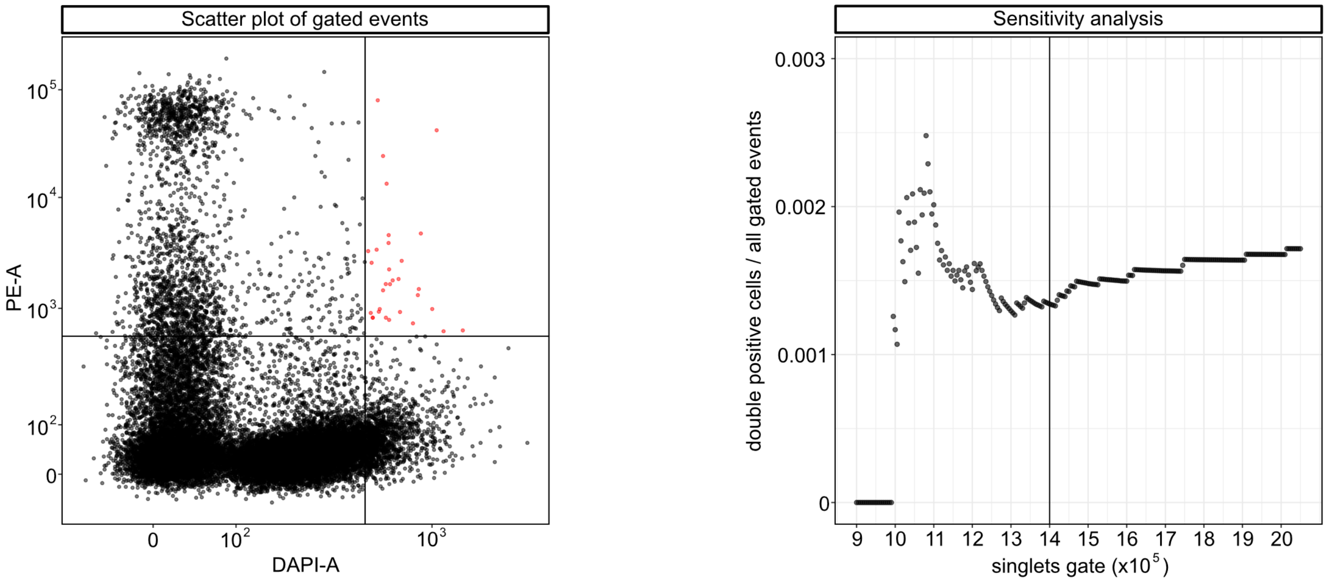


**c.** Untreated third experiment

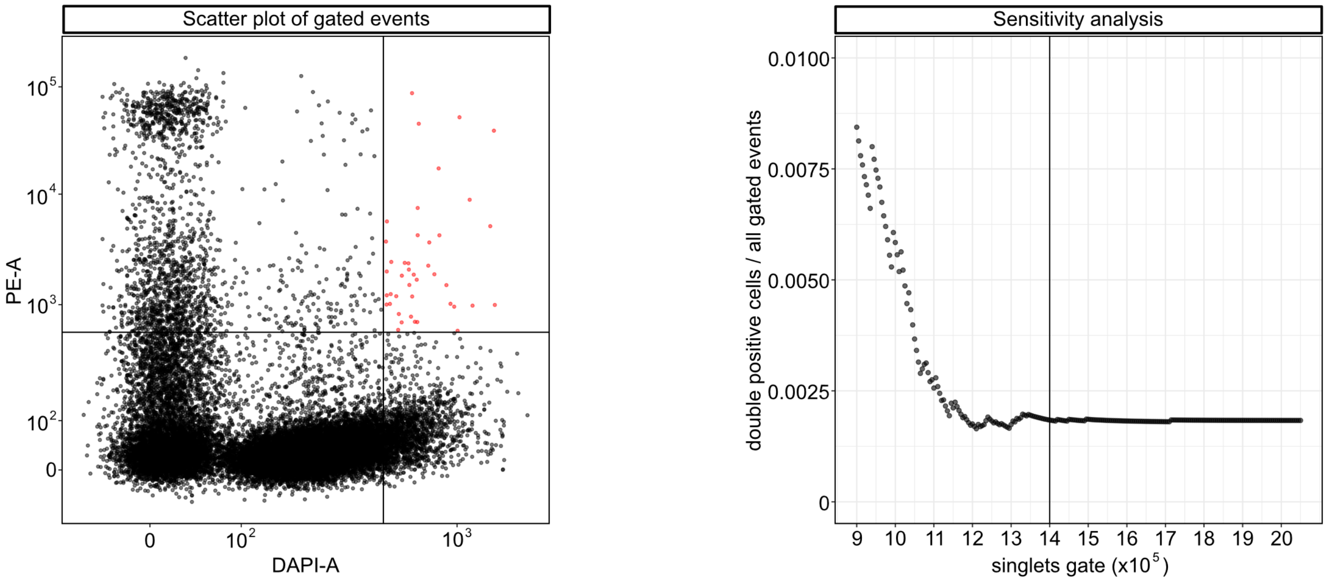

### FIGURE S4F cont.

**Figure S4F.** Doubly positive events  HaloTag TMR (PE-A) and CellTrace Violet (DAPI-A) and singlets gate sensitivity analysis for each of the three mixed population experiments. Untreated experiments (a, b, c) and gemcitabine treated experiments (d, e, f).

**d.** Gemcitabine treated first experiment

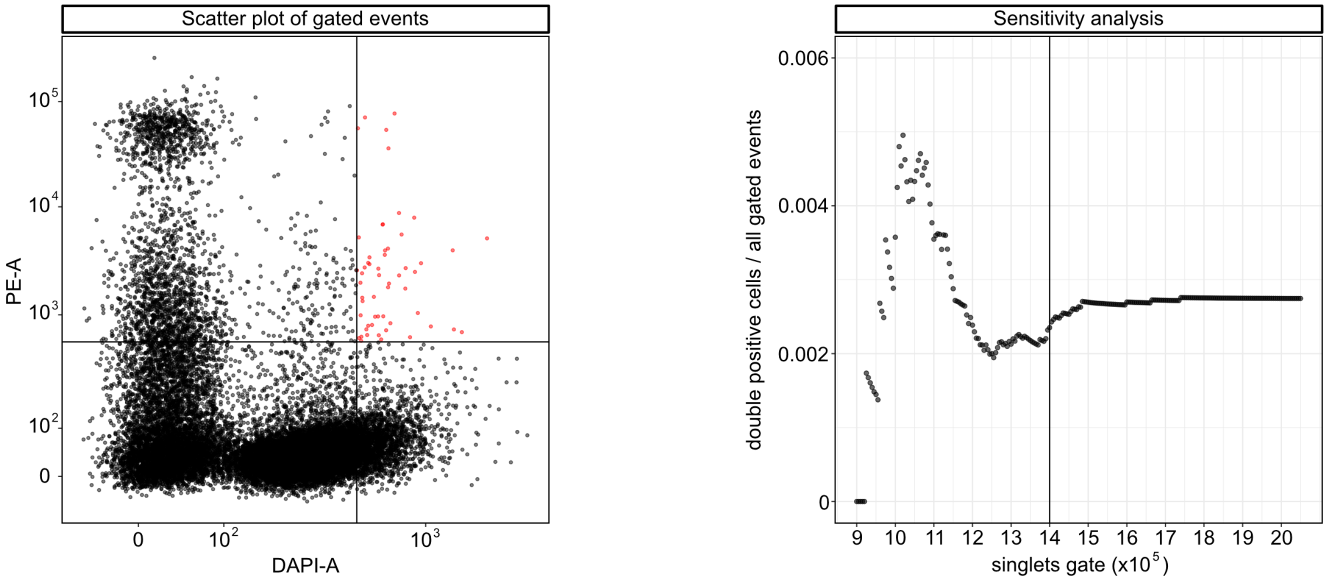


**e.** Gemcitabine treated second experiment

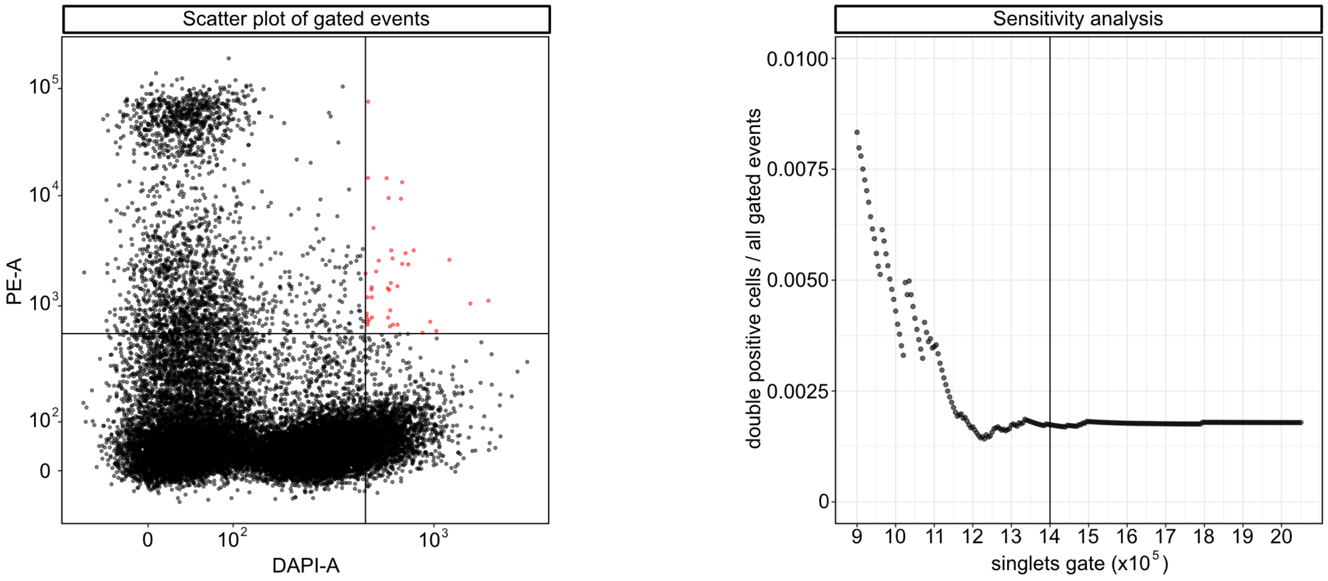


**f.** Gemcitabine treated third experiment

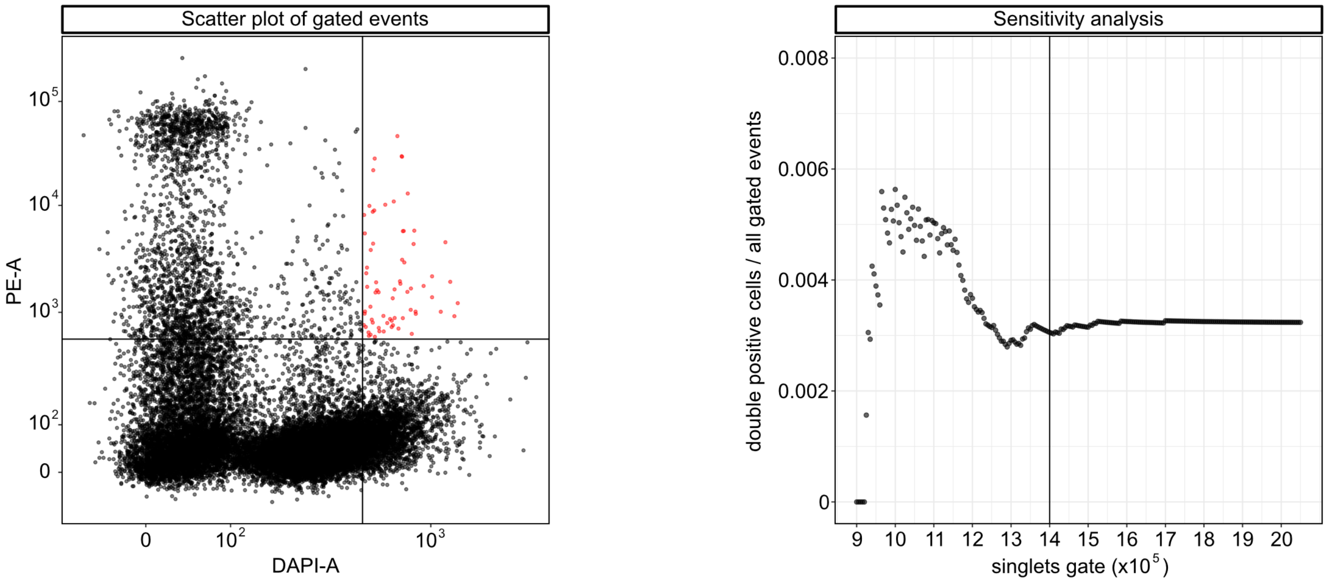

### FIGURE S5A

**Figure S5A.** siRNA-mediated knockdown of ribosomal proteins S6 and L24. Western blot analysis reveals reduced protein levels of RPS6 and RPL24 following siRNA-mediated knockdown of their corresponding transcripts. Protein lysates were collected at 24, 48, and 72 hours post-treatment. Expression levels were quantified using Bio-Rad Image Lab software, normalised to total protein, and compared to control samples collected at the same time points (a, b, c).

**a.** Western blot first experiment

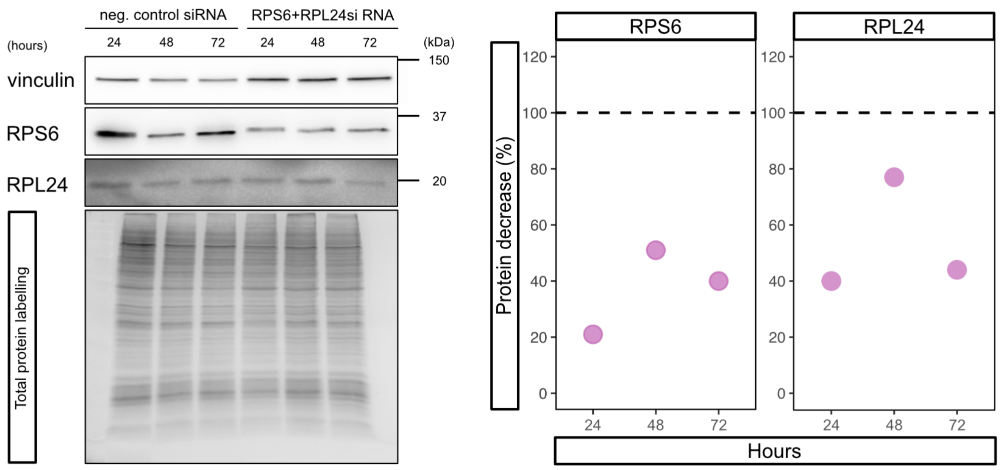


**b.** Western blot second experiment

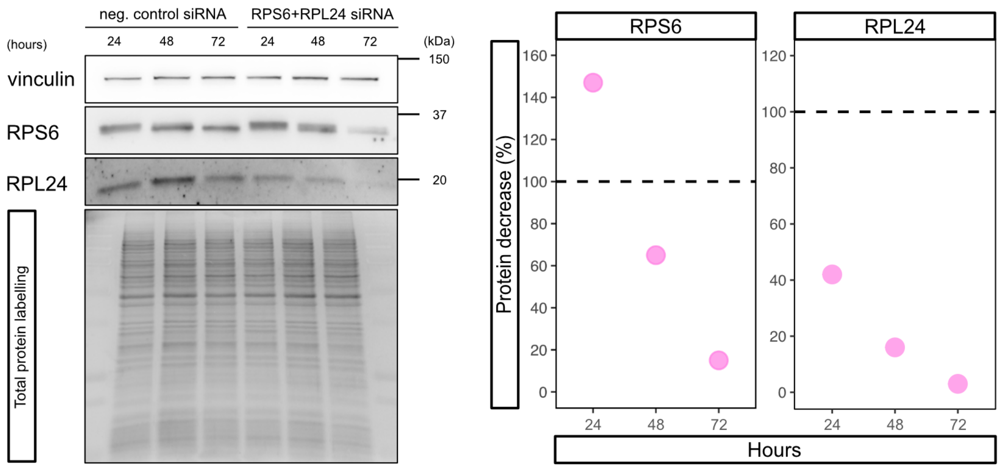


**c.** Western blot third experiment

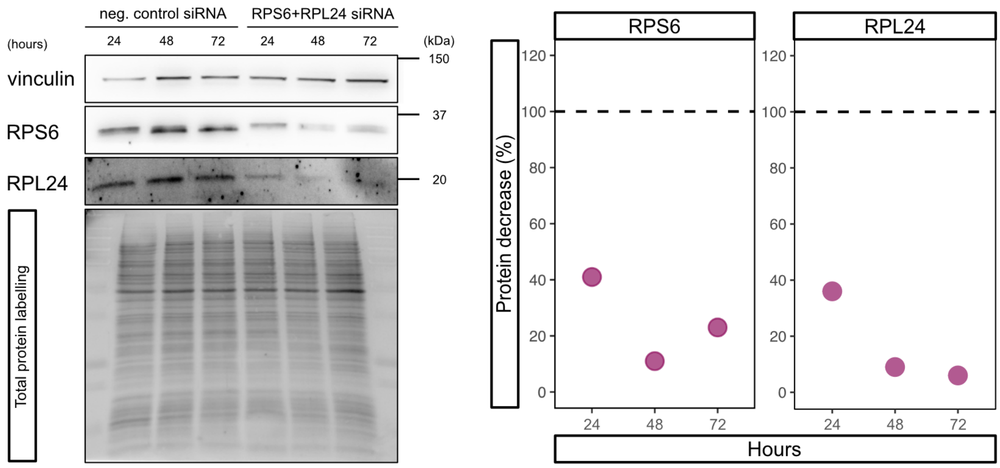

### FIGURE S5D

**Figure S5D.** Comparison of overall protein synthesis of control cells stained with CFSE, non-stained siRNA cells and mixed control (CFSE) & siRNA cells (non-stained). The histogram depict slight CFSE transfer from control cells to siRNA cells via a TNT. The third plot represents a comparison of overall protein synthesis, using strict gate settings to exclude partially stained cells. Scatter plots show the fluorescence intensity of translation (Alexa 647-H) and CFSE staining (FITC-H) (a, b, c).

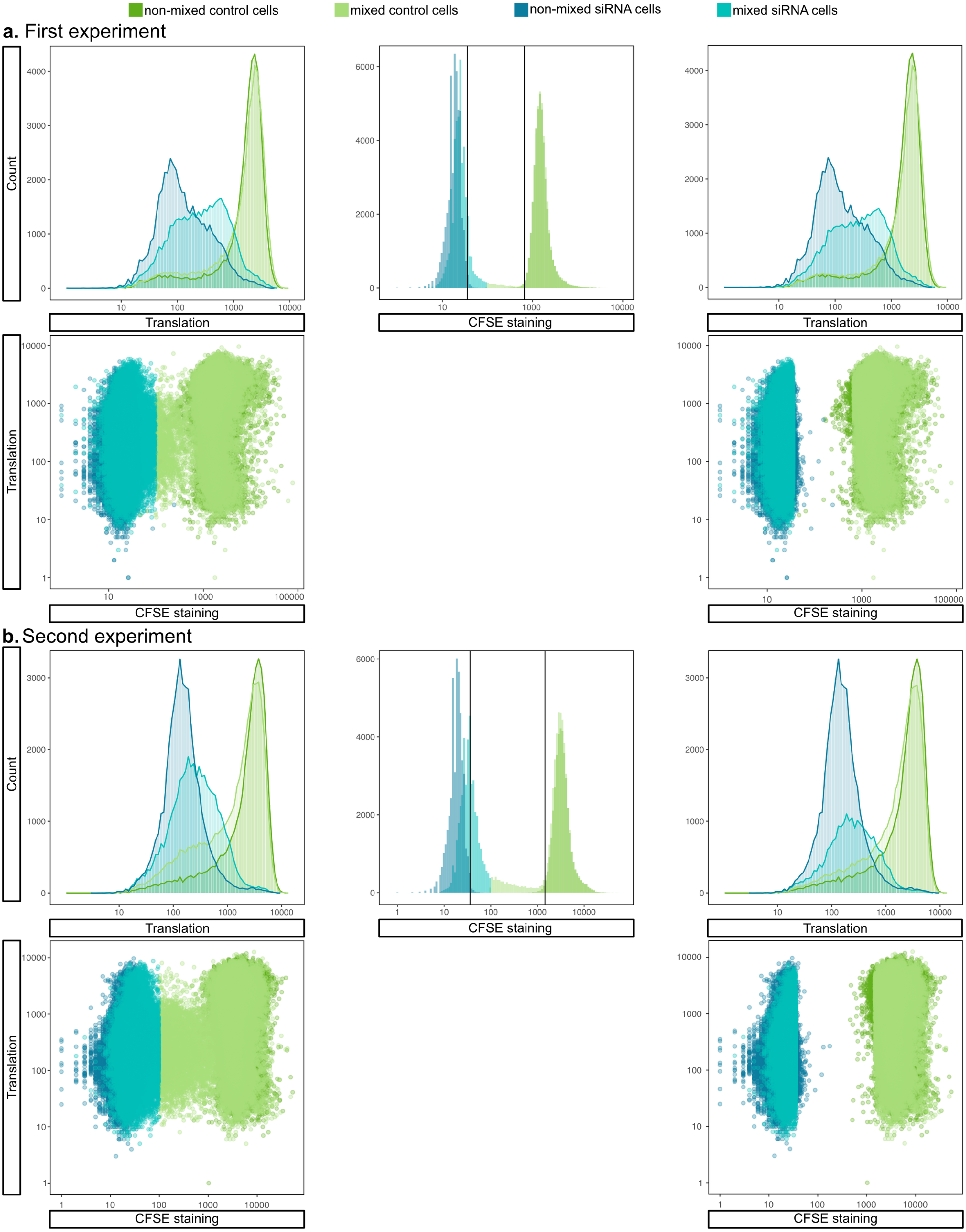

### FIGURE S5D cont.

**Figure S5D cont.** Comparison of overall protein synthesis of control cells stained with CFSE, non-stained siRNA cells and mixed control (CFSE) & siRNA cells (non-stained). The histogram depict slight CFSE transfer from control cells to siRNA cells via a TNT. The third plot represents a comparison of overall protein synthesis, using strict gate settings to exclude partially stained cells. Scatter plots show the fluorescence intensity of translation (Alexa 647-H) and CFSE staining (FITC-H) (a, b, c).

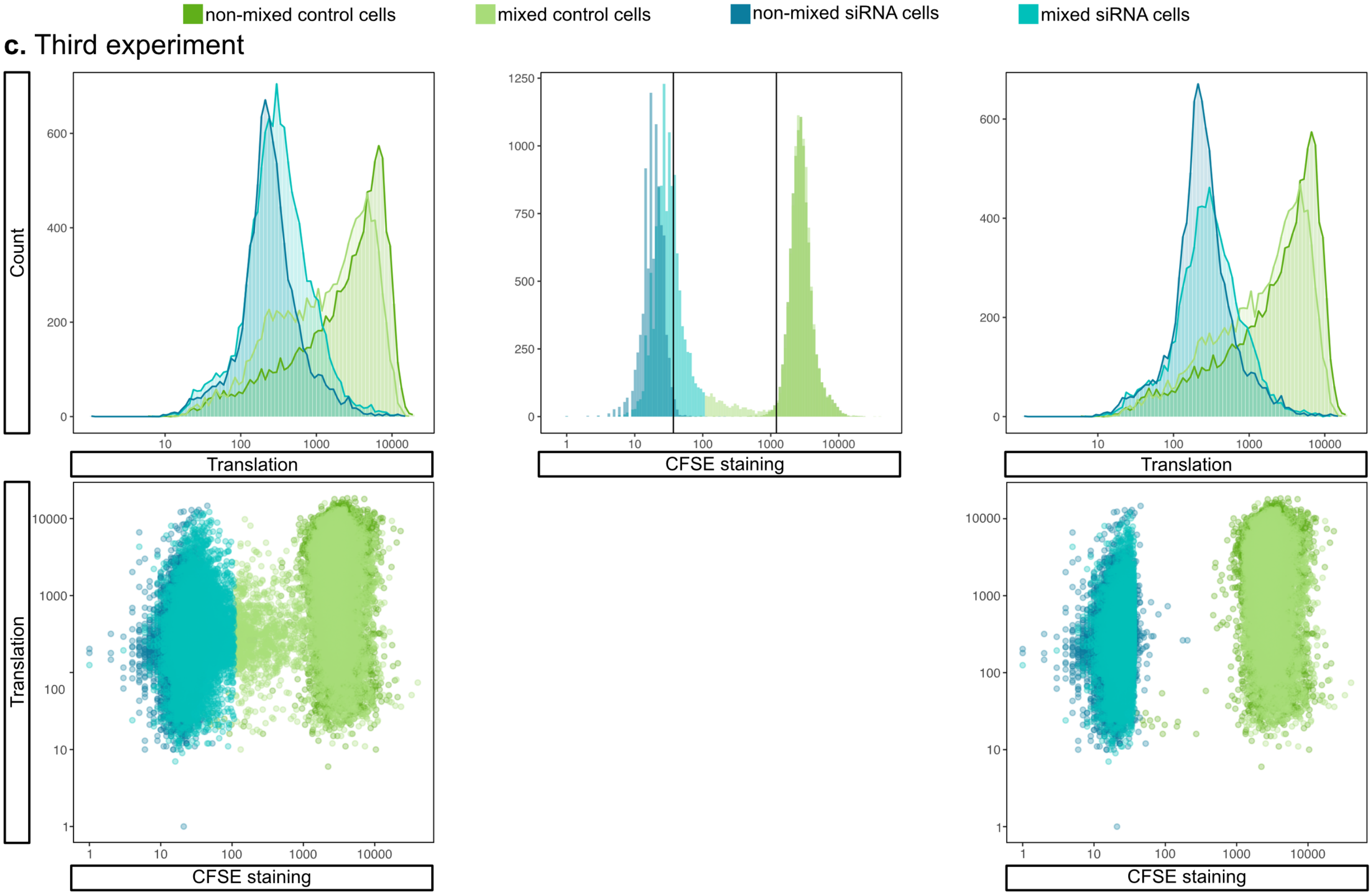

### FIGURE S5E

**Figure S5E.** Comparison of overall protein synthesis of non-stained control cells, siRNA cells stained with CFSE and mixed control (non-stained) & siRNA cells (CFSE) in colour-reversed experiment. The histogram represents slight CFSE transfer from siRNA cells to control cells. The third plot represents a comparison of overall protein synthesis, using strict gate settings to exclude partially stained cells. Scatter plots show the fluorescence intensity of translation (Alexa 647-H) and CFSE staining (FITC-H) (a, b, c).

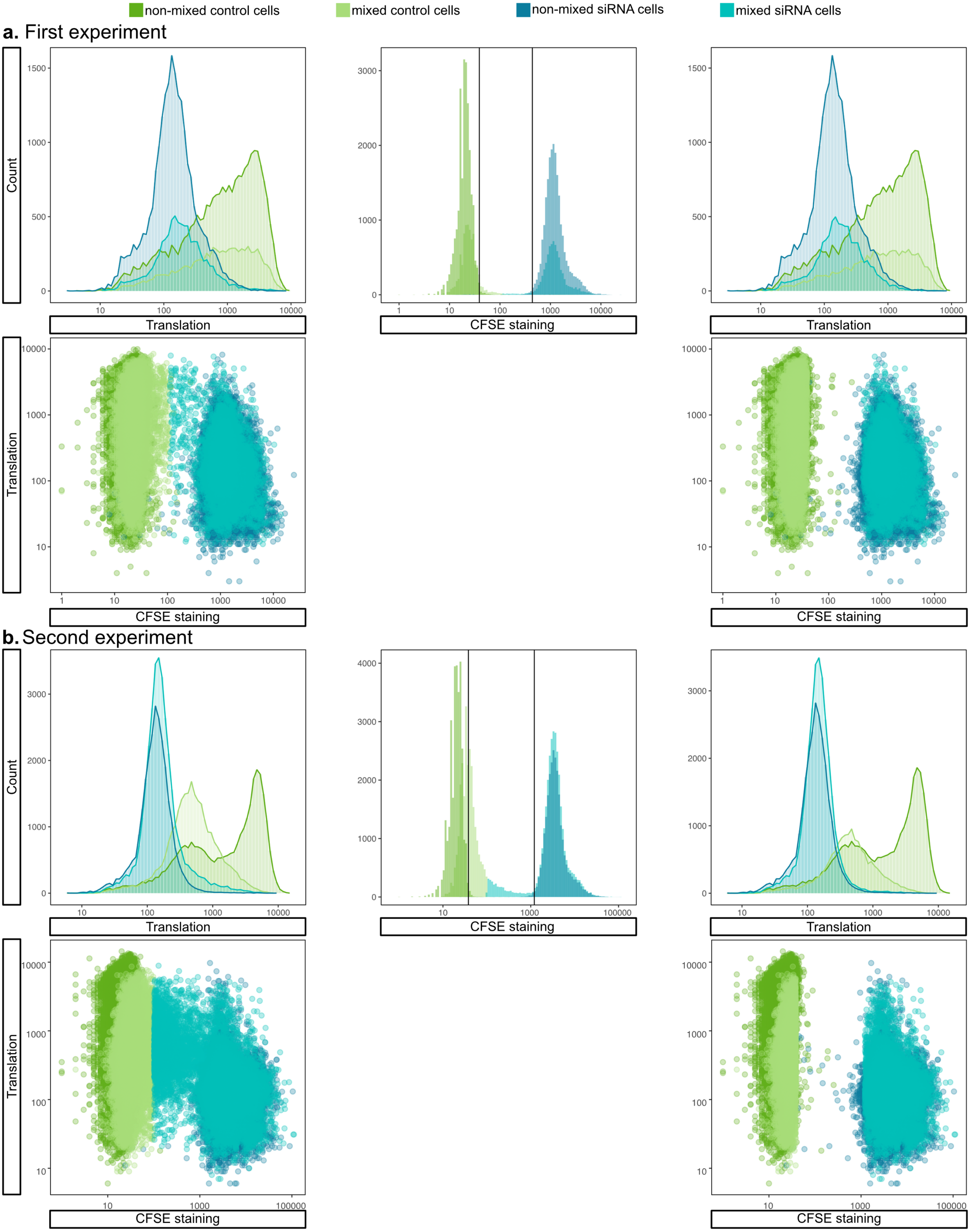

### FIGURE S5E cont.

**Figure S5E cont.** Comparison of overall protein synthesis of non-stained control cells, siRNA cells stained with CFSE and mixed control (non-stained) & siRNA cells (CFSE) in colour-reversed experiment. The histogram represents slight CFSE transfer from siRNA cells to control cells. The third plot represents a comparison of overall protein synthesis, using strict gate settings to exclude partially stained cells. Scatter plots show the fluorescence intensity of translation (Alexa 647-H) and CFSE staining (FITC-H) (a, b, c).

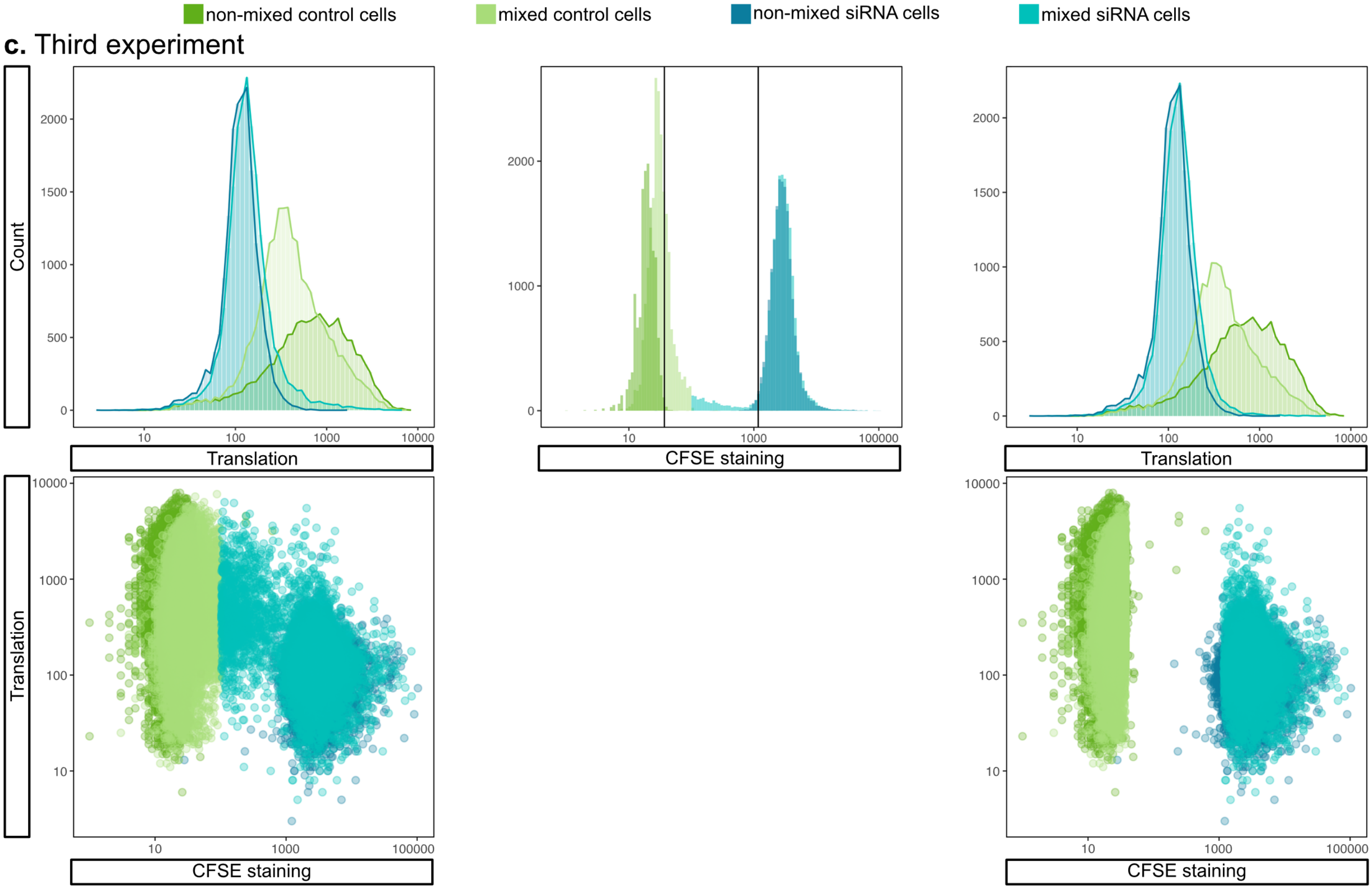

### FIGURE S5F

**Figure S5F.** Bayesian model for posterior predictions of overall protein synthesis of control cells stained with CFSE, non-stained siRNA cells and mixed control (CFSE) & siRNA cells (non-stained) using one gate (a). Colour-reversed experiment: Bayesian model for posterior predictions of overall protein synthesis of control cells (non-stained), siRNA cells (CFSE) and mixed control (non-stained) & siRNA cells (CFSE) using one gate (b) and the stricter two gates strategy (c).

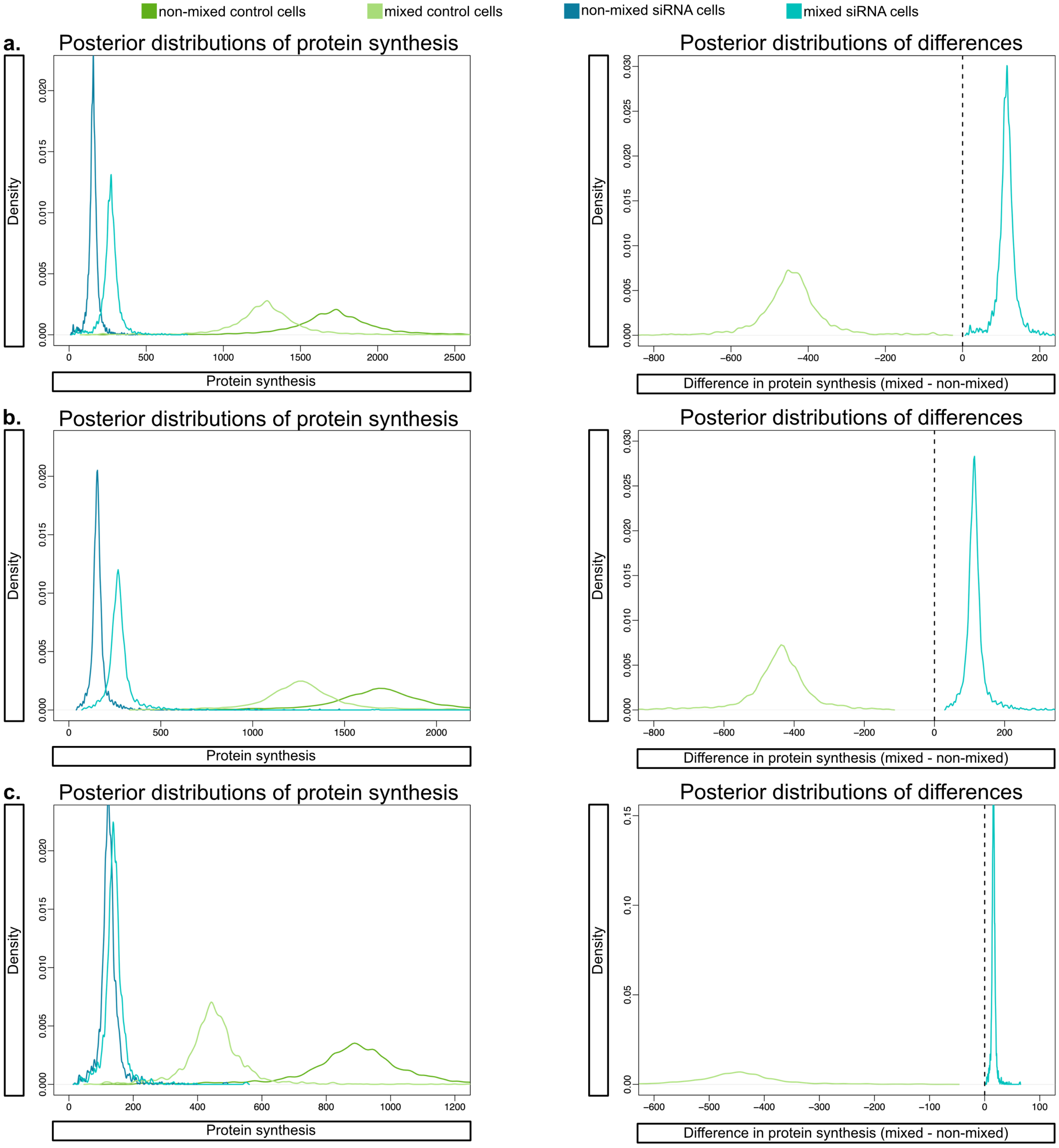

### FIGURE S6.

**Figure S6.** Microscopic quantification of assembled ribosomes using a proximity ligation assay: **A.** PANC-1 cells co-treated with siRNA-RPS6 + siRNA-RPL24 (a, b, c, d) **B.** mixed co-culture experimental group (a, b, c, d, e, f, g, h) **C.** siRNA-negative control PANC-1 cells (a, b, c, d). ROIs of PANC-1 cells co-treated with siRNA-RPS6 + siRNA-RPL24 (blue ROIs), siRNA-negative control PANC-1 cells (green ROIs). The fourth quadrant of each overview shows the ribosomal spot detection and counting strategy within the ROI.

**A.** Microscopic fields with cells co-treated with siRNA-RPS6 + siRNA-RPL24 (a, b, c, d)

**a.**

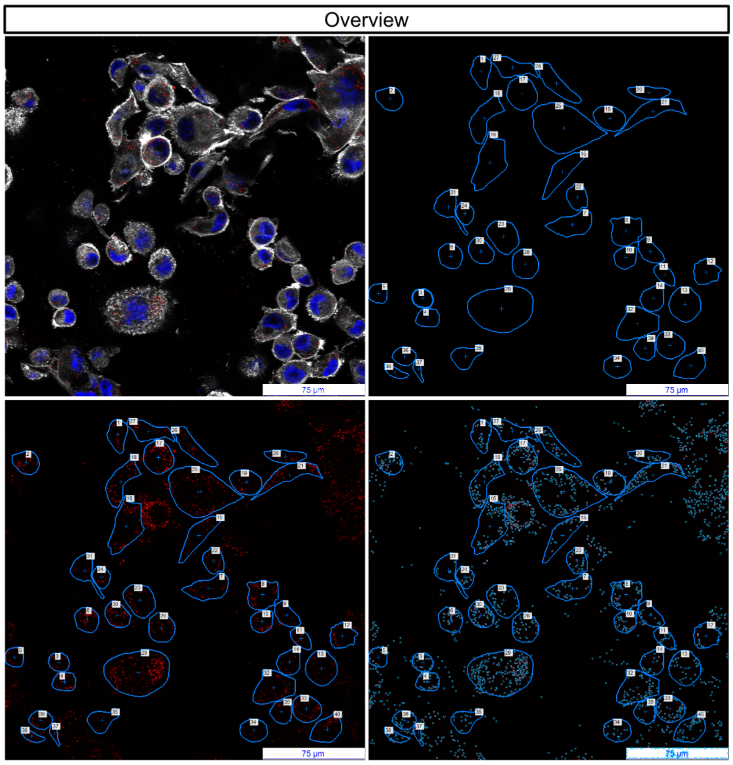


**b.**

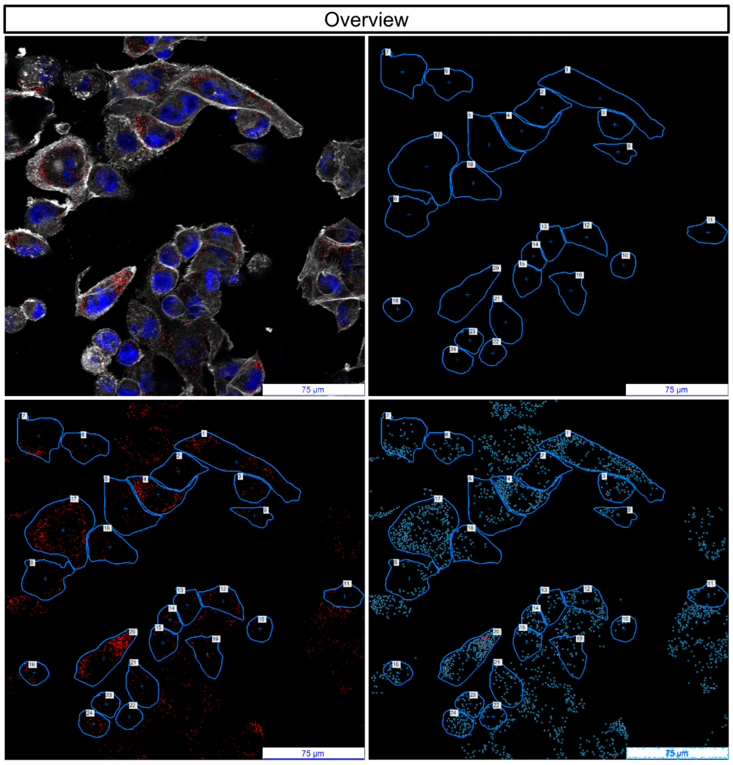


Overlay and channels (DNA – blue, CFSE – green, RPS6+RPL24 proximity ligation – red, actin – white)

**c.**

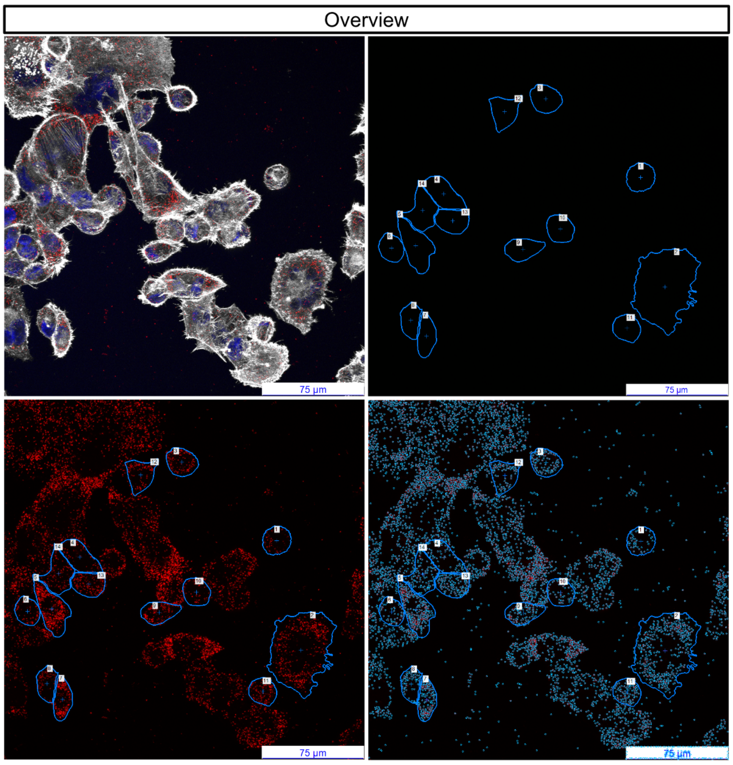


**d.**

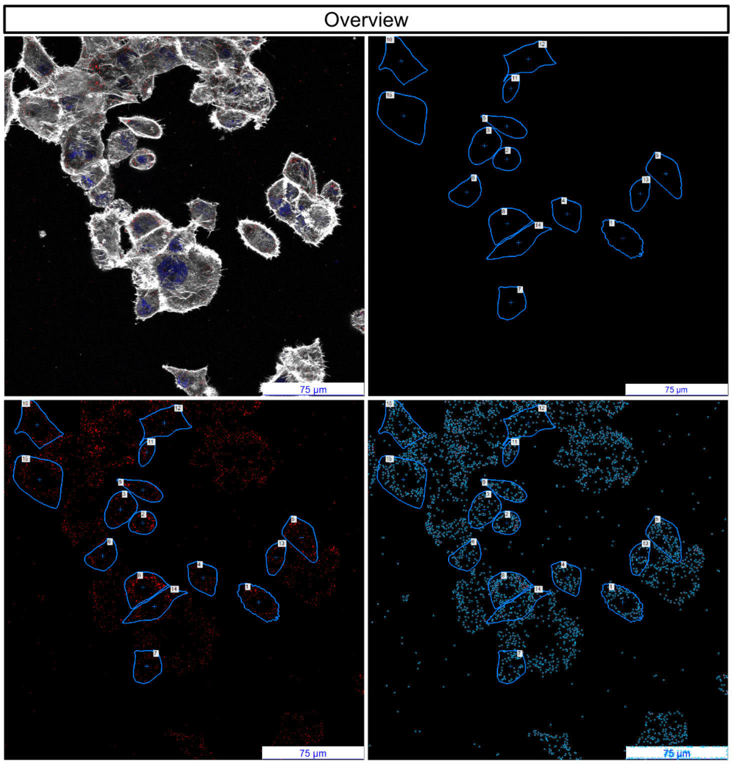


Overlay and channels (DNA – blue, CFSE – green, RPS6+RPL24 proximity ligation – red, actin – white)

#### FIGURE S6 cont.

**Figure S6.** Microscopic quantification of assembled ribosomes using a proximity ligation assay: **A.** PANC-1 cells co-treated with siRNA-RPS6 + siRNA-RPL24 (a, b, c, d) **B.** mixed co-culture experimental group (a, b, c, d, e, f, g, h) **C.** siRNA-negative control PANC-1 cells (a, b, c, d). ROIs of PANC-1 cells co-treated with siRNA-RPS6 + siRNA-RPL24 (blue ROIs), siRNA-negative control PANC-1 cells (green ROIs). The fourth quadrant of each overview shows the ribosomal spot detection and counting strategy within the ROI.

**B.** Microscopic fields with mixed co-culture experimental group (a, b, c, d, e, f, g, h)

**a.**

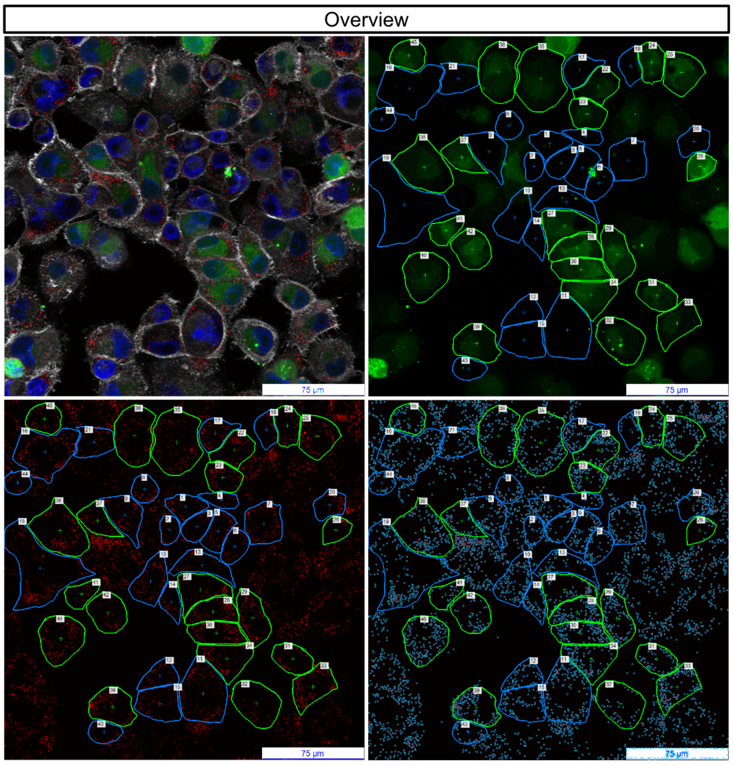


**b.**

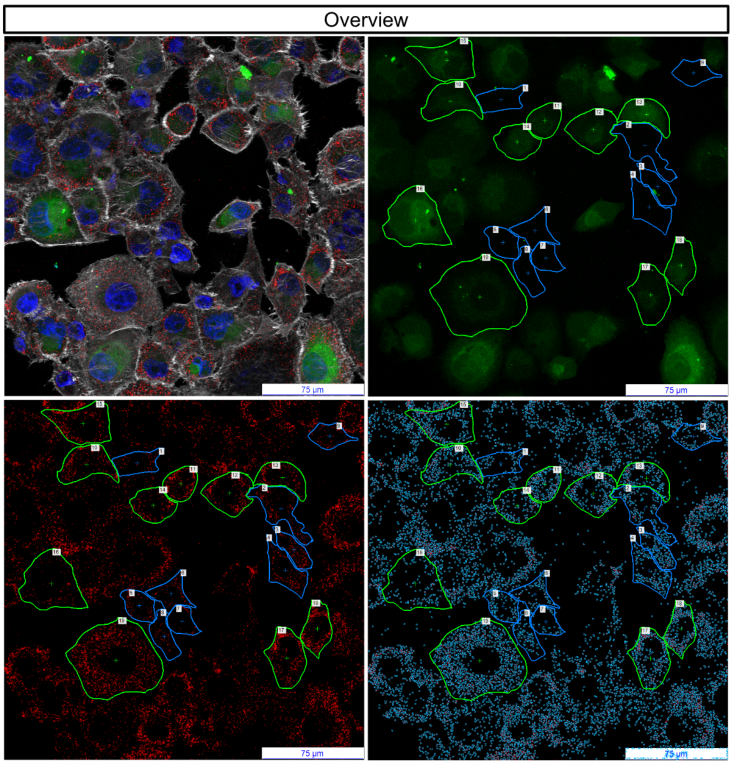


Overlay and channels (DNA – blue, CFSE – green, RPS6+RPL24 proximity ligation – red, actin – white)

**c.**

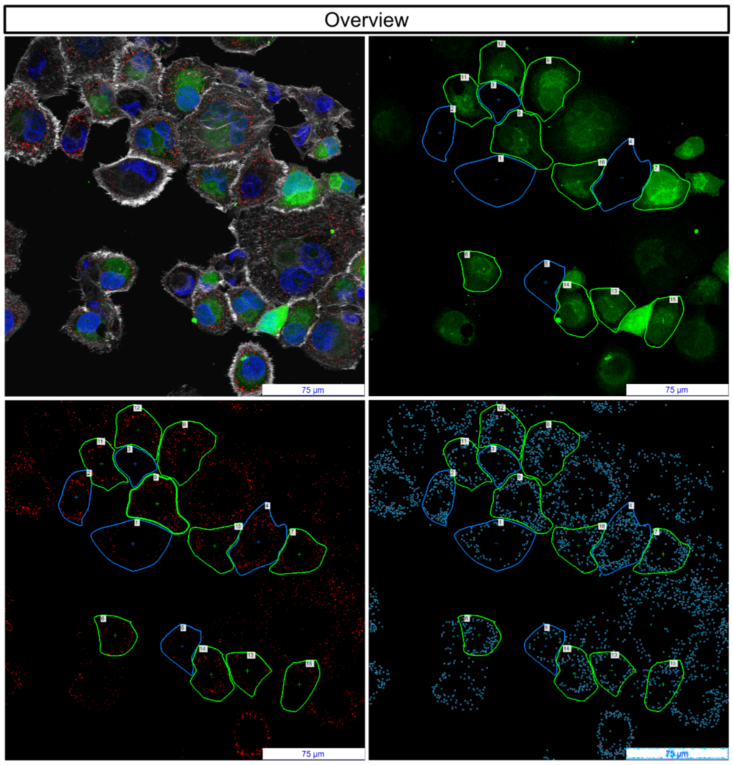


**d.**

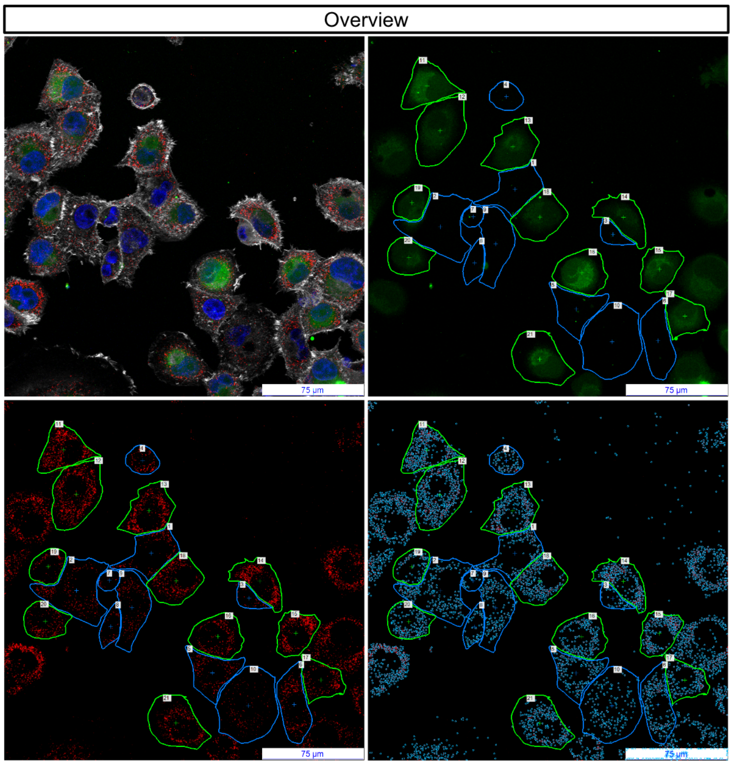


Overlay and channels (DNA – blue, CFSE – green, RPS6+RPL24 proximity ligation – red, actin – white)

FIGURE S6 cont.

**Figure S6.** Microscopic quantification of assembled ribosomes using a proximity ligation assay: **A.** PANC-1 cells co-treated with siRNA-RPS6 + siRNA-RPL24 (a, b, c, d) **B.** mixed co-culture experimental group (a, b, c, d, e, f, g, h) **C.** siRNA-negative control PANC-1 cells (a, b, c, d). ROIs of PANC-1 cells co-treated with siRNA-RPS6 + siRNA-RPL24 (blue ROIs), siRNA-negative control PANC-1 cells (green ROIs). The fourth quadrant of each overview shows the ribosomal spot detection and counting strategy within the ROI.

**B.** Microscopic fields with mixed co-culture experimental group (a, b, c, d, e, f, g, h)

**e.**

Overview

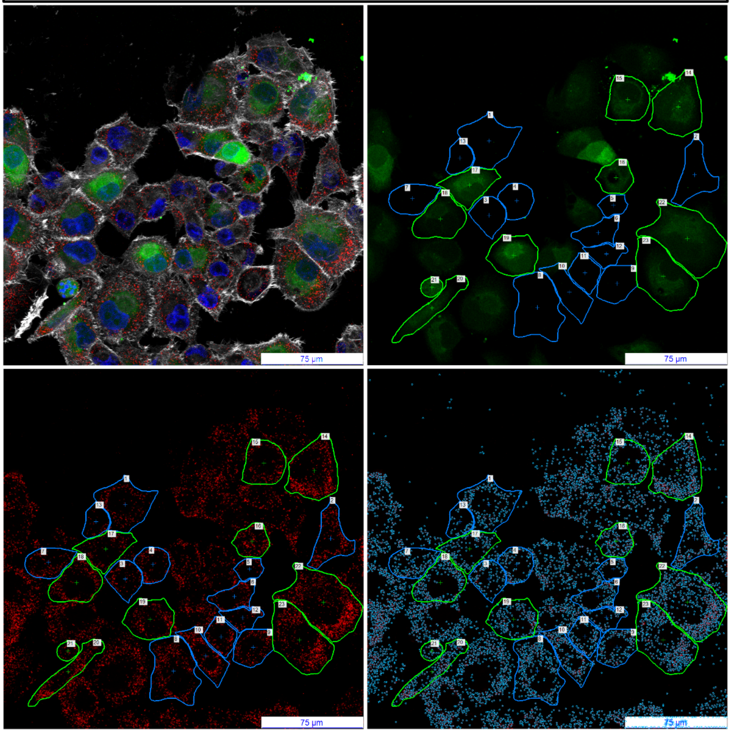

**f.**

Overview

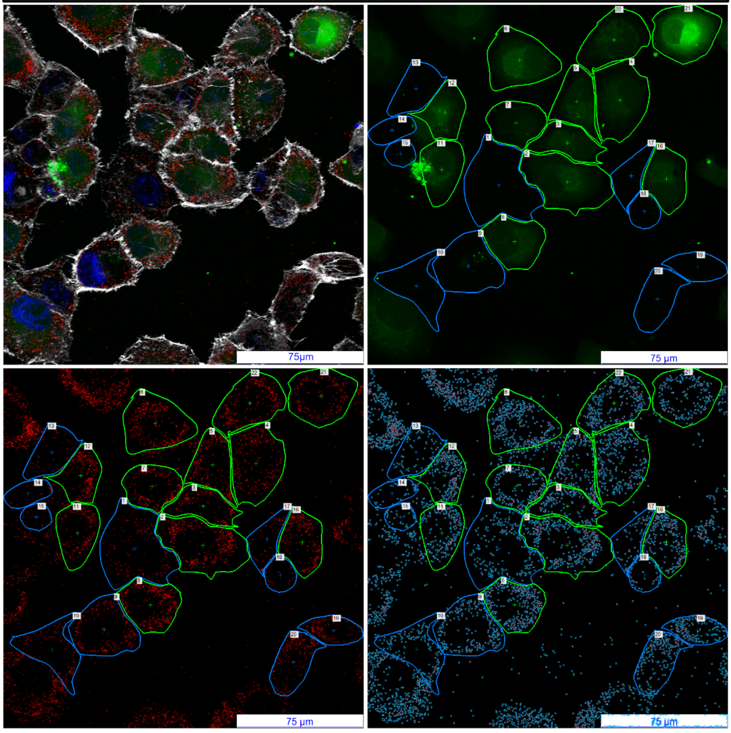

Overlay and channels (DNA – blue, CFSE – green, RPS6+RPL24 proximity ligation – red, actin – white)

**g.**

Overview

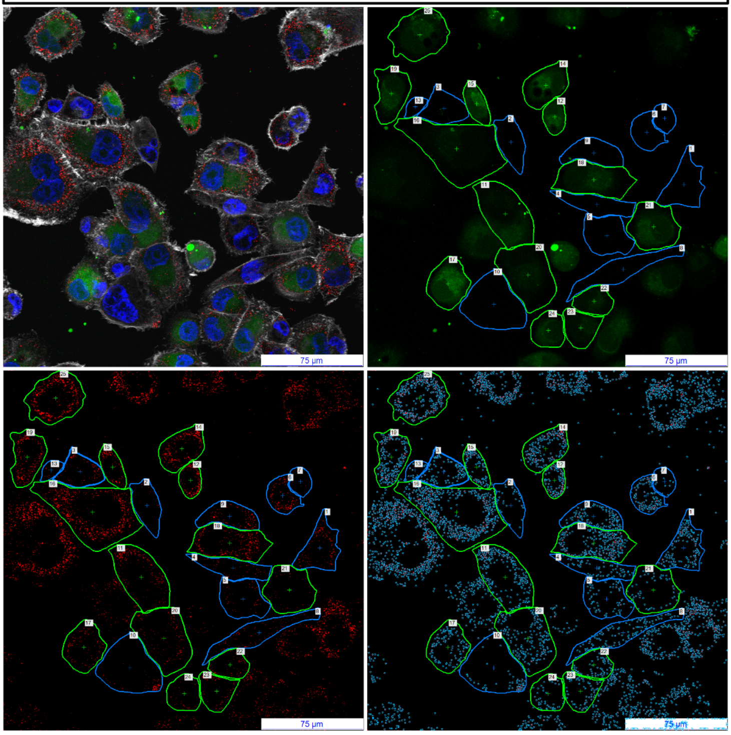

**h.**

Overview

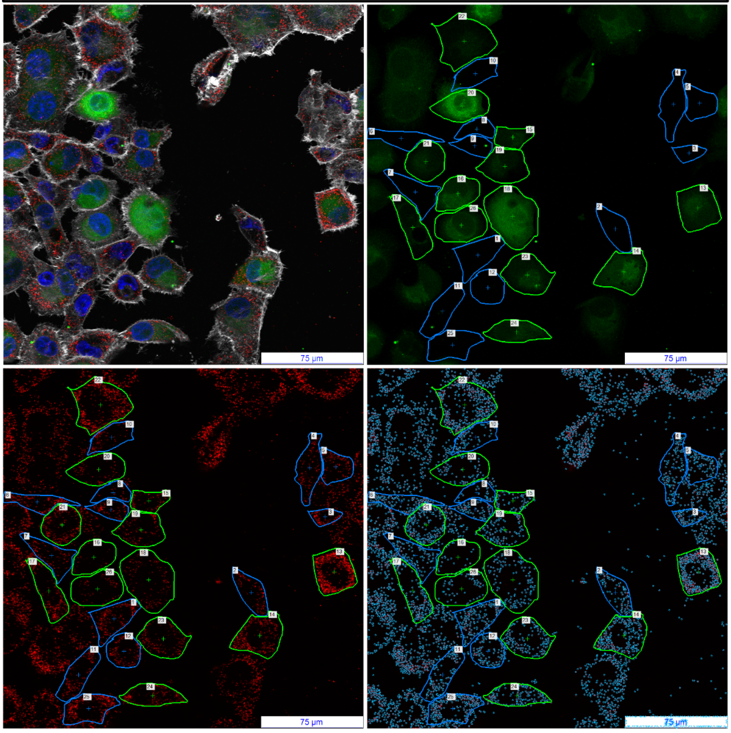

Overlay and channels (DNA – blue, CFSE – green, RPS6+RPL24 proximity ligation – red, actin – white)

#### FIGURE S6 cont.

**Figure S6.** Microscopic quantification of assembled ribosomes using a proximity ligation assay: **A.** PANC-1 cells co-treated with siRNA-RPS6 + siRNA-RPL24 (a, b, c, d) **B.** mixed co-culture experimental group (a, b, c, d, e, f, g, h) **C.** siRNA-negative control PANC-1 cells (a, b, c, d). ROIs of PANC-1 cells co-treated with siRNA-RPS6 + siRNA-RPL24 (blue ROIs), siRNA-negative control PANC-1 cells (green ROIs). The fourth quadrant of each overview shows the ribosomal spot detection and counting strategy within the ROI.

**C.** Microscopic fields with siRNA-negative control PANC-1 cells (a, b, c, d)

**a.**

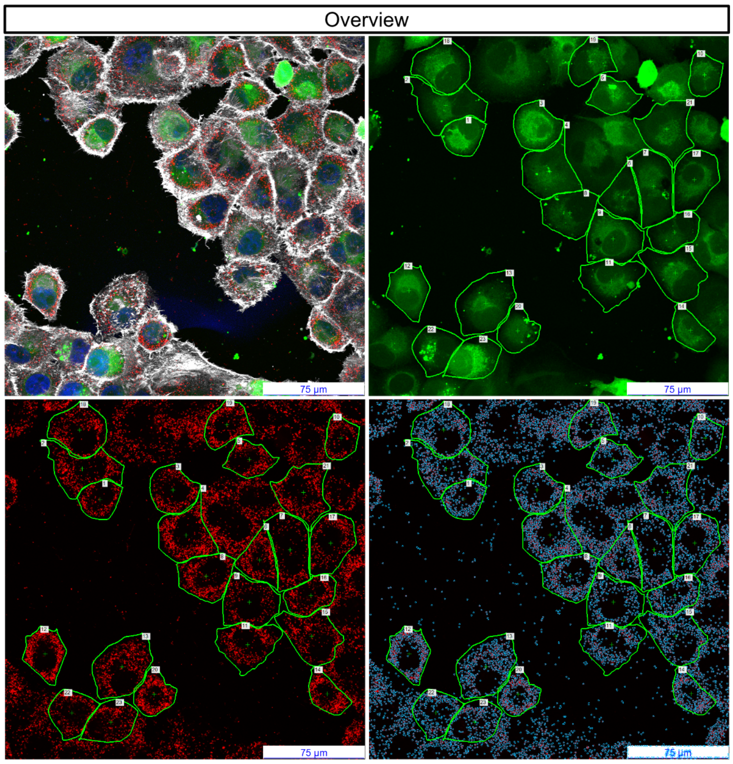


**b.**

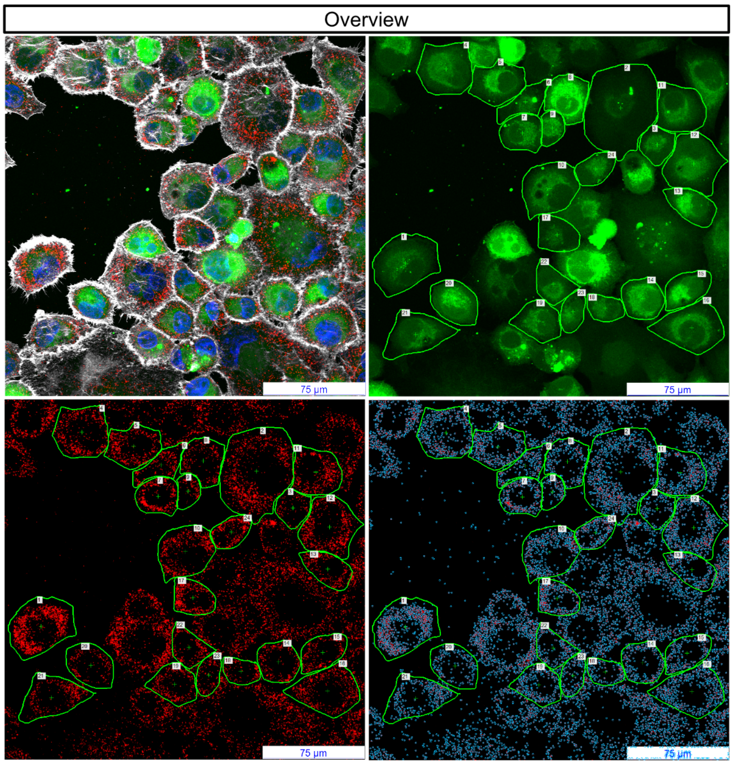


Overlay and channels (DNA – blue, CFSE – green, RPS6+RPL24 proximity ligation – red, actin – white)

**c.**

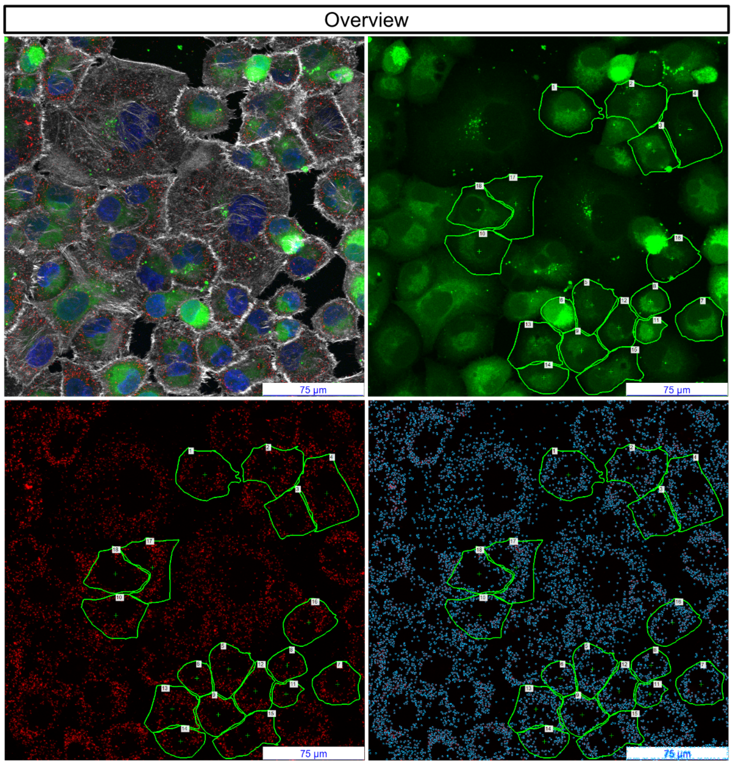


**d.**

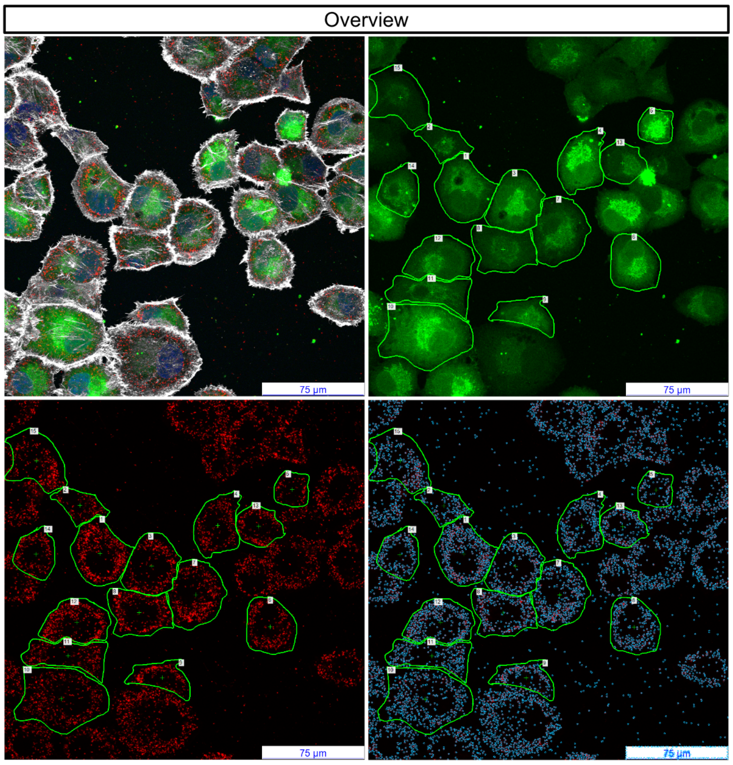


Overlay and channels (DNA – blue, CFSE – green, RPS6+RPL24 proximity ligation – red, actin – white)

### FIGURE S7A

**Figure S7A.** Ribosomal 5.8S rRNA detected in tunnelling nanotubes (a, b), RNase treated negative control stained for 5.8S rRNA (c, d) and no primary, secondary antibody control staining for 5.8S rRNA (e).

**a.** PANC-1 5.8S rRNA * without treatment

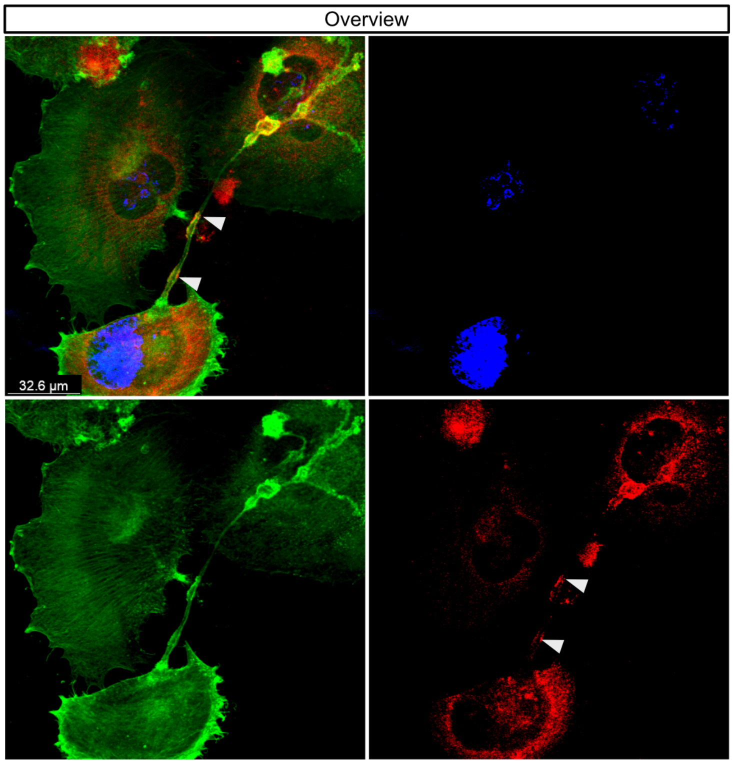


**b.** PANC-1 5.8S rRNA * without treatment

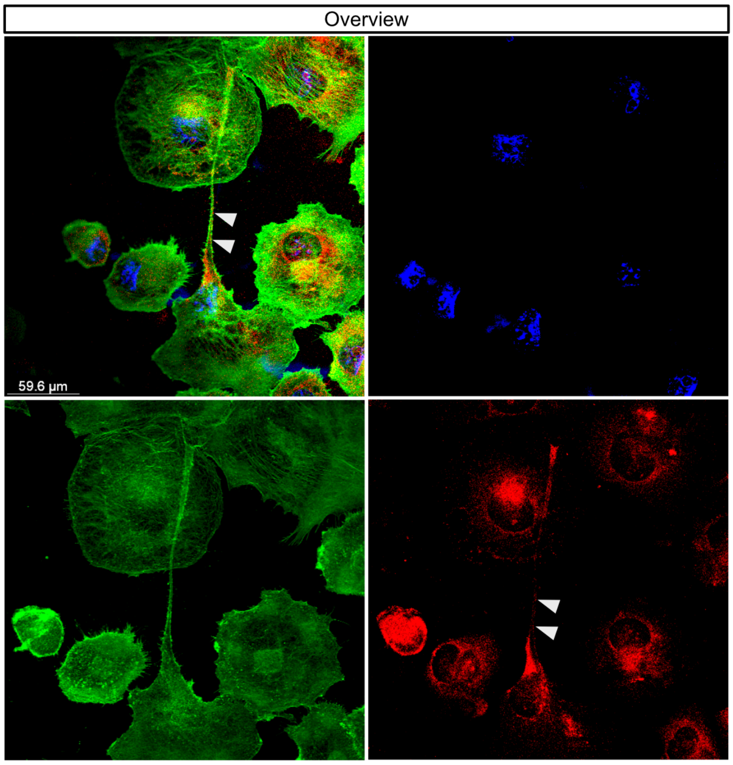


Overlay and channels (DNA – blue, actin – green, 5.8S rRNA antibody – red)

**c.** PANC-1 5.8S rRNA *
RNase treated negative control

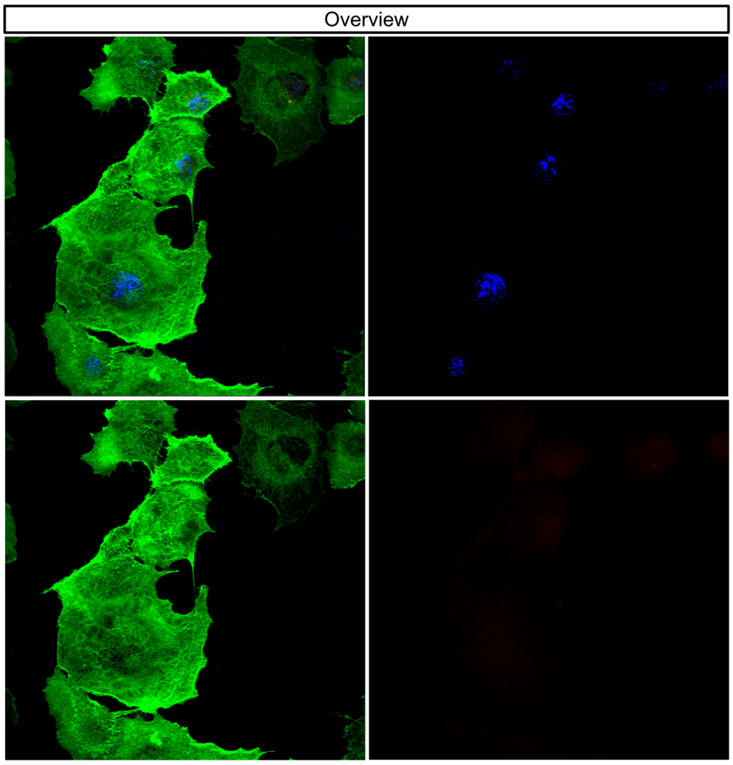


**d.** PANC-1 5.8S rRNA *
RNase treated negative control

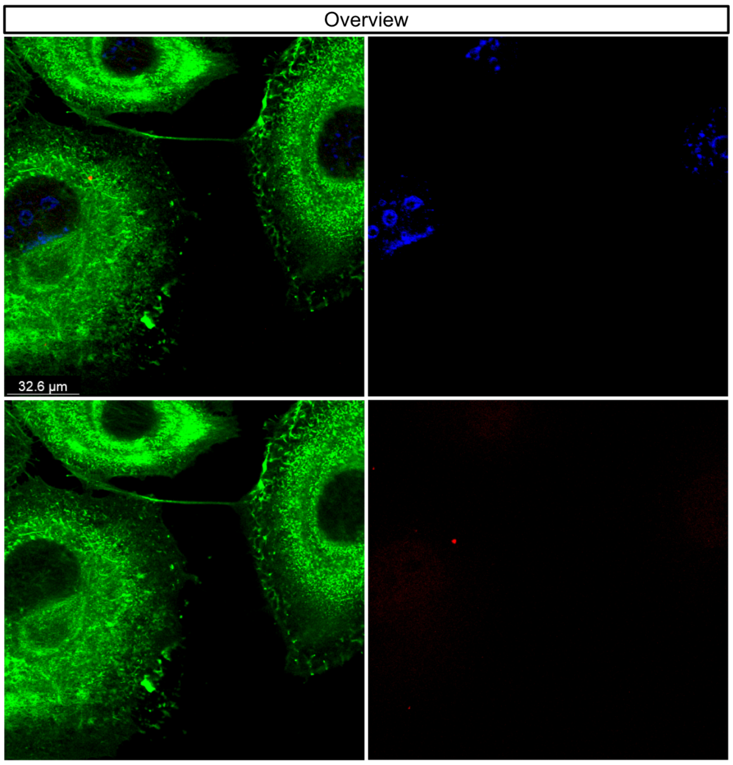


Overlay and channels (DNA – blue, actin – green, 5.8S rRNA antibody – red)

### FIGURE S7A cont.

**Figure S7A.** Ribosomal 5.8S rRNA detected in tunnelling nanotubes (a, b), RNase treated negative control stained for 5.8S rRNA (c, d) and no primary, secondary antibody control staining for 5.8S rRNA (e).

**e.** PANC-1 / no primary, secondary antibody control staining * for 5.8S rRNA

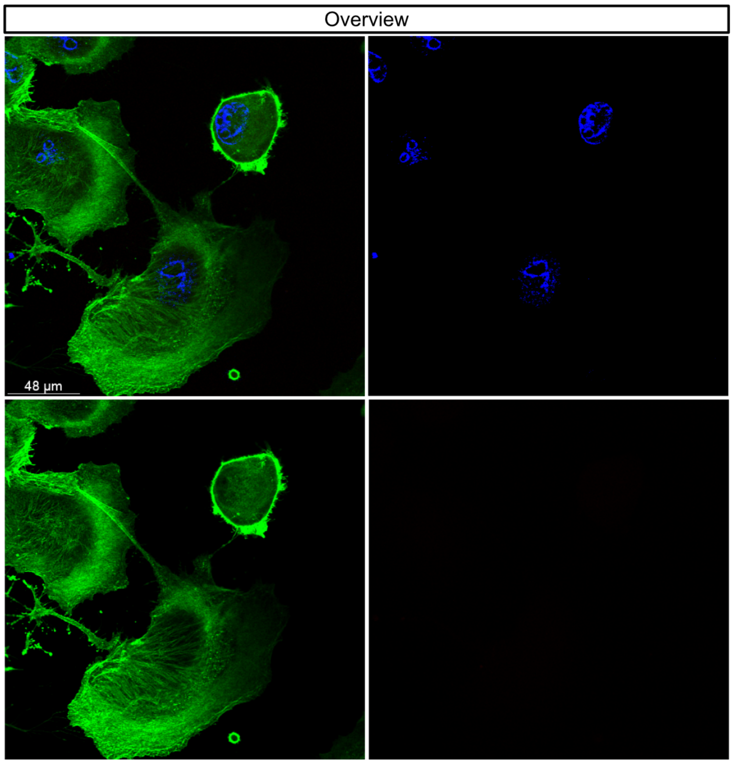


Overlay and channels (DNA – blue, actin – green, Secondary Antibody, Alexa Fluor™ 647– red)

### FIGURE S7D

**Figure S7D.** Polyadenylated RNA (mRNA) in tunnelling nanotubes (a), RNase treated negative control stained for mRNA (b) and no primary, secondary antibody control staining for mRNA (c).

**a.** PANC-1 mRNA * without treatment

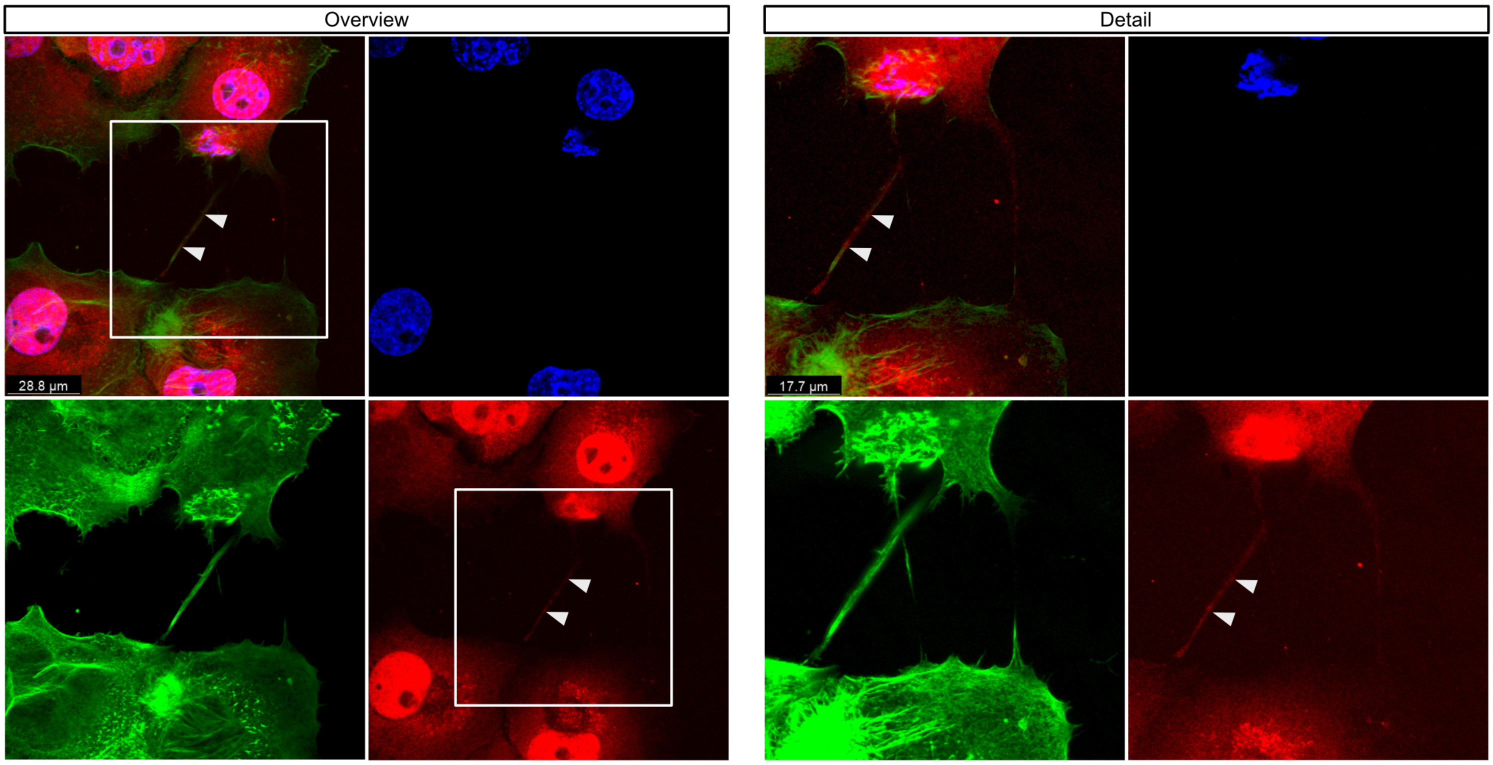


Overlay and channels (DNA – blue, actin – green, polyA probe – red)

**b.** PANC-1 mRNA * RNase treated negative control

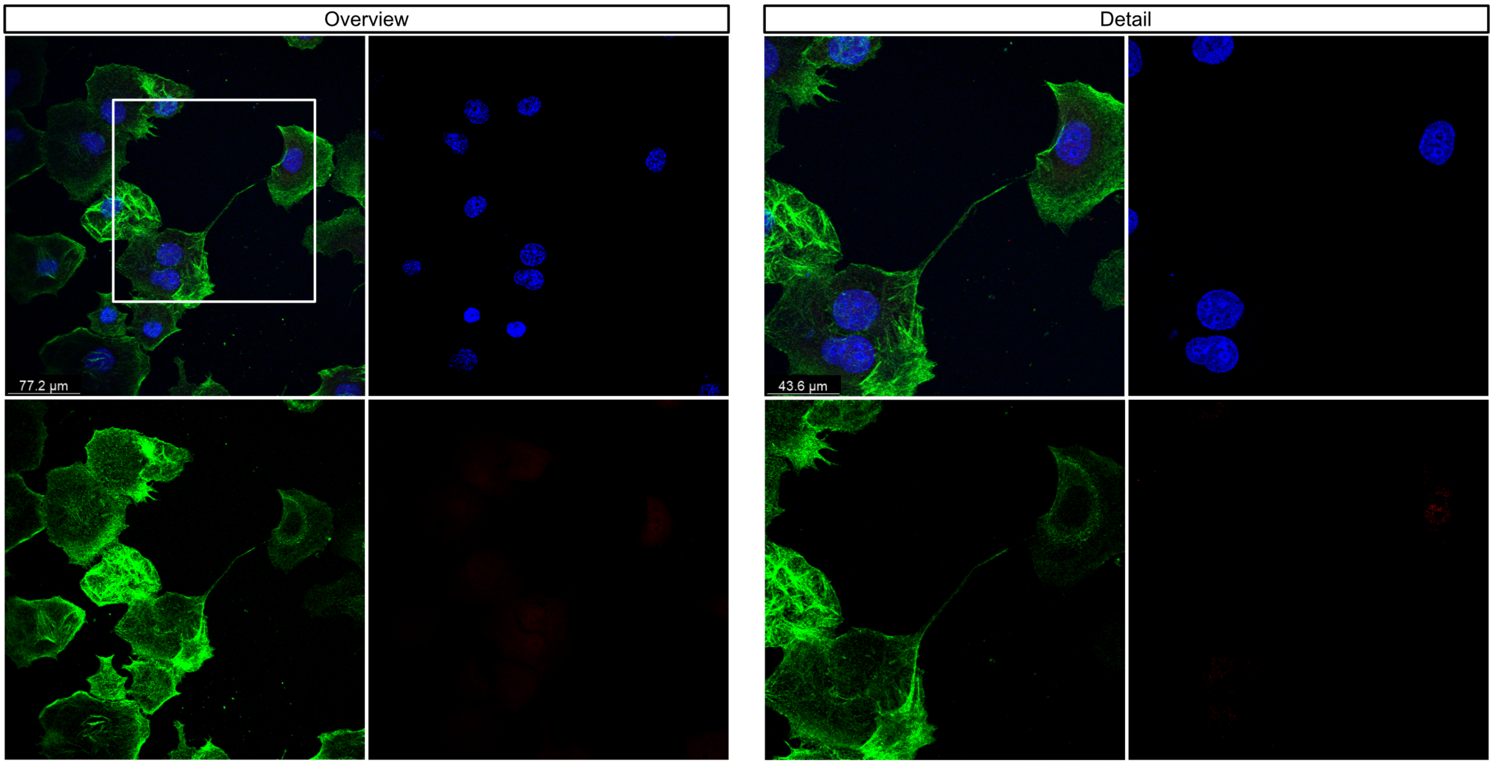


Overlay and channels (DNA – blue, actin – green, polyA probe – red)

### FIGURE S7D cont.

**Figure S7D.** Polyadenylated RNA (mRNA) in tunnelling nanotubes (a), RNase treated negative control stained for mRNA (b) and no primary, secondary antibody control staining for mRNA (c).

**c.** PANC-1 / no primary, secondary antibody control staining * for mRNA

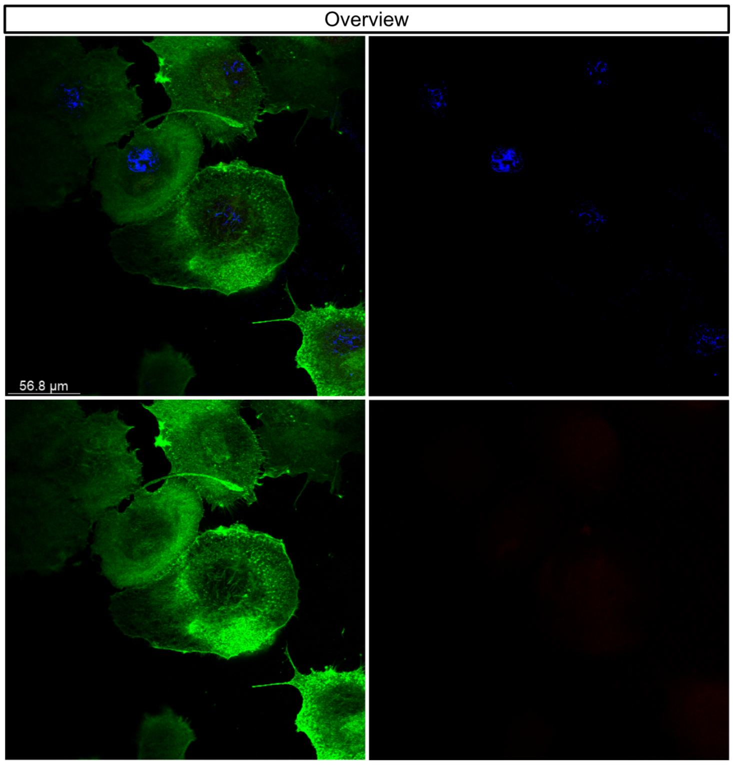


Overlay and channels (DNA – blue, actin – green, Secondary Antibody, Alexa Fluor™ 647– red)

### FIGURE S7E

**Figure S7E.** Complete ribosomes detected in tunnelling nanotubes (a, b) and puromycin treated negative control (c,d).

**a.** PANC-1 complete ribosomes (PLA) *

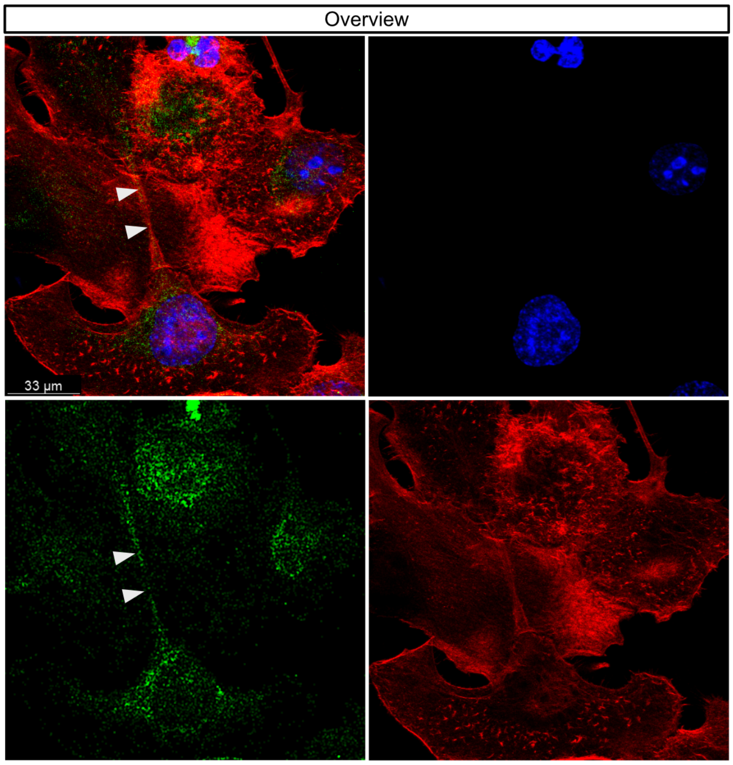


**b.** PANC-1 complete ribosome (PLA) *

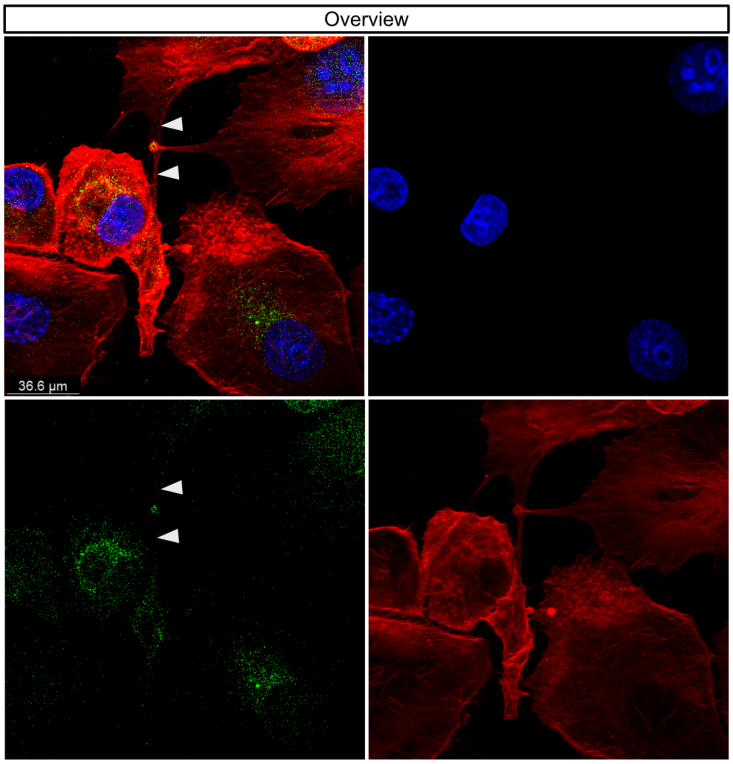


Overlay and channels (DNA – blue, RPS6+RPL24 proximity ligation – green, actin – red)

**c.** PANC-1 complete ribosomes (PLA) *
Puromycin treatment

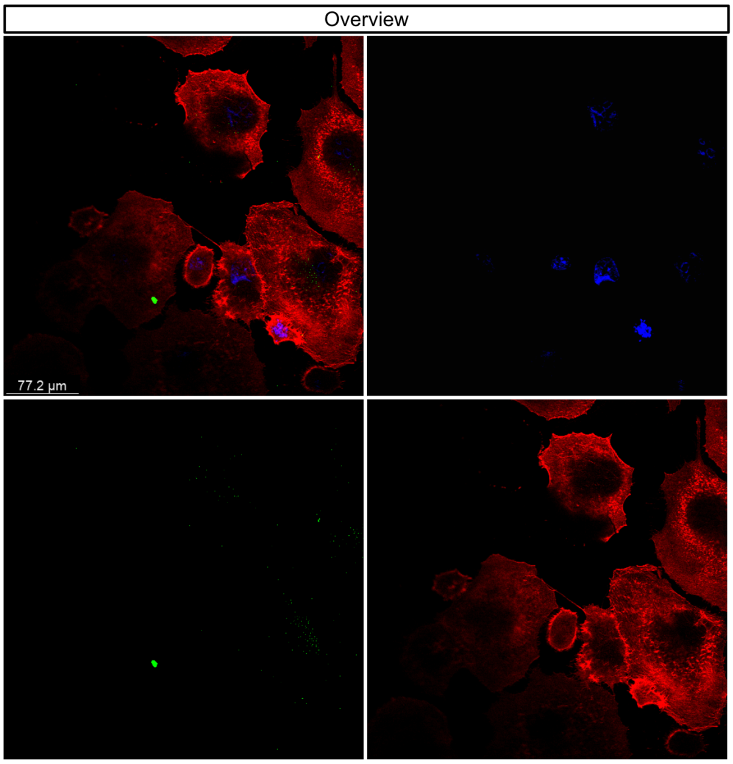


**d.** PANC-1 complete ribosome (PLA) *
Puromycin treatment

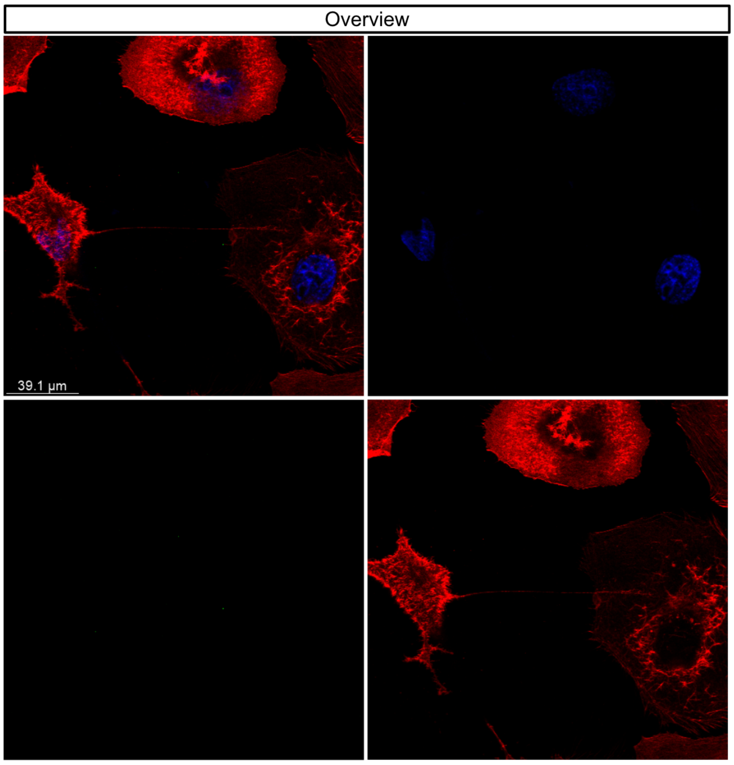


Overlay and channels (DNA – blue, RPS6+RPL24 proximity ligation – green, actin – red)

### FIGURE S8

**Figure S8.** Visualisation of pHTC-hsRPS9-HaloTag® CMV-neo vector created in SnapGene software.

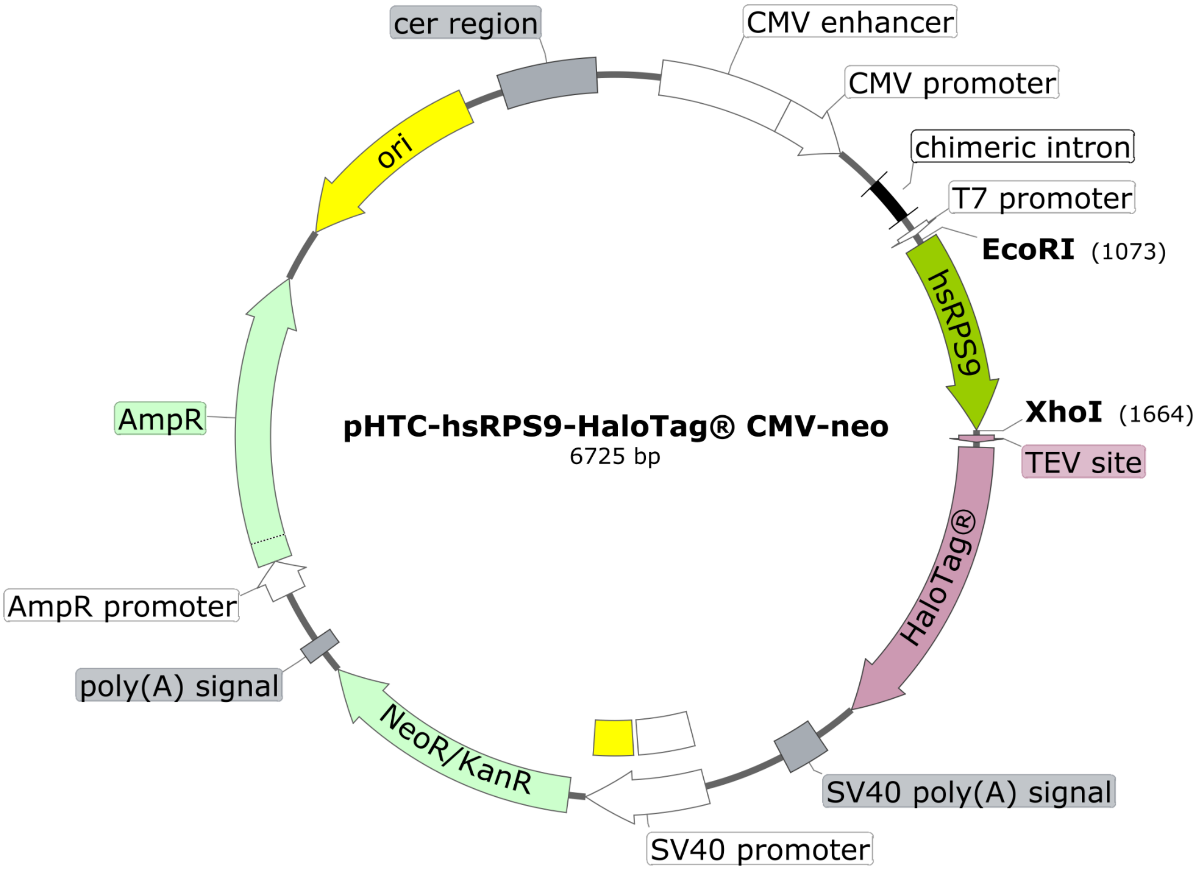

### FIGURE S9

**Figure S9.** The efficiency of siRNA transfection in PANC-1 cells was verified using Silencer™ Cy3-labelled negative control siRNA (a) #.

**a.** PANC-1 cells transfected with fluorescent siRNA

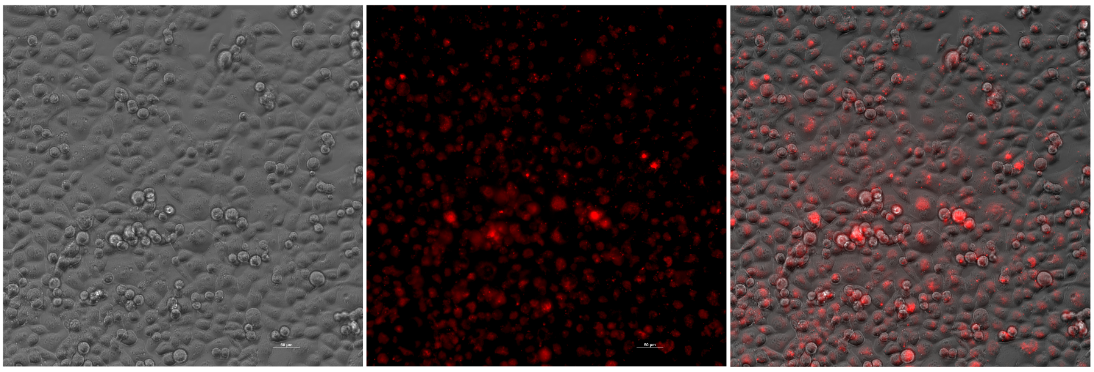

Overlay and channels (Bright-field, fluorescently labelled negative control siRNA – red)
